# Acute rapamycin treatment reveals novel mechanisms of behavioral, physiological, and functional dysfunction in a maternal inflammation mouse model of autism and sensory over-responsivity

**DOI:** 10.1101/2024.07.08.602602

**Authors:** JE Le Belle, M Condro, C Cepeda, KD Oikonomou, K Tessema, L Dudley, J Schoenfield, R Kawaguchi, D Geschwind, AJ Silva, Z Zhang, K Shokat, NG Harris, HI Kornblum

## Abstract

Maternal inflammatory response (MIR) during early gestation in mice induces a cascade of physiological and behavioral changes that have been associated with autism spectrum disorder (ASD). In a prior study and the current one, we find that mild MIR results in chronic systemic and brain inflammation, mTOR pathway activation, mild brain overgrowth followed by regionally specific volumetric changes, sensory processing dysregulation, and social and repetitive behavior abnormalities. Prior studies of rapamycin treatment in autism models have focused on chronic treatments that alter or prevent physical brain changes. Here, we have focused on the acute effects of rapamycin to uncover novel mechanisms of dysfunction related to mTOR pathway signaling. We find that within 2 hours, rapamycin treatment could rapidly rescue neuronal hyper-excitability, seizure susceptibility, functional network connectivity and brain community structure, repetitive behaviors, and sensory over-responsivity in adult offspring with persistent mild brain overgrowth. These CNS-mediated effects are also associated with alteration of the expression of several ASD-, ion channel-, and epilepsy-associated genes in the same time frame. Reduction of microglia with CSF1R inhibitors or inhibition of NADPH oxidase in young animals reduces the development of some of the behavioral phenotypes, but neither is as effective as acute mTOR inhibition. Our findings indicate that mTOR dysregulation in MIR offspring is a key contributor to various levels of brain dysfunction. However, we demonstrate that the adult MIR brain is also amenable to rapid normalization of these functional changes which results in the rescue of both core and comorbid ASD-like behaviors in adult animals without requiring long-term physical alterations to the brain. Restoring excitatory/inhibitory imbalance and sensory functional network modularity may therefore be important targets for therapeutically addressing both primary sensory and compensatory repetitive behavior phenotypes.

## Introduction

Neurodevelopmental disorders result from the disruption of brain development *in utero* or in early life^1^. Genetic, environmental, epigenetic, and immunological factors can all to contribute to their complex pathogenesis^1–5^. For example, there are numerous different anatomical abnormalities in brain structure and cellular changes that have been observed in the brains of autistic individuals and in animal models of autism that indicate a multifactorial pathophysiology^3,4,6–19^. Despite this complex etiology and the spectrum of phenotypes in Autism Spectrum Disorders (ASD), a common pathogenic mechanism has been identified for both genetic and non-genetic factors contributing to several neurodevelopmental disorders: dysregulated mTOR pathway signaling in the brain^6,7,20–31^.

Abnormal PI3K/Akt/mTOR signaling pathway activity in response to divergent underlying causes can lead to aberrant brain structure, synapse formation, synaptic pruning, excitatory/inhibitory balance, neuro-inflammatory signaling, and brain size^7,7,30,32–55^. Hyperactive mTOR signaling is known to contribute to seizures, sensory sensitivity, intellectual disability, macrocephalic brain growth, and a number of ASD-associated behavioral abnormalities^4,5,26,56–59^. This is thought to be related to the role of this important pathway in controlling cell growth, survival, energy balance, proliferation, autophagy, and ion channel expression. The mTOR pathway can become hyper-activated by several known mutations in ASD-associated genes such as PTEN, TSC1, TSC2, FMR1, Mecp2, DEPDC5, CACNA1C, UBE3A, and CNTNAP2^28,60–66^. ASD behaviors, macrocephaly, and epilepsy have also been recapitulated in a variety of PTEN and TSC murine knockout models in different cellular populations, developmental timeframes, and quantity of gene deletion. Furthermore, the Simons Foundation Autism Research Initiative (SFARI) has also identified a number of high-confidence autism risk genes in humans with several belonging to mTOR signaling gene sets^63,67–70^.

However, genetic mutations are not the only cause of increased mTOR activation and brain overgrowth. Hyper-activated mTOR signaling can both induce and be induced by neuro-inflammation of varying etiology^57,71–82^. There is considerable evidence of an elevated inflammatory state in ASD from gene expression studies, pro-inflammatory cytokines found in the blood and CSF and from increased activated microglia found throughout post-mortem autistic brains in both children and adults^19,83–97^. There is also evidence of brain overgrowth in ASD that is not linked to any specific genetic variation. Some studies have demonstrated that there is a limited period of increased, albeit not macrocephalic, brain growth early in development that has no known genetic or non-genetic etiology to date^98–100^. Maternal immune system activation is a good candidate mechanism for this phenomenon because it is also known to contribute to brain overgrowth and ASD in humans and in rodent models^19,85,101–117^. We have previously shown that exposure to a low-level maternal inflammatory response (MIR) induced by lipopolysaccharide early in gestation produces offspring with mild brain overgrowth, hyperactive mTOR signaling in the neurogenic SVZ niche, and abnormal ASD-related behaviors in mice^118^.

In addition to the core features of ASD (atypical social, communication, and stereotyped behaviors), there are many common comorbid phenotypes including sensory over-responsivity (SOR), epilepsy, disordered sleep, gastrointestinal disturbances, anxiety, depression, and irritability^119^. Rather than approaching treatment as a “cure” for ASD, a common approach for current pharmacological therapy is focused on treating these non-core symptoms with existing approved drugs. The rationale for treating each aberrant phenotype separately rather than tackling core symptomology is the complex and not fully understood etiology of ASD, the spectrum of ASD phenotypes and severity, and a lack of a core molecular target that underlies multiple aberrant behaviors. However, mTOR signaling is potentially one such convergent target that may impact both core and non-core phenotypes. Rapamycin, an inhibitor of mTOR, is known to be immunosuppressive, anti-proliferative, and an inducer of autophagy. It has been used in pre-clinical models of PTEN and TSC gene mutations to rescue the epileptic phenotype and several ASD-related behaviors in young mice by successfully preventing the development of aberrant brain physiological changes^9,120–124^. For example, developmental inhibition of mTOR over several weeks prevents brain overgrowth, loss of inhibitory neurons, seizures, and abnormal synaptic connectivity^9,79,125–130^. Rapamycin has, perhaps surprisingly, also been shown to rescue several abnormal phenotypes in adult mice despite the persistence of physiological brain abnormalities associated with PTEN and TSC gene mutations and mTOR activation, including macrocephalic brain overgrowth^9,19,124,125,131,132^. To date rapamycin analogs have been used clinically to treat patients with ASD and PTEN or TSC gene mutations for weeks to months with mixed results, improving some cognitive functions, reducing seizure frequency, and improving social behaviors in some studies but not in others^133–145^. Chronic rapamycin treatment has also been shown to rescue some phenotypes in idiopathic ASD mouse models with elevated mTOR signaling such as *in utero* valproic acid exposure, ASD-seizure, and BTBR models^122,125,146–149^.

While there has been encouraging progress at rescuing mTOR-mediated brain pathology and behavioral abnormalities in pre-clinical models with chronic rapamycin treatments, there could still be risks to normal brain development with juvenile treatment in humans without specific mTOR pathway-related gene mutations and adverse side effects from the use of such a potent growth inhibitor and immune suppressor. Therefore, we wanted to further understand the mechanisms that may underlie the effects of adult mTOR inhibition, where such treatment isn’t aimed at preventing or reversing structural brain abnormalities. Here, we focus on the effects of mTOR inhibition within an acute timeframe, revealing new mechanisms and functional changes that are different from those reported with a chronic inhibition treatment approach. We find that within two hours following rapamycin treatment of adult mice that had been exposed to mild maternal inflammation *in utero* results in significant normalization of deficits in behavior, sensory processing, brain functional connectivity and modularity, and electrophysiological hyperactivation. Furthermore, acute rapamycin affects specific cellular compartments and expression of ASD-, epilepsy and ion channel-associated genes. Thus, with these findings we have identified several potentially less disruptive targets like systemic inflammation, sensory brain circuitry, functional network modularity, and neuronal hyper-excitability, for pharmacological or neuromodulatory intervention in several ASD physiological and behavioral phenotypes in the CNS.

## Materials and Methods

The following methods have been communicated using the ARRIVE 2.0 guidelines^150^.

### Study Design

The primary objective of this study was to evaluate the low-dose, early gestation model of maternal inflammation on offspring studied chronically from young adult (P60-90) to old adult (P200-400), to assess the effects of acute rapamycin treatment at these ages, and to identify mechanisms contributing to its rapid effects on behavior. Our main hypotheses were that the MIR model replicates multiple phenotypes relevant to autism (brain overgrowth, chronic systemic inflammation, microglia contributions to dysfunction, sensory dysregulation, autism-associated gene expression changes, etc.), and that the rapid effects of rapamycin treatment could identify novel mechanisms of dysfunction that are different from published chronic effects of rapamycin treatment involving physical synaptic remodeling.

### Experimental Framework

A parallel group design was used for most experiments where there are separate MIR and Control offspring groups for every treatment and for different ages (young adult and old adult). The fMRI experiments used a within-subjects crossover design where mice were first tested at baseline and then again after rapamycin treatment. The experimental unit used for all studies are individual mouse offspring. Multiple litters of maternal inflammation-exposed offspring and vehicle control-exposed offspring were generated and equal numbers of male and female offspring were randomly selected from all litters to generate each experimental group. Some mice were used as young adults and other animals from the same litters were kept for experiments on older adult mice. These experiments were performed with 52 total litters generated over several years of experimentation (see supplemental table 2). Every new litter generated was evaluated for repetitive behaviors in an open field to confirm that the MIR phenotype was reproduced in the new offspring before using them for experiments, including non-behavior experiments. No MIR litters failed to display a significant repetitive behavior phenotype in offspring, demonstrating the reproducibility of the model. Because batch variations in LPS strength/effectiveness have been reported we used the same batch of LPS, stored in small aliquots at -80 degrees C, and made up fresh in saline for injection for every MIR litter generated. See Fig. 1A for an outline of this experimental timeframe.

**Figure 1.**
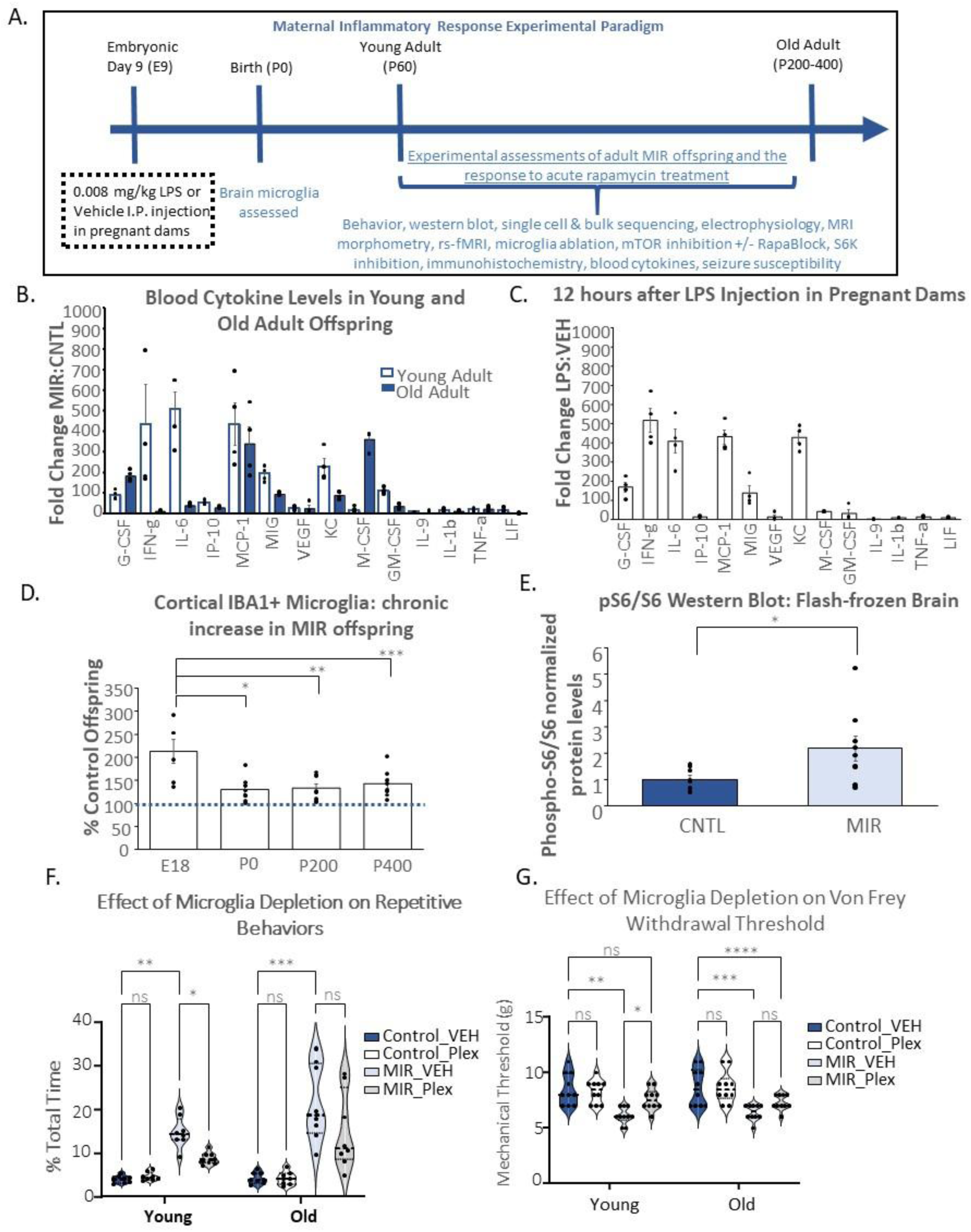
MIR offspring have a chronically increased inflammatory state and mTOR pathway activation and abnormal repetitive and social behaviors and sensory sensitivity in adult mice following a single, low-dose LPS injection early in gestation that is reduced by microglia ablation in young adult but not old adult offspring. (**A**) Experimental timeline summary; (**B**) Inflammatory blood cytokines are significantly increased in both young and old adult MIR offspring relative to age matched controls (Two-way ANOVA with multiple comparisons cytokine x group effect F_(12, 44)_, p<0.0001 and group effect F_(5, 18)_, p<0.0001) and are expressed as fold-change in MIR relative to controls. Post-hoc differences (Tukey’s) for G-CSF=granulocyte colony stimulating factor (young control v MIR, adjusted p=0.0024, old control v MIR, adjusted p=0.0048), IFN-g=interferon gamma (young control v MIR, adjusted p=0.0137, old control v MIR, adjusted p=0.0001), IL-6=interleukin 6 (young control v MIR, adjusted p=0.0064, old control v MIR, adjusted p=0.0015), IP-l0=interferon gamma induced protein factor(young control v MIR, not significant p=0.1132, old control v MIR, adjusted p=0.0036), MCP-l=monocyte chemoattractant factor (young control v MIR, adjusted p<0.0001, old control v MIR, adjusted p=0.0341), MIG= monokine induced by gamma protein interferon factor(young control v MIR, not significant, p=0.2705, old control v MIR, adjusted p=0.0419), VEGF=vascular endothelial growth factor(young control v MIR, adjusted p=0.0085, old control v MIR, adjusted p<0.0001), KC=keratinocyte chemoattractant factor(young control v MIR, adjusted p=0.0184, old control v MIR, adjusted p=0.0027), M-CSF=macrophage colony stimulating factor (young control v MIR, adjusted p<0.0001, old control v MIR, adjusted p=0.0006), GM-CSF=granulocyte macrophage colony stimulating factor(young control v MIR, adjusted p=0.0044, old control v MIR, adjusted not significant, p=0.0505), IL-9=interleukin 9 factor (young control v MIR, adjusted p=0.0107, old control v MIR, adjusted p=0.0009), IL-lb=interleukin 1 beta factor (young control v MIR, adjusted p=0.0003, old control v MIR, adjusted p=0.0062), TNF-a=tumor necrosis factor alpha factor (young control v MIR, adjusted p=0.0006, old control v MIR, adjusted p=0.0036), LIF=leukemia inhibitory factor (young control v MIR, adjusted p=0.0009, old control v MIR, adjusted not significant, p=0.7465), n= 4/group; data shown as mean+/- SEM; (**C**) Blood cytokine changes in LPS injected pregnant dams (0.008 mg/kg) graphed as fold-change relative to vehicle-injected control pregnant dams at 12 hours post-injection, (Two-way ANOVA cytokine x group effect F_(12, 44)_, p<0.0001 and group effect F_(5, 18),_, p<0.0001) and are expressed as fold-change in LPS-injected relative to vehicle controls. G-CSF (control v LPS, adjusted p<0.0001), IFN-g (control v LPS, adjusted p=0.0001), IL-6 (control v LPS, adjusted p<0.0012), IP-10 (control v LPS, adjusted p<0.0001), MCP-1 (control v LPS, adjusted not significant, p=0.2705), MIG (control v LPS, adjusted p=0.0016), VEGF (control v LPS, adjusted p=0.0021), KC (control v LPS, adjusted p=0.0197), M-CSF (control v LPS, adjusted p=0.0011), GM-CSF (control v LPS, adjusted p<0.0001), IL-9 (control v LPS, adjusted p=0.0267), IL-lb (control v LPS, adjusted p=0.0022), TNF-a (control v LPS, adjusted p=0.0002), LIF(control v LPS, adjusted p=0.0140)), N=4/group; Data shown as mean+/- SEM; (**D**) Stereoinvestigator quantification of IBA+ immunohistochemistry staining of microglia in the sensory-motor cortex of adult MIR offspring, graphed as % control, One-way ANOVA with multiple comparisons p<0.0001, F_(3, 26)_ with significant (Tukey’s) differences between E18 v PO (*p<0.00001), E18 v P200 (**p<0.0001), E18 v P400 (***p<0.0001) and non-significant differences for PO v P200 (p=0.9975), PO v P400 (p=0.8248) and P200 v P400 (p=0.9068), N= 8/group; Data shown as mean+/- SEM (**E**) Phospho-S6 protein measured by western blot on microdissected, flash-frozen brain tissue from cortex, amygdala, and striatum (combined), Paired t-test (two-tailed) showed significant difference between control and MIR offspring, *p=0.0220, df=8, n=3/group; data shown as mean+/- SEM; (**F**) 2-way ANOVA with multiple comparisons shows significant overall group (p<0.0001, F_(3,28)_ and age (p=0.0140, F_(1, 28)_) effects in repetitive behaviors that are significantly increased in post-hoc (Tukey’s) tests for control v MIR offspring in young (**p<0.0001) and old (***p=0.0001) mice in the control diet groups and significantly decreased in MIR-vehicle (VEH) diet vs MIR-Plex diet in young (*p=0.0301) but not old (p=0.0590) mice after 3 week Plexxikon 5622 (Plex) treatment to ablate brain microglia, N=8/group (**G**) Two-way ANOVA with multiple comparisons showed a significant overall group effect (p<0.0001, F_(3, 36)_) in the limb withdrawal threshold of the hind paw in the Von Frey test in control and MIR offspring after 3 weeks of Plexxikon 5622(Plex) or vehicle control (VEH) diets, post-hoc (Tukey’s) analysis shows a significantly lower withdrawal threshold for MIRmice compared to control offspring in young (**p<0.0001) and old (***p=0.0002) mice in the control diet groups and significantly increased withdrawal threshold in MIR-vehicle (VEH) diet vs MIR-Plex diet in young (*p=0.0232) but not old (p=0.3031) mice after 3 week Plexxikon 5622 (Plex) treatment to ablate brain microglia, there is also no significant difference (ns) in control offspring on the vehicle diet compared to the MIR offspring on the Plex diet, demonstrating the rescue effect in the young mice (p=0.1810) but this difference remains significant in the old mice, demonstrating a reduced effect with age (****p=0.0002), n=l0/group.

### Ethical Approval

All experimental procedures were designed to minimize animal suffering and were conducted in strict accordance with the Public Health Service Policy on Humane Care and Use of Laboratory Animals. The protocols were prospectively reviewed and approved by the University of California, Los Angeles (UCLA) Chancellor’s Animal Research Committee (ARC). This manuscript is reported in accordance with the ARRIVE 2.0 guidelines.

### Sample Size

Sample sizes were determined based on our own previous experiments and relevant experiments published in the literature to achieve a statistical power of 0.8 with an alpha of 0.05 to detect an effect size (Cohen’s d) of 1.2. The significance and the effect size were reported for every result in the figure legends and results section. Group numbers has been reported in all figure legends and individual data points are plotted in result figures.

### Inclusion and Exclusion Criteria

Mice from multiple litters were randomly assigned to each group and no selection criteria were used. An equal number of male and female mice were used in each group. Mice injured due to fighting in home cages, which happened occasionally, and were excluded from study due to possible confounding effects of that would be brought about by single housing and medical treatments. No animal exclusions were necessary due to a failure to perform the behavioral tests. Because we used an outbred strain of mice (CD1) that have some genetic variability, a range of phenotype severities was expected. Therefore, outlier data and outlier animals were not removed from any experiment since these mice likely represent the possible range of outcomes in the model.

### Blinding

Animal group identification was not written on cage cards so that researchers were blinded to experimental groups during data collection. Group identification was known to the investigator analyzing the data only after all data had been collected. Treatment group but not litter group was known when rapamycin or vehicle treatment was administered. However, due to the development of extreme repetitive behaviors in a third of the MIR offspring at older ages it was not possible to be fully blinded to the group for the old-adult mice. MIR mice at older ages also frequently weighed less due to constant repetitive activity compared to control mice which normally become more sedentary and increase in weight significantly with age. MIR mice at young adult ages were not readily identifiable by casual inspection of home cage behavior.

### Experimental Animals

Time-mated, pregnant CD-1 wild-type mice were obtained from Charles River Laboratories (CA, USA). On day E9 of gestation, pregnant dams received a single intraperitoneal (I.P.) injection of low-dose lipopolysaccharide (LPS, E. coli serotype 0111:B4; Sigma) at 0.008 mg/kg dissolved in sterile saline, or an equivalent volume of sterile saline (Vehicle Control). This low dose was chosen as it induces a cytokine response without causing overt sickness behavior in the pregnant dam and mild brain overgrowth in the offspring (15-20% greater than vehicle control offspring but not macrocephalic) as we have previous published^118^. The same batch of LPS was used for the generation of all MIR litters in this manuscript. A different batch of LPS used at the same dosage generated MIR offspring in our previous publication of this model, demonstrating its reproducibility. Litter sizes ranged from 9-16 offspring and mice were weaned at post-natal day 21. After weaning, mice from the same litter were socially housed together 2-4 per cage by sex. Mice were randomly selected from multiple litters to form experimental groups and equal numbers of male and female mice were used in each group for all experiments. Separate cohorts of mice were used as young adult ages (P60-90) and old adult ages (P200-400) for specific experiments. Mouse age group have been reported in figures and figure legends. The CD-1 strain of mice was used because of the similarity that their stem and progenitor brain cells have to human brain Hif1-alpha redox responses that are relevant to an inflammation model, their large litter sizes, and their consistent responses in behavioral testing as seen in our original publication of this MIR model^118^ (Le Belle, et al., 2014). **Husbandry Details:** Animals were socially housed under standard pathogen-free conditions on a 12-hour light/dark cycle at a constant temperature (20-26°C) and humidity (50%). They had ad libitum access to standard rodent chow and sterile water. Cages contained aspen chip bedding and were provided with nesting material for enrichment. At the end of all experiments, mice were euthanized by CO2 and rapid decapitation or by perfusion-fixation following pentobarbitol overdose and succession of respiration. No unexpected animal deaths occurred and the only adverse events that occurred were due to a small number of cases of animal fighting in the home cages outside of experimental work.

### Pharmacological Treatments

All pharmacological treatments were administered to the mice as young adult (P60-90) or old adults (P200-400), apart from the juvenile mice (P25) in the reciprocal social interaction test (see supplemental methods and Fig. S1B). Juvenile mice are commonly used for this test because they are less territorial and more likely to engage in reciprocal behaviors without aggression or dominance behaviors.

### Rapamycin

Rapamycin (5 mg/kg; Sigma-Aldrich) or vehicle (DMSO) was administered via intraperitoneal (I.P.) injection in the flank. For all acute treatment experiments in young and old MIR and control offspring, this was given 2 hours prior to data collection. The rapamycin dose was chosen based on previous published work that showed behavioral rescue in genetic murine models of autism in adult offspring^151^. For the chronic treatment experiment, daily I.P. injections (5 mg/kg) were given for 5 weeks.

### RapaBlock

To differentiate central vs. peripheral effects, young adult mice were co-administered rapamycin and the brain-impermeable inhibitor RapaBlock. RapaBlock is a small molecule^152,153^ that binds FKBP12, and acts to prevent rapamycin from accessing this necessary factor for mTOR inhibition in the peripheral nervous system. Therefore, mice co-administered both rapamycin and RapaBlock would still have mTOR inhibition in the central but not in the peripheral nervous system. RapaBlock was dissolved in DMSO and dissolved in DPBS containing Tween80 and PEG300. Mice were injected simultaneously with 5 mg/kg dose of rapamycin or vehicle control (DMSO) and with 40 mg/kg of RapaBlock or vehicle control, given at the same time via intraperitoneal injection in opposite flanks. Animal behavior was measured in control and MIR offspring groups in treatment conditions with and without rapamycin and with and without RapaBlock. The comparison of the treatment effects of Rapamycin given alone compared to Rapamycin co-administered with RapaBlock indicated whether any rescue effects are due to central nervous system effects of Rapamycin.

### S6K1 Inhibitor

The S6K1 inhibitor PF-4708671 (75 mg/kg; formulated in 10% DMSO, 10% Tween-80, 80% water) as described in Huang, et al^154^., or vehicle was given I.P. 2 hours or 6 hours prior to data collection in young adult MIR and Control offspring.

### NOX Inhibitor

The NADPH oxidase (NOX) inhibitor Apocynin (Sigma-Aldrich, cat no. 178385) or vehicle was given I.P. 2 hours prior to data collection or daily for 5 weeks for chronic treatment in young adult MIR and Control offspring.

### Microglia Depletion

Young and old adult mice were fed a diet containing the CSF1R-inhibitor Plexxikon 5622 (Plexxikon-Daiichi, San Francisco) formulated at 1200 mg/kg in rodent chow (AIN-76A; Research Diets, New Brunswick, NJ) or control AIN-76A chow ad libitum for 3 weeks prior to and during behavioral testing. Microglia depletion was confirmed (>98%) with post-mortem immunohistochemistry with the myeloid cell marker, IBA1 (Wako 019-19741).

### PTZ treatment

Mice were tested for seizure susceptibility using an escalating pentylenetetrazole (PTZ, Millipore P6500) dosing protocol (0-40 mg/kg, I.P.) modified from Shimada and Yamata^155^. Following intraperitoneal injection with PZT, a GABAA receptor antagonist, animals were monitored continuously for 30 minutes. High doses of PTZ are known to induce an acute, severe seizure, but sequential injections of a sub-convulsive dose are used for the development of chemical kindling, an epilepsy model. With this method, vulnerability to PTZ-mediated seizures can be estimated. PTZ was dissolved at 2 mg/mL in sterile 0.9% (w/v) NaCl and prepared on the day of use. Mice were injected i.p. with a starting dose of 10 mg/kg and then injected again every 2 days, increasing the dose by 10mg/kg. Mice were monitored for the following seizure-related escalating behaviors for 30 minutes following the injection: (0) no seizure behaviors, (1) immobilization, (2) head nodding and partial myoclonus (sudden, brief involuntary twitching or jerking of a muscle or group of muscles), (3) continuous whole body myoclonus, (4) rearing or tonic seizure (falling on the side followed by forelimb tonic contraction and hindlimb tonic extension), (5) tonic-clonic seizure with rushing and jumping. Both control and MIR mice were given escalating doses of PTZ until all mice from one group were displaying seizure-like activity anywhere on the 1-5 scale and the number and severity of seizure symptoms at this maximum PTZ dose (40mg/kg) were then compared between groups to determine relative induced seizure susceptibility. The effects on seizure-behavior level at the maximum PTZ dose after acute rapamycin treatment in MIR and vehicle-control offspring was also compared. Humane endpoint: the experiment was to be terminated for an animal if it experienced a tonic-clonic seizure lasting longer than 60 seconds or failed to recover normal posture within 5 minutes bur no animals met these criteria. The primary outcome of the PTZ experiment was identifying the number of mice which reached each seizure-behavior level (1-6) with and without rapamycin treatment at the maximum PTZ dose (40mg/kg). A quantitative seizure score was generated for each mouse based on what seizure level (1-6) the mouse was at for each dosage of PTZ and PTZ + rapamycin for statistical analysis in supplemental results (Table S3).

### Blood Cytokine Analysis

Blood was collected from pregnant dams on E9 at 12-hours after LPS injection (0.008 mg/kg, I.P., Fig. 1C) and from MIR and control offspring as adults. Blood plasma was separated and frozen before analysis by the UCLA Pathology Immune Assessment Core using the Luminex’s xMAP® Mouse 32-plex Cytokine/Chemokine Panel Immunoassay according to standard assay methods. The primary outcome was the blood cytokine levels resulting from acute LPS injection in pregnant dams and in adult MIR-exposed and vehicle-control-exposed offspring.

### Regional Brain Microdissection and Western Blot

The sensory-motor cortex, amygdala, and striatum were microdissected from adult MIR and control offspring and from a separate cohort of adult offspring after acute rapamycin treatment and flash frozen for standard western blot analysis. Tissue homogenates (cytosol) were lysed in RIPA buffer (25 mM Tris-HCl, 150 mM NaCl, 1% NP-40, 1% sodium deoxycholate, 0.1% SDS, pH 7.6 at 4℃) containing a cocktail of protease inhibitors (Calbiochem). Equal amounts of protein were separated by SDS-PAGE (4–12% Bis-Tris gels, Invitrogen) and transferred to PVDF membranes (Invitrogen), blocked in TBST plus 5% non-fat milk and then incubated with the following primary antibodies: phospho-S6 (S235/236; 1:2000, Cell Signaling, #4858), S6 (1:2000, cell signaling, #2217), and b-actin (1:2000, Cell Signaling, #4970) overnight at 4℃. The membranes were then washed with TBST and incubated for 1 h at room temperature. The washed membranes were then treated with an enhanced chemiluminescence detection reagent (Thermo Scientific). All Blots were developed using ChemiDoc XRS+ Molecular Imager (BioRad) and analyzed using Quantity One software (BioRad). Band densities were normalized to the total amount of protein loaded per lane using Sypro Ruby (BioRad). The 3 regions of brain tissue were pooled from MIR and vehicle-control offspring for comparison of phospho-S6 protein levels normalized to total S6 protein and GAPDH protein levels in mice treated with a 2-hour rapamycin or vehicle treatment.

#### Behavioral Testing

Mice were housed 4 per cage on a 12-hour light/dark cycle with ad libitum access to food and water. All behavioral experiments were performed during the light phase (10:00-16:00) by a blinded experimenter. Animals were habituated to handling for 3 days prior to experiments and habituated to the testing room for 60 minutes the day before and the day of any test. Mice which took part in the open field, light/dark, and Von Frey testing were habituated to the testing apparatus in a 10-minute habituation session on two consecutive days before testing. A battery of tests was performed on the same cohort of mice in order of least to most stressful, with at least 48 hours between tests: open field, tactile light/dark box, Von Frey, and finally pre-pulse inhibition. A separate cohort of young mice (P25) were used for the juvenile reciprocal social interaction testing. Behavioral equipment was cleaned between animals with 70% ethanol followed by tap water and then dried to remove residual odor cues.

### Open Field Repetitive Behaviors

Measurement of the cumulative time spent performing repetitive behaviors in an open field chamber was performed during a 10-minute session according to the methods of Peñagarikano et al^156^. The total amount of time that the mice spent performing repetitive behaviors (grooming while stationary or while moving on all body regions and repetitive circling seen in some of the older mice) was measured. The primary outcome measure was the percentage of total testing time spent performing repetitive behaviors of any kind. A square open-field arena (40 × 40 x 40 cm) constructed of opaque gray PVC was used. The floor was non-textured and non-absorbent and no bedding was present. Illumination at the arena was set to 50 lux during testing.

### Von Frey SODU

The Von Frey Simplified Up-Down (SODU) method from Bonin^157^ was used for assessing mechanical (tactile) sensitivity on the hind paws. We used standard Von Frey filaments (Harvard Apparatus), which are thin nylon fibers of varying diameters which are used to apply a controlled mechanical force onto the mouse’s paw to assess the force threshold at which the animal senses (notices) the touch and lifts/withdraws its paw. The paw withdrawal threshold (PWT) is a quantified measure of fiber force strength at which this behavioral response is observed. A wire mesh platform was used as the testing area. Mice were habituated to this platform while in a clear plastic holding chamber for 1 hour prior to testing. During the test the mice were placed on top of the wire mesh platform and allowed to freely explore. A low threshold Von Frey fiber was used to touch the plantar surface of one of the hind paws of the mouse through the mesh frame for 1-2 seconds. The response of the mouse was recorded (either lifting its paw or no response). If the mouse withdrew its paw then the next lower filament was used for the second touch. If the mouse did not respond then the next higher filament was used for the second touch. This process was repeated for 5 touches in total. Outcome Variables: We used the final response to the fifth filament to estimate the PWT. If the mouse did not withdraw its paw on the fifth touch, 0.5 was added to the logarithmic value of the fifth filament’s diameter. If the mouse withdrew its paw on the fifth touch, 0.5 was subtracted from the logarithmic value of the fifth filament’s diameter. The resulting logarithmic value was converted to a linear scale to obtain the PWT in grams. The primary outcome measure was the mechanical threshold in grams at which the mice felt/responded to the fiber pressure on the bottom of their hind paw on the 5th touch.

### Light/Dark Box Tactile Avoidance Testing

The tactile avoidance light/dark box is a novel method developed to examine tactile sensitivity of the paws and avoidance behaviors. This test used the classic light/dark chamber box which is traditionally a test for anxiety in mice where normal mice will explore both light and dark chambers but mice prefer the dark chamber and mice with heightened anxiety will spend much more time in the “safe” dark chamber compared to normal controls^158^. We adapted this test to use either a smooth plastic floor or a rough textured floor in one chamber using rough, 80-grit anti-slip stair adhesive strips. The time mice spend in the left or right chamber when both are lit and the floor is smooth plastic (trial 1), when both are lit but the floor in one chamber has the rough texture (trial 2), when one chamber is dark but both floors are smooth (trial 3--the classic light/dark box test), and finally when the dark (preferred) chamber had the rough textured floor and the lit chamber had a smooth floor (trial 4) was measured. Each trial was 10-minutes long and was performed 24 hours apart with the order of the trials randomized for each group. The primary outcome measures were the time spent in each chamber for each trial. Spending less time in the normally preferred dark chamber when the floor was a rough texture but not when the floor was smooth demonstrated an aversive tactile sensitivity and avoidance behavior. Each chamber was a square arena (40 × 40 x 40 cm) constructed of black acrylic. Illumination at the arena floor was set to 150 lux during all trials.

### Pre-Pulse Inhibition and Habituation to Sensory Startle

The San Diego Instruments SR-Lab Startle Response System was used to test for sensory gating ability in the different animal groups. The system software is programed to produce a randomized combination of a low-intensity airpuff onto the fur on the back of the mouse as the pre-pulse sensory “warning” followed by a loud (20 dB, startle-inducing) sound at an increasing distance in time (50-1000 ms) between the pre-pulse airpuff and the audible startle pulse according to the methods of Orefice, et al^159^. Startle-only pulses were also included at the beginning, middle, and end of the full test. The amplitude of the startle response by the mouse was measured by a sensitive gyroscope within the sound-proof testing chamber. Mice were placed within the startle enclosure and testing time was approximately 30 minutes per session. When sensory gating is normal, the closer in time that the tactile pre-pulse warning (light airpuff) is given to the auditory startle sound, the greater the inhibition (decrease) there is in startle response from the mouse. Primary outcome measures: The SR-LAB software automatically records the mouse’s startle response (i.e., movement detected on the platform). This data was then used to calculate the Pre-Pulse Inhibition percentage by comparing the startle response amplitude in the presence of the changing time distance of the prepulse stimulus. In addition, a startle only (without prepulse warning) at the beginning, and end of the PPI program was used to test for habituation to startle over time and was shown as a ratio (amplitude of end startle: amplitude of beginning startle) where values >1 indicate a lack of habituation.

#### Single Cell and Bulk Genetic Sequencing and Analysis

Mice were euthanized according to approved protocols and sensory-motor cortex tissue was microdissected from MIR and control offspring and flash frozen before being prepared for sequencing analysis. For bulk RNA sequencing, 12 MIR and 12 control mice were given rapamycin or vehicle 2 hours before euthanizing and flash freezing the brain tissue for sequencing. RNA was isolated using a Qiagen RNeasy Mini kit (Cat. 74106) followed by library preparation using a KAPA Stranded mRNA-Seq Kit (Cat. KR0960). The sequencing libraries were pooled and sequenced to generate minimum of 220 M 50 paired-end reads on NovaSeq6000 (Illumina). Reads were aligned to mouse mm10 genome reference using STAR aligner (v2.4.0). Read counts for refSeq genes were generated by HT-seq 0.6.1. Various QC were performed after alignment to examine the level of mismatch rate, mapping rate to the whole genome, repeats, chromosomes, key transcriptomic regions (exons, introns, UTRs, genes), insert sizes, AT/GC dropout, transcript coverage and GC bias. Outliers were removed based on QC results. Differential expression analysis was conducted with R-project and the Bioconductor package EdgeR. Differentially expressed genes (DEGs) between groups were defined as those meeting all of the following criteria: false discovery rate (FDR) < 0.05, LogFC > 0.5, and p < 0.05 threshold (see supplemental data for complete gene list). We also tested for overlap between these DEGs and high-confidence autism-associated genes in ASD databases^68,96^.

For single-nucleus RNA sequencing, 6 MIR and 6 control mice were analyzed. Single nuclei were isolated from frozen brain tissue using iodixanol-based density gradient centrifugation and submitted to UCLA Technology Center for Genomics and Bioinformatics for library preparation (via Chromium Single Cell 3’ v3 kit from 10x Genomics) and sequencing (via NovaSeq 6000 S2 platform from Illumina). Cell Ranger (10x Genomics) was used for alignment, filtering, barcode identification, and generation of gene-cell matrices for each sample. Ambient RNA was removed using CellBender^160^. Pre-processing (minimum cells 3, minimum features 200, doublet removal via scDblFinder^161^, maximum mitochondrial content 5%), integration (3000 features), and clustering (resolution 0.8) were performed using the R package Seurat^162–164^. Canonical lineage markers were used to identify cell types (microglia, astrocytes, oligodendrocytes, pericytes, endothelial cells, upper layer and deep layer excitatory neurons, and interneurons). Differential abundance of cell types in MIR vs CTRL was assessed using the R package propeller^165^. Differential expression testing was performed using the pseudobulk approach (edgeR with likelihood ratio test) via the R package Libra^166^. Gene set enrichment analysis (GSEA)^167^ was performed to assess differential enrichment of the “Hallmark mTORC1 Signaling” gene set^168,169^ using the R package fgsea^170^. The above comparisons underwent correction for multiple comparisons using the native functionality of the respective packages. Significance was defined as FDR or adjusted-p < 0.05.

#### Brain Slice Electrophysiology

Adult (2-6 mo) control and MIR mice (N=6 animals per group) were deeply anesthetized with isoflurane and perfused transcardially with ice-cold, high-sucrose slicing solution containing (in mM): 26 NaHCO_3_, 1.25 NaH_2_PO_4_, 208 sucrose, 10 glucose, 2.5 KCl, 1.3 MgCl_2_, and 8 MgSO_4_. Mice were swiftly decapitated, brains extracted and immediately placed in oxygenated high-sucrose slicing solution. Coronal slices containing both sensorimotor cortex and dorsolateral striatum were cut at 300 μm and transferred to an incubating chamber containing artificial cerebrospinal fluid (ACSF) (in mM): 130 NaCl, 5 KCl, 1.25 NaH_2_PO_4_, 26 NaHCO_3_, 2 MgCl_2_, 2 CaCl_2_, and 10 glucose) oxygenated with 95% O_2_–5% CO_2_ (pH 7.2–7.4, osmolality 290-310 mOsm/L, 32–34°C). Slices were allowed to recover for 60 min and electrophysiological recordings were obtained at room temperature. Cortical pyramidal neurons (CPNs) in layers 2/3 or striatal medium-sized spiny neurons (MSNs) were visually identified using infrared illumination with differential interference contrast optics (IR-DIC). Whole-cell patch clamp recordings in voltage- or current-clamp modes were obtained using a MultiClamp 700B Amplifier (Molecular Devices) and the pCLAMP 10.5 acquisition software (Molecular Devices). For CPN recordings (in voltage and current clamp), the internal pipette solution contained (in mM): 112.5 K-gluconate, 4 NaCl, 17.5 KCl, 0.5 CaCl_2_, 1 MgCl_2_, 5 ATP (dipotassium salt), 1 NaGTP, and 10 HEPES (pH 7.2, 270–280 mOsm/L). For striatal MSN recordings (in voltage clamp only), the internal pipette solution contained (in mM): 130 Cs-methanesulfonate, 10 CsCl, 4 NaCl, 1 MgCl_2_, 5 MgATP, 5 EGTA, 10 HEPES, 5 GTP, 10 phosphocreatine, and 0.1 leupeptin (pH 7.2, 270 mOsm). Electrode impedance was typically 4-5 MΩ in the bath. Cells with access resistance exceeding 25 MΩ were discarded. Cells were first voltage clamped at -70 mV to assess passive membrane properties (membrane capacitance, input resistance and decay time constant) and measured by applying a 10 mV depolarizing step voltage command and using the membrane test function integrated in the pClamp10 software. Active membrane properties of CPNs (resting membrane potential, intrinsic excitability, and rheobase) were determined in current clamp mode and by injecting current pulses of increasing intensity. Rheobase (minimal current intensity required to evoke an action potential), consisted of a single depolarizing current pulse (5 ms duration) of increasing intensity (Δ=15 pA). Spontaneous synaptic currents were recorded at -70 mV holding potential in the presence (MSNs) or absence (CPNs) of Bicuculline (BIC, 10-20 µM), a competitive antagonist of GABAA receptors, to better isolate glutamatergic events. Spontaneous synaptic currents were analyzed using the MiniAnalysis software (version 6.0, Synaptosoft, Fort Lee, NJ). The effects of rapamycin *ex vivo* (1-2 µM) on synaptic activity were examined by incubating slices from control or MIR mice for 2 hr before and during electrophysiological recordings. *In vivo* rapamycin treatment was also given to MIR mice (5 mg/kg, I.P.) or vehicle (DMSO) 2 hours before decapitation and slice electrophysiology. Primary outcomes for synaptic activity were the frequency and amplitude of spontaneous excitatory postsynaptic currents. For intrinsic excitability the primary outcomes were rheobase measurements and the firing patterns following injection of depolarizing current pulses.

#### Magnetic Resonance Imaging

Adult MIR and Control offspring were imaged using a 7 Tesla (T) Biospec small animal MRI system using Paravision 5.2 software (Bruker) at the UCLA Brain Mapping Center. Mice were briefly anesthetized with 2% isoflurane vaporized in oxygen flowing at 1 L/min, then placed on an MRI-compatible cradle and a single-channel surface coil (Bruker) secured over the head. For resting-state scans, a 3 mm-thick agar gel cap (Sigma, 3% in distilled water) was placed between the head and the surface coil, in order to reduce signal distortion in the blood-oxygen-level-dependent (BOLD) signal^171^. To minimize time-dependent effects of the anesthetic on the BOLD signal, these initial steps were performed within a 10–15-minute time window^172^. Isoflurane was gradually discontinued and sedation was initiated with a single subcutaneous (s.c.) injection of dexmedetomidine (Dexdomitor®, Zoetis; 0.15 mg/kg) followed by continuous s.c. infusion at 0.3 mg/kg/hr throughout the duration of the imaging as described in a previous publication^171^. Respiration and body temperature of the mouse were continuously monitored remotely, and maintained in a physiological range (37 ± 1 °C; Small Animal Instruments Inc.) by a homeothermically-controlled forced warm air over the body (SA11 Instr, Inc., USA). At the end of the imaging session, sedation was reversed by atipamezole (Antisedan®, Pfizer) at 1.5 mg/kg (i.p.). Data were acquired using the S116 Bruker gradients (400 mT/m) in combination with a single-channel surface coil (described above) and a 72 mm birdcage transmit coil. An initial series of scans was performed to confirm proper head position, then localized FASTMAP shimming was performed to improve field homogeneity. T2-weighted structural scans were acquired with a Rapid-Relaxation-with-Enhancement (RARE) sequence (RARE factor=8, Echo time (TE)=56 ms, repetition time (TR)=6,018 ms, 4 averages, field-of-view (FOV)=1.8× 1.2cm, slice thickness=0.5 mm, 18 slices, FA=90 deg, bandwidth (BW)=50 kHz, matrix=60×40). Then, 45-minute long functional (BOLD) data were acquired using the same image geometry as the structural scans, with a one-shot, interleaved, gradient-echo echo planar imaging sequence with the following parameters: TE=16 ms, TR=2,000 ms, FA=70 degrees, BW 400 kHz and a data matrix of 60×40). 10 dummy scans were used to allow the T1 signal to reach steady-state prior to signal acquisition. Resting-state functional data were acquired using above-described parameters, with the following modifications: field-of-view (FOV)=3.0×3.0 cm, matrix=128×64; functional (BOLD) scans acquired using a two-shot gradient-echo echo-planar imaging (GE-EPI) sequence, TE=19 ms, TR=1,000 ms (2,000 ms per volume), FA=70 degrees, BW 400 kHz and a data matrix of 128×64, Fourier transformed to 128×128, voxel resolution 0.23×0.23×0.5 mm). For each mouse, baseline data were collected and on the following week rapamycin (5mg/kg) or vehicle control (DMSO) was injected i.p. 2 hours before they were imaged in an identical manner.

#### MRI Analyses

Data were converted to nifti format and entered into a python-scripted, preprocessing pipeline to do the following steps: brain extraction^173^, movement correction^174^ and co-registration to a study-created brain template^175^, slice timing correction, Gaussian smoothing to 0.8mm, bandpass filtering 0.01-0.2Hz using the FSL Toolbox^176^. Data were masked for inclusion of brain voxels common to all mice to prevent bias. A mouse brain atlas (Ekam Solutions, USA) was registered to the study brain template, masked for common brain regions, and then inverse warped back to subject space. Functional adjacency matrices were calculated from the pre-processed, time-series rsfMRI data using 136 brain regions common to all brains to obtain Pearson correlation coefficients that were then Fisher z-transformed. Statistical contrasts were generated between MIR and Control mice and within group between baseline and rapamycin (rapa) intervention using network-based statistics^177^ implemented in GraphVar^178^. Subnetwork edge connectivity was evaluated at P<0.01, 2-tailed, using non-parametric permutation testing over 1000 iterations. Seed analysis was conducted by temporal correlation of whole brain data to mean signal extraction from regions of interest at the pre-processed subject-level data, and then at the group-level within the general linear model framework implemented by using FSL Feat, with statistical correct at 2.1z, P<0.01 cluster-based correction^179^. Distribution of brain networks into a community architecture was evaluated by calculation of modularity using 1000 iterations of the Louvain algorithm^180^. Modules derived from this process were then further classified at the nodal level by computing the classification diversity and classification consistency as described previously^181^ as implemented under GraphVar.

Tensor-based deformation analysis: Structural RARE data were converted to nifti image format, brain extracted from extracranial tissue using FSL BET^182^. All T2-weighted anatomical data were used to construct a study-specific, mean deformation template (MDT) images using the ants Multivariate Template Construction script^183^ from Advanced Normalization Tools (ANTs, v.2.3.3)^184^ that consisted of bias field correction and a three-stage coregistration procedure of rigid, affine and non-linear deformable registrations. The raw data were then separately coregistered to the MDT template using a two-stage affine and symmetric diffeomorphic registration under ANTs. The resulting deformation field was then used to derive the Jacobian determinant that provides an indication of the local tissue expansion or atrophy compared to the MDT. All data were masked for a common space to ensure a contribution from all mice to each voxel, following which statistical group differences in voxel-based Jacobian values were computed using non-linear permutation testing implemented under FSL Randomise^185^ under the general linear model framework using cluster-based thresholding set at z=3.1 (P<0.001) and variance smoothing of 0.1mm. Data were then corrected for multivoxel comparison using false discovery rate correction at q=0.01^186,187^. The primary outcomes for resting-state fMRI was functional connectivity changes assessed by network-based statistics and by network modularity measurements at baseline and after acute rapamycin treatment. The primary outcome for structural imaging were voxel-wise tissue volume changes (Jacobian determinant) in multiple brain regions.

#### Immunofluorescence

Brains were flash frozen and cut on a cryostat at 30uM. Brain sections were post-fixed on slides with 4% paraformaldehyde before being permeabilized with 3% TritonX-100 and immunostained with IBA1 (Wako 019-19741), NeuN (ThermoFisher), and Egr1 (Cell Signaling Technology) primary antibodies followed by the appropriate Alexa fluorescent secondary antibodies (ThermoFisher), and Hoescht counterstain. Positive cells were quantified on a Zeiss AxioImager Z1 fluorescent microscope with Apotome 2.0, using the unbiased optical fractionator approach (StereoInvestigator; MicroBrightField).

#### Statistical Analyses

Parallel group design was used for most experiments where there are separate MIR and Control offspring groups for every treatment and for different ages (young adult and old adult). Therefore, all univariate data were tested for normality and transformed to Gaussian where required. Behavioral data were analyzed by 2-WAY analysis of variance with repeated-measures, two-tailed paired t-tests, and mixed effects analysis for multiple comparisons when appropriate (GraphPad v8). Following an overall significant (P<0.05) group and/or interaction effect, Tukey’s and Sidak’s post-hoc tests were used to evaluate changes between groups and factors to account for multiple comparisons. Effect size (cohen’s d) was calculated for each result.

## Results

### Chronic pathophysiology in MIR offspring

In our mouse model of mild maternal inflammation, we have previously shown that juvenile and young adult mice have mild brain overgrowth and increased PI3K/AKT/mTOR pathway activation in the neurogenic SVZ niche^118^. In our current study we have observed that the pathophysiologic changes caused by exposure to MIR continue to persist and evolve throughout adulthood in the offspring. These changes include gross brain weight and in regional volumetric brain sizes (Fig2A-C), chronically elevated blood cytokines (Fig 1B) and brain microglia (Fig 1D), chronic mTOR pathway activation indicated by elevated brain phospho-S6 levels in western blots (Fig 1E), decreased reciprocal social interactions (Fig S1B), increased repetitive behaviors (Fig 1F,3C, 5A, 5C-D, S1B), and abnormal sensory behavioral responses in a tactile avoidance test, Von Frey fiber limb withdrawal test, and pre-pulse inhibition to startle test (Fig. 4A-F).

**Figure 2.**
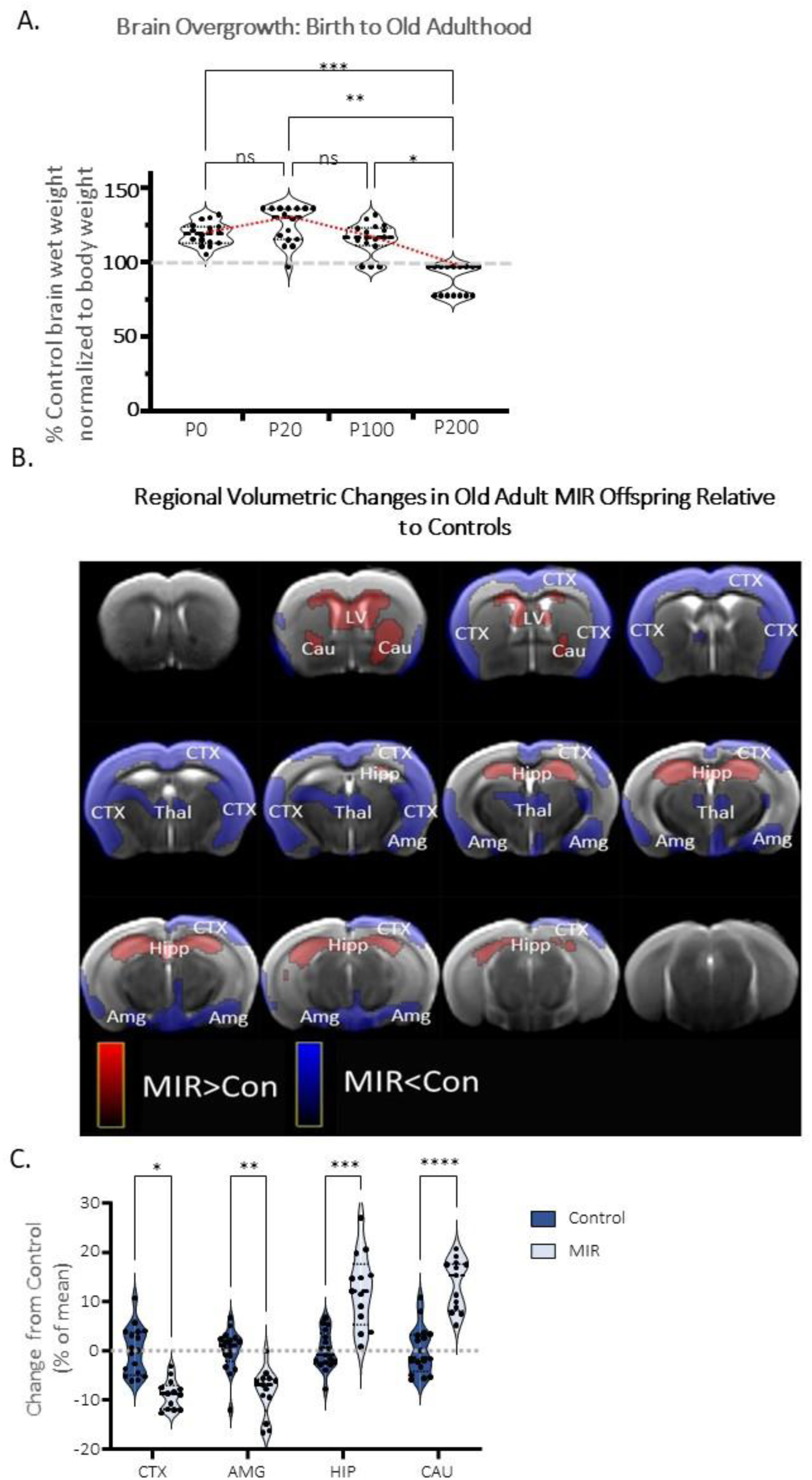
MIR offspring demonstrate growth patterns consistent with reports in humans with ASD of early brain overgrowth followed by undergrowth with cortical thinning and enlargement in other subcortical structures in adulthood. (**A**) Brain wet weights normalized to body weight of MIR offspring relative to control offspring from birth to post-natal day 200 (P200) show early overgrowth with most MIR brains being larger than average control brain (gray 100% line). A one-way ANOVA with multiple comparisons was overall significantly different between brain ages (p<0.0001, F_(3, 60)_) with a significant (Tukey’s) decrease in brain size at P200 compared to P0 (***p<0.0001), P20 (**p<0.0001), and P100 (*p<0.0001) but no significant size differences (ns) between P0 and P20 (p=0.3602) or P20 and P100 (p=0.0620), ages where brain overgrowth was maintained, n=16/group (**B**) Diffusion weighted MRI regional volumetric brain analysis of adult offspring (P400) showing mean percent differences between MIR and Control mice (blue = decreased red = increased brain volume) and variable size changes based on brain region with significant (p<0.0001) decreased cortical thickness (CTX), thalamus (Thal) and amygdala (Amg) volumes but significant (p<0.05) increases in lateral ventricle (LV), caudate (Cau), and hippocampus (Hipp) size, (**C**) Quantification of these regional deformation changes were significant overall by brain region and group in a 2-way ANOVA with multiple comparisons (brain region x group, p<0.0001, F_(2, 66)_, brain region, p<0.0001, F_(2, 66)_, group, p=0.0106, F_(1,28)_) with significant decrease in volume in MIR offspring compared to controls post-hoc tests (Tukey’s) in cortex (CTX, *p<0.0001, DF=27), amygdala (AMG, **p<0.0001, DF=24) and significant increases in volume in MIR compared to control in the hippocampus (HIP, ***p<0.0001, DF=17) and caudate (CAU, ****p<0.0001, DF=25), n=17/group.

**Figure 3.**
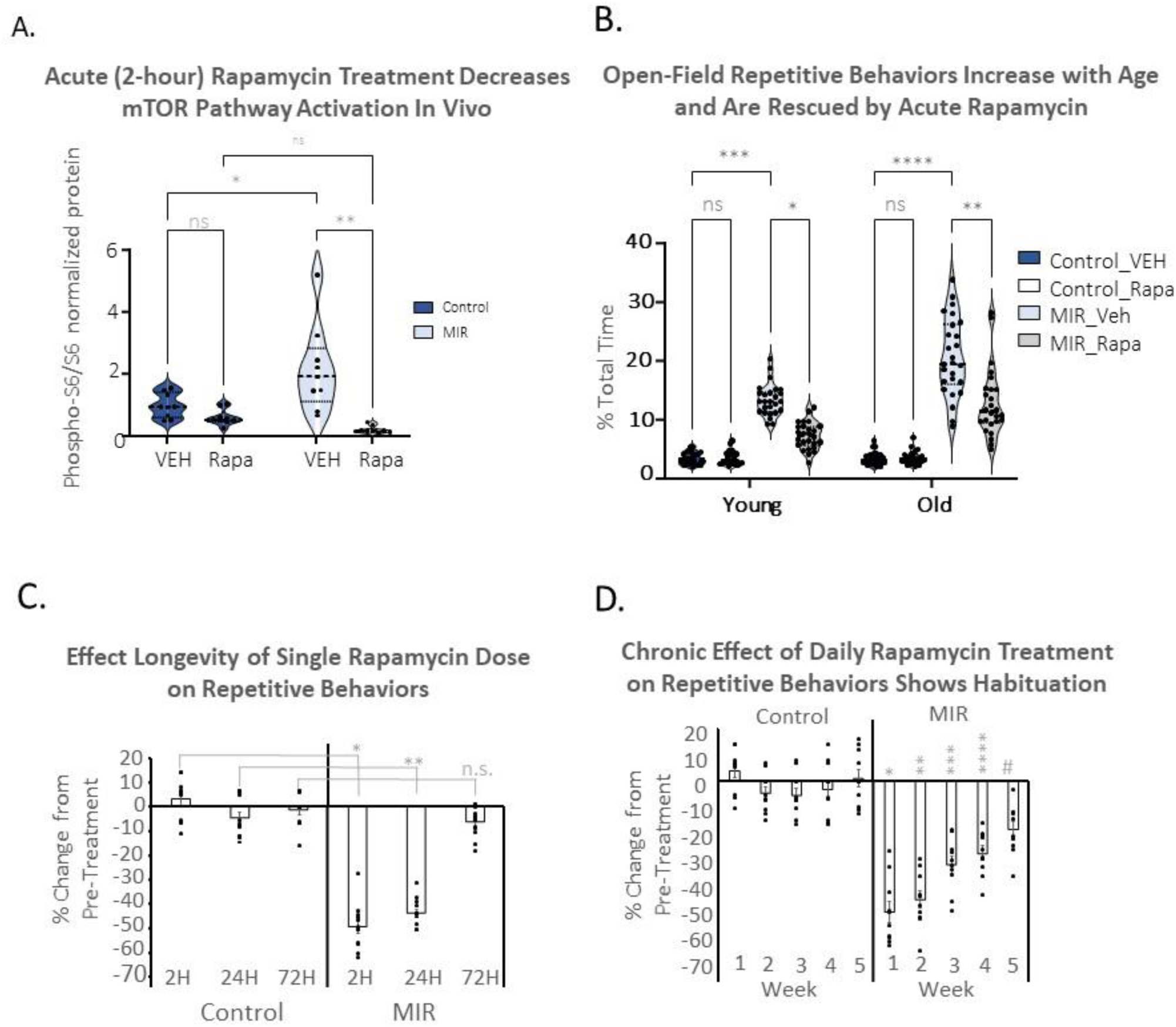
Acute (2-hour) rapamycin treatment rescues elevated brain mTOR signaling and repetitive and social behavior abnormalities in young and old adult MIR offspring. **(A)** Two-way ANOVA analysis for multiple comparisons of western blot data from pooled brain tissue (cortex, striatum, and amygdala) showing overall significant differences in treatment x group (p<0.0053, F_(l_,_16_)) and treatment (p=0.0002, F_(l,16)_). Post-hoc analysis (Tukey’s) analysis shows a significant increase in normalized phospho-S6/S6 in MIR offspring compared to controls treated with vehicle (*p=0.0020, DF=32) and a significant reduction in MIR mice treated with acute rapamycin (rapa) compared to vehicle (VEH; **p<0.0001, DF=16), demonstrating that MIR mice have chronically upregulated mTOR pathway activation and that rapamycin gets into the brain and reduces that activation within the 2 hour acute treatment time-frame, n=3/group/brain region; **(B)** Two-way ANOVA analysis for multiple comparisons of open field repetitive behaviors shows an overall significant difference in age x group (p<0.0001, F_(3_,_100)_), age (p<0.0001, F_(l,100)_), and group (p<0.0001, F(3, 100)). Post-hoc analysis (Tukey’s) shows a significant increase in repetitive behaviors in MIR mice treated with vehicle (VEH) compared to controls treated with vehicle in young (***p<0.0001, DF=200) and old (****p<0.0001, DF=200) mice and a significant decrease in behaviors in MIR mice treated with acute rapamycin (Rapa) compared to MIR mice treated with vehicle in young (*p<0.0001, DF=200) and old (**p<0.0001, DF=200) offspring, n=26/group.(**C**) Two-way ANOVA analysis for multiple comparisons of open field repetitive behaviors over time after a single rapamycin treatment indicate an overall significant time x group (p<0.0001, F(2, 39)), time (p<0.0001, F(2, 39)), and group (p<0.0001, F_(l,22)_) effects. Post-hoc (Tukey’s) analysis shows significant difference in Control vs MIR offspring at 2 hours (2H, *p<0.0001, DF=22) and 24 hours (24H, **p<0.0001, DF=20) but not at 72 hours (72H, p=0.0738, DF=22); n=12/group, data shown as mean+/- SEM; (**D**) Two-way ANOVA analysis for multiple comparisons of open field repetitive behavior after repeat daily doses of rapamycin show overall time x group (p<0.0001, F_(3, 48)_), time (p<0.0001, F_(3, 48)_), and group (p<0.0001, F_(l,18)_) effects. Post-hoc analysis (Tukey’s) indicate that MIR offspring treated with rapamycin had significantly greater effect of repetitive behaviors compared to controls offspring treated with rapamycin at 1 week of daily treatment (W1, *p<0.0001, DF=16), week 2 (W2, **p<0.0001, DF=16), week 3 (W3, ***p<0.0001, DF=17), week 4 (W4, ****p<0.0001, DF=17), and week 5 (WS, #p=0.0006, DF=18) although the difference gets smaller each week, indicating a gradual tolerance and loss of efficacy on repetitive behaviors over time with chronic administration. N=l0/group; Data shown as mean +/-SEM.

**Figure 4.**
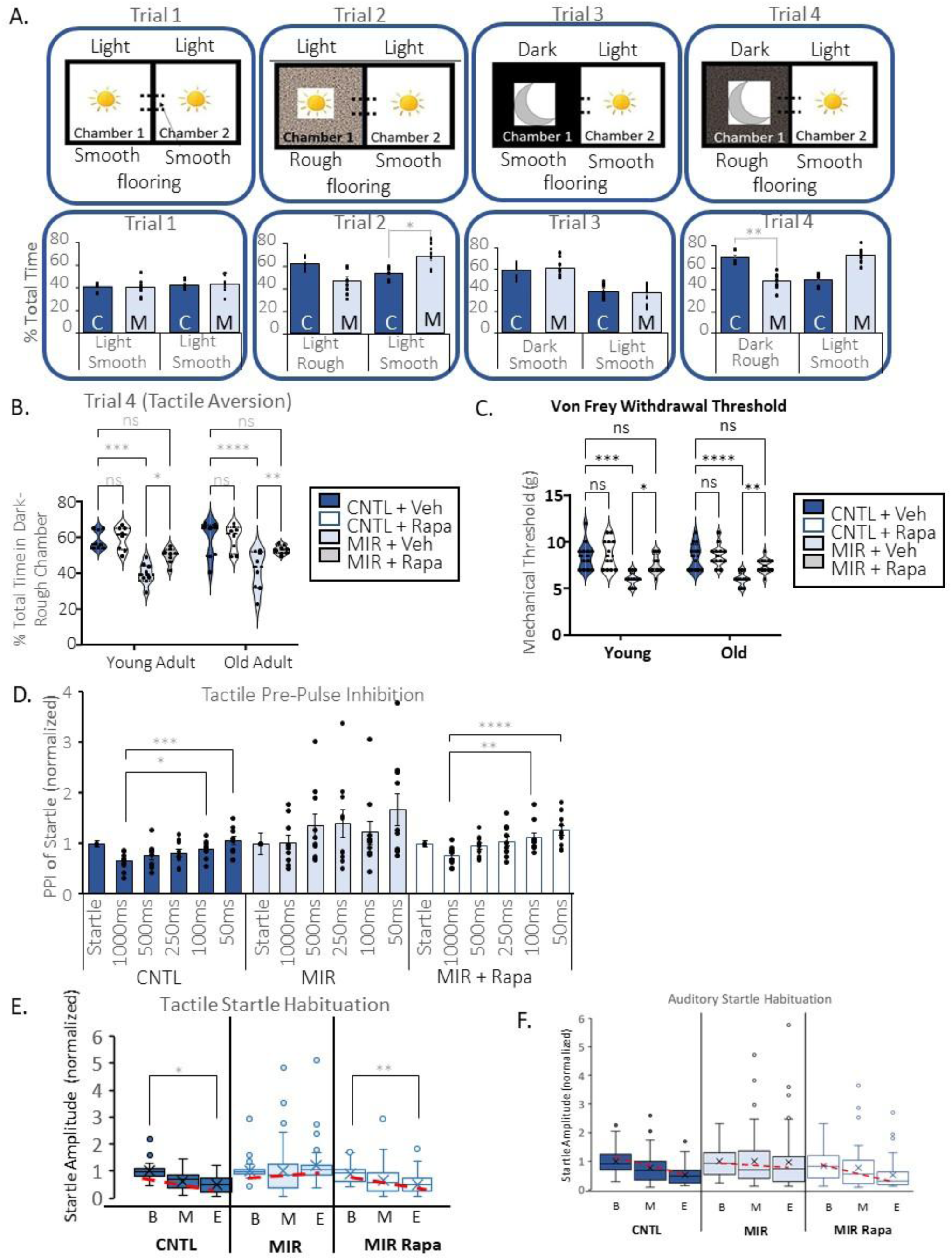
MIR mice display significant sensory over-responsivity (SOR), a common phenotype in ASD, which is rescued by acute rapamycin treatment. **(A)** A novel tactile avoidance test was used to assess MIR and control offspring for aversion to rough tactile textures on the hind-limbs and fore-limbs using a light/dark box with varying light and floor textures. This demonstrates that MIR mice have normal exploratory behavior (trial 1), normal time spent in a chamber with rough flooring when both chambers are lit the same (Trial 2), normal time spent in a dark rather than light chamber when the flooring is the same, (Trial 3), and time spent in the normally preferred dark chamber when the flooring is a rough (Trial 4). The percent total time spent in one chamber was analyzed for MIR and control offspring in young adult mice in each of these trials using a two-tailed paired t-test which shows that there is no significant difference in the trial 1 conditions (p=0.7074, DF=7) and trial 3 (p=0.5272, DF=7), but a significant difference in trial 2 (*p=0.0083, DF=7), in trial 4 (**p=0.0002, DF=7) with MIR mice spending significantly less time in the dark-rough chamber than control mice, n=8/group. **(B)** A two-way ANOVA analysis for multiple comparisons of the percent of total time spent in the dark-rough chamber in trial 4 found an overall significant group effect (p<0.0001, F_(3, 28)_). Post-hoc comparisons (Tukey’s) show that MIR mice spend significantly less time in dark-rough chamber in both young (p<0.0001, DF=56) and old (p<0.0001, DF=56) offspring and MIR mice treated with acute rapamycin spent significantly more time in the dark-rough chamber than MIR mice treated with Vehicle (Veh) in both young (p=0.0380, DF=56) and old (p=0.0102, DF=56). MIR mice treated with rapamycin were not significantly different to control mice treated with vehicle, indicating a significant rescue effect in both young (p=0.0882, DF=65) and old (p=0.3626, DF=65) mice, n=8/group.(**C**) A two-way ANOVA analysis for multiple comparisons of Von Frey limb withdrawal threshold found an overall significant effect of group (p<0.0001, F_(3,44)_). Post-hoc analysis (Tukey’s) shows that MIR mice treated with vehicle have a significantly lower threshold compared to control mice treated with vehicle in both young (***p<0.0001, DF=88) and old (****p<0.0001, DF=88) offspring and a significantly increased threshold in MIR mice treated with acute rapamycin compared to MIR mice treated with vehicle in both young (*p=0.0073, DF=88) and old (**p=0.0123, DF=88) mice. There also was no significant difference between control mice treated with vehicle and MIR mice treated with rapamycin in both young (p=0.2293, DF=88) and old (p=0.2223, DF=88) mice indicating a rescue effect by the acute treatment, n=12/group. (**D**) A two-way ANOVA analysis for multiple comparisons of the amount of startle amplitude inhibition produced by pairing a tactile pre-pulse (air puff) “warning” at decreasing time intervals with an audible startle (the pre-pulse inhibition test, PPI) shows overall significant interval (p=0.0015, F_(2, 65)_) and group (p=0.0132, F_(2, 27)_) differences. Both control mice treated with vehicle and MIR mice treated with rapamycin show significant changes in startle response at l000ms vs l00ms prepulse (control adjusted *p=0.0041, DF=9 and MIR+Rapa *p=0.0211, DF=9) and at l000ms vs 50ms (control adjusted ***p=0.0042, DF=9 and MIR+Rapa adjusted ****p=0.0053, DF=9) in Tukey’s post-hoc analyses and the normal trend of increased startle inhibition with decreasing prepulse delay. MIR mice treated with vehicle on the other hand were very variable in startle response and no significant changes in any startle responses between prepulse delay times was found, n=l0/group; **(E)** Startle response (amplitude normalized to beginning startle) to a tactile (airpuff) startle alone at the beginning (B), middle (M), and end (E) of the PPI test was used to assess sensory habituation. Two-way ANOVA analysis for multiple comparisons found significant time x group (p=<0.0001, F_(4, 268)_), time (p=0.0003, F_(2, 268)_), and group (p<0.0001, F_(2, 147)_) effects. Post-hoc analysis (Tukey’s) shows that there are significant decreases in startle response to the airpuff startle at the beginning vs the end startle periods for control mice treated with vehicle (adjusted *p<0.0001, DF=49) and MIR mice treated with rapamycin (adjusted *p<0.0001, DF=49) but no significant difference for MIR mice treated with vehicle (adjusted p=0.1835, DF=49). **(F)** Startle response to audible startle alone at the beginning, middle, and end of the PPI test was also used to assess sensory habituation. Two-way ANOVA analysis for multiple comparisons found significant effects of time (p=0.0005, F_(4, 288)_) and group (p=0.0145, F_(2, 288)_)-Post-hoc analysis (Tukey’s) shows that there are significant decreases in startle response to the audible startle between the beginning and end startles in the control (adjusted p=0<0.0001, DF=49) and MIR mice treated with rapamycin (adjusted p=0.0011, DF=49) but no significant difference in MIR mice treated with vehicle (adjusted p=0.9757, DF=49), n=l0/group.

In order to determine what might be driving this long-term dysfunction in our neuro-inflammatory MIR model we examined blood cytokines in the young and old adult MIR offspring compared to age-matched control offspring and found evidence of an ongoing, chronically elevated systemic inflammatory state with several pro-inflammatory cytokines remaining elevated in blood at all ages (Fig 1B). Large-fold increases in several pro-inflammatory cytokines were observed in blood samples from MIR offspring compared to controls in both young and old adult mice in G-CSF, IFN-g, IL-6, MCP-1, MIG, KC, M-CSF, GM-CSF and there were also smaller but significant increases in IL-9, IL-1beta, TNF-alpha, VEGF, IP-10, and LIF (Fig 1B). The largest increases in pro-inflammatory cytokines in the young adult MIR mice were in IFN-g, MCP-1, and IL-6 and in old adult mice were M-CSF, MCP-1, and G-CSF (Fig 1B). Acute 2-hour treatment with the mTOR inhibitor rapamycin had negligible effects on blood cytokines in adult MIR and control offspring (Fig S1A). Therefore, the effects of acute rapamycin in our experiments are not directly related to reducing the on-going systemic inflammation in MIR offspring, but still could be due to a reduction in the effects of this inflammation. Immunohistochemistry staining and quantification of microglia (IBA1+) further showed that chronically elevated neuro-inflammation with increased brain microglia populations from before birth to older adulthood in MIR offspring compared to controls (Fig 1D). Furthermore, western blot analysis of mTOR pathway activation in pooled tissue microdissections from sensory-motor cortex, striatum, and amygdala, brain regions showed elevated mTOR pathway activation in the MIR brains compared to controls (Fig 1E).

One of our hypotheses about the mechanisms behind the effects of MIR on behavior was that it may be driven by the increased inflammation seen in the blood and brain throughout the lifespan of these mice. Increased microglia in the brain in response to this chronic state could be one mechanism driving abnormal mTOR pathway activation in the brain and the abnormal behaviors observed in MIR offspring. Therefore, we used the CSFR1 inhibitor, Plexxikon5622, to deplete brain microglia in order to test this hypothesis. We found that ablation of microglia in young adult mice partially rescues repetitive, social, and sensory abnormalities (Fig 1F-G), suggesting that microglia do play a role in the pathophysiology of MIR exposure. However, microglia ablation does not significantly rescue these behaviors in older adult MIR mice even though rapamycin treatment does rescue them at this age, as described in the following section (Fig 1F-G), suggesting that while microglia may play a role in the development and evolution of behavioral abnormalities, they are unlikely to be the primary mediator of ongoing behavioral alterations.

Exposure to mild maternal inflammation is the only pre-clinical model to date that produces a pattern of brain growth changes that have been reported to occur in 50-70% of humans with ASD, namely mild brain overgrowth at young ages followed by a decrease in growth at older ages^98^. This can be seen in the MIR mice in both gross brain wet weight measurements from birth to just over a year in age (Fig.2A) and in regional volumetric analysis of individual brain structures using MRI structural imaging (Fig. 2B-C). Adult MIR mice show significant increases in the volumetric size of the hippocampus, caudate nucleus, and the amygdala but decreases in the thalamus and in sensory, motor, and association cortical areas (Fig. 2B-C). These regional changes are consistent with human volumetric studies which have identified similar changes in these structures in different adult autistic populations^188–194^.

### Acute Rapamycin Treatment Effects on ASD-related Behaviors

Because of the persistently elevated mTOR pathway activation in MIR offspring, we examined the effects of the mTORC1 inhibitor, rapamycin, on animal physiology and behavior. While most studies of the effects of rapamycin on mouse behavior have used repeated or chronic administration of the drug, we have focused on its acute effects in order to more fully understand whether it has effects that do not necessarily involve any significant structural changes to the brain or physical connectivity mediated by axonal outgrowth or synaptic remodeling. Therefore, we examined the effect of an acute 2-hour treatment with rapamycin on multiple ASD-associated behaviors. We confirmed via phospho-S6 western blot of pooled tissue microdissections from the cortex, striatum, and amygdala that a 2-hour treatment time frame allowed for the rapamycin to cross the blood brain barrier and significantly inhibit the mTOR pathway in the brain (Fig 3A). We then tested the rapamycin and vehicle-treated MIR and control offspring for reciprocal social interactions (Fig S1B) and repetitive behaviors (grooming and circling in an open field) in both young and old adult mice (Fig. 3B). We have previously demonstrated that MIR offspring have significantly reduced apparent social approach in the three chamber test and increased repetitive behaviors^118^. Here we found high levels of repetitive behaviors in MIR offspring that become more severe in old MIR offspring compared to young offspring (Fig 3B) and that acute rapamycin treatment significantly and markedly reduces repetitive behavior and social deficits at both young and old ages (Fig 3B, S1B). A single dose of rapamycin has a temporary rescue effect on repetitive behaviors with significant behavioral abnormalities returning after 72 hours in MIR offspring (Fig 3C). However, repeated daily treatment with rapamycin leads to an apparent tolerance effect over 5 weeks leading to a loss of effectiveness on repetitive behaviors (Fig 3D).

Additionally, we also determined that MIR offspring display behaviors consistent with sensory over-responsivity (SOR), a common and highly disabling ASD phenotype^195–198^ that has also been demonstrated in genetic mouse models of Fragile X and Rett Syndrome^159,199^. We showed that compared to control offspring, MIR offspring have a significantly increased aversion to rough tactile textures on the paws in a novel light/dark avoidance test (Fig 4A-B), a decreased threshold for limb withdrawal in the Von Frey fiber sensitivity test (Fig 4C), and abnormal sensory gating and lack of sensory startle habituation in the pre-pulse inhibition of startle test (PPI) using a tactile (air puff) pre-pulse and auditory and tactile startles (Fig 4D-F). These tests suggest significant sensory processing abnormalities in MIR offspring. These abnormalities were all at least partially rescued by acute rapamycin treatment in both young and old adult mice (Fig 4B-F).

### Central Nervous System Actions of Rapamycin

Sensory abnormalities in ASD models can be mediated specifically by peripheral neuron abnormalities^159,199^. Therefore, given the strong rescue effects that we observed on sensory over-responsivity in addition to other ASD-related behaviors in the MIR mice, we wanted to assess whether the drug was acting on the peripheral or central nervous system. To do this we tested the effects of RapaBlock, a small molecule that does not cross the blood brain barrier and prevents the effects of rapamycin on the peripheral nervous system^152,153^ on behavior in MIR and control animals. We found that there was no significant change in rapamycin rescue effects on repetitive behaviors, reciprocal social interactions, or sensory aversion with simultaneous RapaBlock treatment (Fig 5A-B, S1B), suggesting that the main effect of rapamycin is on the central nervous system in MIR mice.

**Figure 5.**
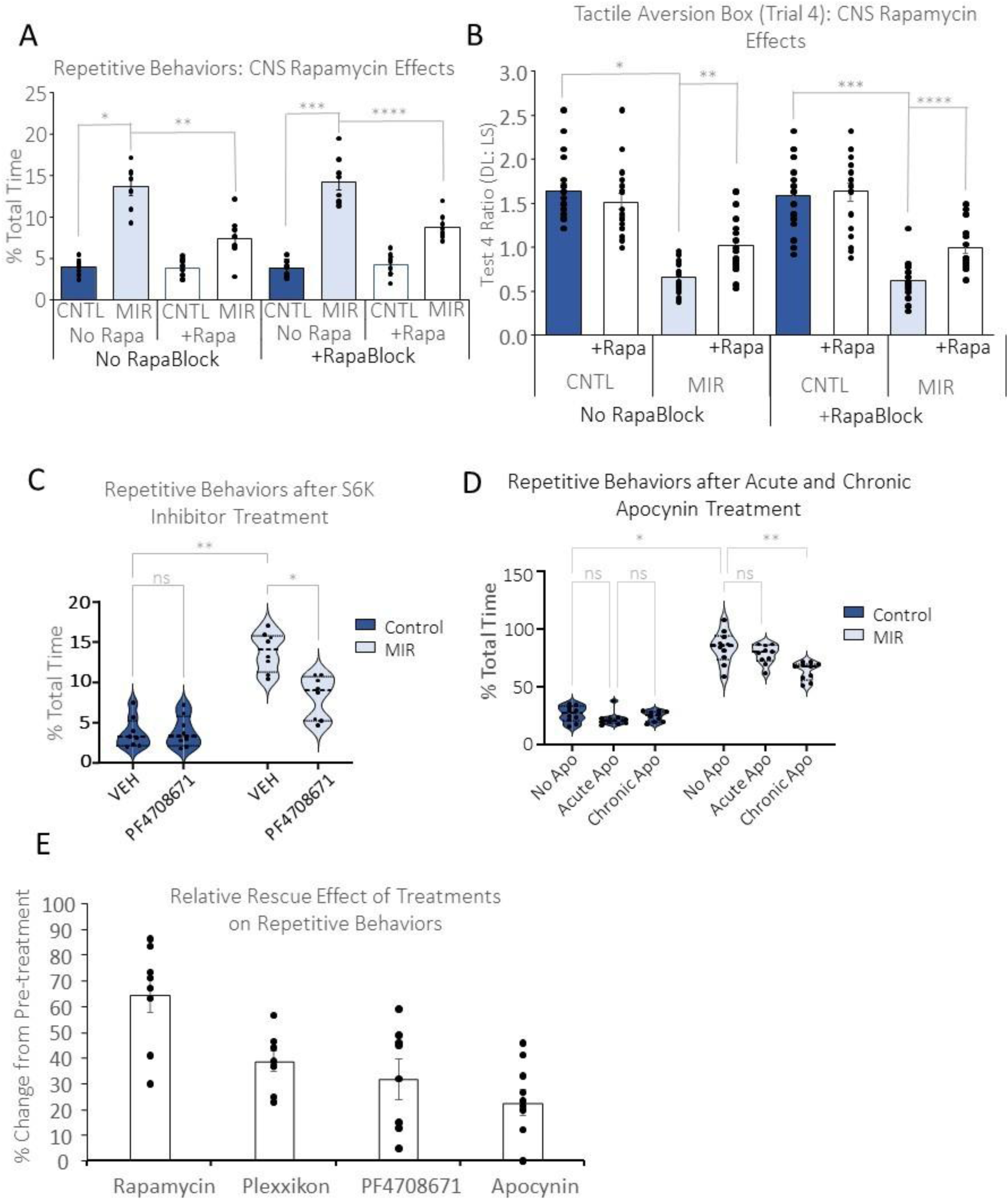
RapaBlock co-treatment with Rapamycin confirms the central nervous system action of rapamycin rescue of MIR offspring and S6K inhibition confirms that rescue of repetitive behaviors occurs via the mTOR signaling pathway. **(A)** A two-way ANOVA analysis for multiple comparisons of repetitive behaviors test in MIR and control offspring treated with vehicle or rapamycin in combination with RapaBlock or RapaBlock vehicle found an overall significant effect of Group (p<0.0001, F_(3, 56)_) but not in RapaBlock treatment (p=0.2398, F_(1, 56)_). Post-hoc analysis (Tukey’s) shows that MIR mice had significantly more repetitive behaviors than control offspring when there was no RapaBlock (vehicle, adjusted *p<0.0001, DF=56) or Rapablock given (adjusted **p<0.0001, DF=56) showing the abnormal MIR phenotype is not affected by Rapablock alone. MIR mice treated with rapamycin had significantly reduced repetitive behaviors compared to MIR mice treated with rapa-Vehicle when there was no RapaBlock (vehicle, adjusted ***p<0.0001, DF=56) or there was RapaBlock co-treatment (****p<0.0001, DF=56), suggesting that rapamycin is not acting on the peripheral nervous system in MIR mice. N=8/group, Data shown as mean+/- SEM. **(B)** A two-way ANOVA analysis for multiple comparisons of the ratio of time spent in the dark-rough vs light-smooth (Trial 4) chamber in the tactile light-dark box test shows there is an overall significant group effect (p<0.0001, F_(3, 136)_). Post-hoc analysis (Tukey’s) shows that MIR mice spent significantly less time in the dark-rough chamber than control offspring when there was no RapaBlock (vehicle, adjusted *p<0.0001, DF=136) or Rapablock given (adjusted **p<0.0001, DF=136) showing the abnormal MIR phenotype is not affected by Rapablock alone. MIR mice treated with rapamycin spent significantly more time in the dark-rough chamber compared to MIR mice treated with rapa-Vehicle when there was no RapaBlock (vehicle, adjusted ***p<0.0001, DF=136) or there was RapaBlock co-treatment (****p=0.0085, DF=136), also suggesting that rapamycin is not acting on the peripheral nervous system in MIR mice. N=18/group, Data shown as mean+/- SEM **(C)** Two-way ANOVA analysis for multiple comparisons of repetitive behaviors in an open field test in control and MIR offspring treated with the s6-kinase inhibitor PF4708671 or vehicle control shows and overall significant effect of treatment (p=0.0018, F_(1, 28)_), group (p<0.0001, F_(1,28))_, and interaction (p=0.0011, F_(1, 28)_). Post-hoc analysis (Tukey’s) shows a significant increase in repetitive behaviors in MIR mice compared to control offspring with vehicle treatment (**p<0.0001, DF=28) and significantly less repetitive behaviors in MIR mice treated with the S6K drug compared to MIR mice treated with vehicle (*p<0.0001, DF=28) but no effect of the drug on control offspring (p=0.8875, DF=28), confirming that downstream targets in the mTOR pathway may also produce some improvement in behavior, n=l0/group; **(D)** Two-way ANOVA analysis for multiple comparisons of repetitive behaviors in MIR and control offspring treated with acute and chronic apocynin, an inhibitor of the NADPH oxidase enzyme in the redox-mTOR pathway, shows that there are significant treatment x group (p=0.0006, F(2, 34)), treatment (p=0.0002, F(2, 34)), and group (p<0.0001, F(l, 18)) effects. Post-hoc analysis (Tukey’s) shows that MIR mice have significantly more repetitive behaviors compared to control offspring in apocynin (Apo) vehicle-control conditions (adjusted *p<0.0001, DF=13) and MIR mice have less repetitive behaviors in the chronic apocynin group compared to the MIR mice with vehicle-apocynin (adjusted **p=0.0061, DF=9) but not with the acute apocynin group (p=0.1589, DF=9), n=l0/group. **(E)** The relative rescue effects of rapamycin, Plexxikon 5622, and PF4708671 are compared demonstrating that the greatest effect on repetitive behaviors is from rapamycin; n=l0/group; data shown as mean± SEM

### mTOR Pathway Specificity of Effects

To confirm that rapamycin is having its rescue effects on MIR mice via its normal signaling to downstream mTOR targets like S6 Kinase and to rule out significant off-target effects, we administered the S6K inhibitor, PF4708671, according to the methods of Huang et al.^154^ and evaluated the effect on repetitive behaviors. We found that it does partially rescue repetitive behaviors after an acute 6-hour treatment (Fig 5C). This timepoint was chosen, rather than 2 hours, because there we found no measurable effect at 2 hours (data not shown) possibly due to drug absorption and brain penetration differences relative to rapamycin. We also tested the effects of acute (2H) and chronic (daily for 5 weeks) apocynin (Apo) treatment on repetitive behaviors (Fig 5D). We have previously shown that apocynin, an inhibitor of the NADPH oxidase enzyme, can reduce PI3K/AKT/mTOR pathway activity in the brain through post-translational redox regulation of the tumor suppressor PTEN^118,200^. We found that while acute Apo treatment of MIR mice did not significantly alter the repetitive behaviors in MIR offspring, chronic treatment did result in a significant decrease (Fig. 5D). However, a comparison of the relative levels of rescue of repetitive behaviors between rapamycin-treated, S6 kinase inhibition, microglia ablation, and chronic apocynin treatment in young adult MIR offspring indicates that rapamycin was significantly more effective than these other treatments (Fig 5E).

### Neuronal Hyper-excitability & Seizure Susceptibility

In order to understand potential physiological mechanisms underlying the behavioral changes in MIR mice and the effects of rapamycin we examined the electrophysiological properties of neurons in brain slice preparations. We found that cortical pyramidal neurons (CPNs) in layer 2/3 of the somatosensory cortex of adult MIR offspring displayed a depolarization shift, resulting in a resting membrane potential that is closer to the action potential threshold and decreased Rheobase (Table 1) and significantly increased frequency, amplitude, and area in spontaneous excitatory postsynaptic potentials (EPSPs, Fig 6A) and voltage-evoked responses (Fig S2A-B), which indicated a marked increase in neuronal hyper-excitability compared to controls. Under current-clamp conditions (depolarizing current injection of pulses 500-1000 ms pulses) the cortical neurons from MIR brains produced 4 different categories of action potential activity^201^: regular spiking-slowly adapting (RS-SA), regular spiking-fast adapting (RS-FA), regular spiking-doublet (RS-Doublet--when increasing current intensity evoked two action potentials at the beginning of the pulse), and intrinsic bursting (IB). In layers 2/3, CPNs generally belong to the RS-SA category which was the case for the neurons from control mice which had 75% RS-SA, 15% RS-FA, 10% RS-Doublet, and no intrinsic bursting (Fig 6B). CPNs from MIR brains had a dramatically altered spiking profile with mainly RS-Doublets (45%), only 40% RS-SA, 3% RS-FA, and 12% intrinsic bursting patterns which are not normally observed in upper layer neurons and was not seen in CPNs from control mice, indicating significantly altered firing patterns (Fig 6B). The effects of rapamycin on the hyper-excitable phenotype of MIR neurons was assessed via *in vitro* treatment where the rapamycin was added to the slice media (2uM) and via *in vivo* treatment (5 mg/kg) where the mice were injected with rapamycin I.P. 2 hours prior to euthanasia and tissue preparation. In vitro rapamycin significantly rescued the resting membrane properties of the CNPs from MIR offspring, normalizing the resting membrane potential and increasing Rheobase (Table 1). The increased frequency, amplitude, and area of spontaneous EPSCs in cortical neurons was significantly rescued following the *in vitro* rapamycin treatment (Fig 6C) and the frequency of EPSCs was similarly significantly reduced following *in vivo* rapamycin treatment (Fig 6D). Representative traces of spiking activity and bursting in cortical pyramidal neurons from MIR and control brains could be seen in voltage clamp and current clamp conditions in Fig S2A-B. Spontaneous EPSCs in striatal medium spiny neurons (MSNs) showed a trend of increasing frequency that didn’t reach significance (Fig S2C) and was reduced by acute *in vitro* rapamycin treatment (Fig S2D).

**Figure 6.**
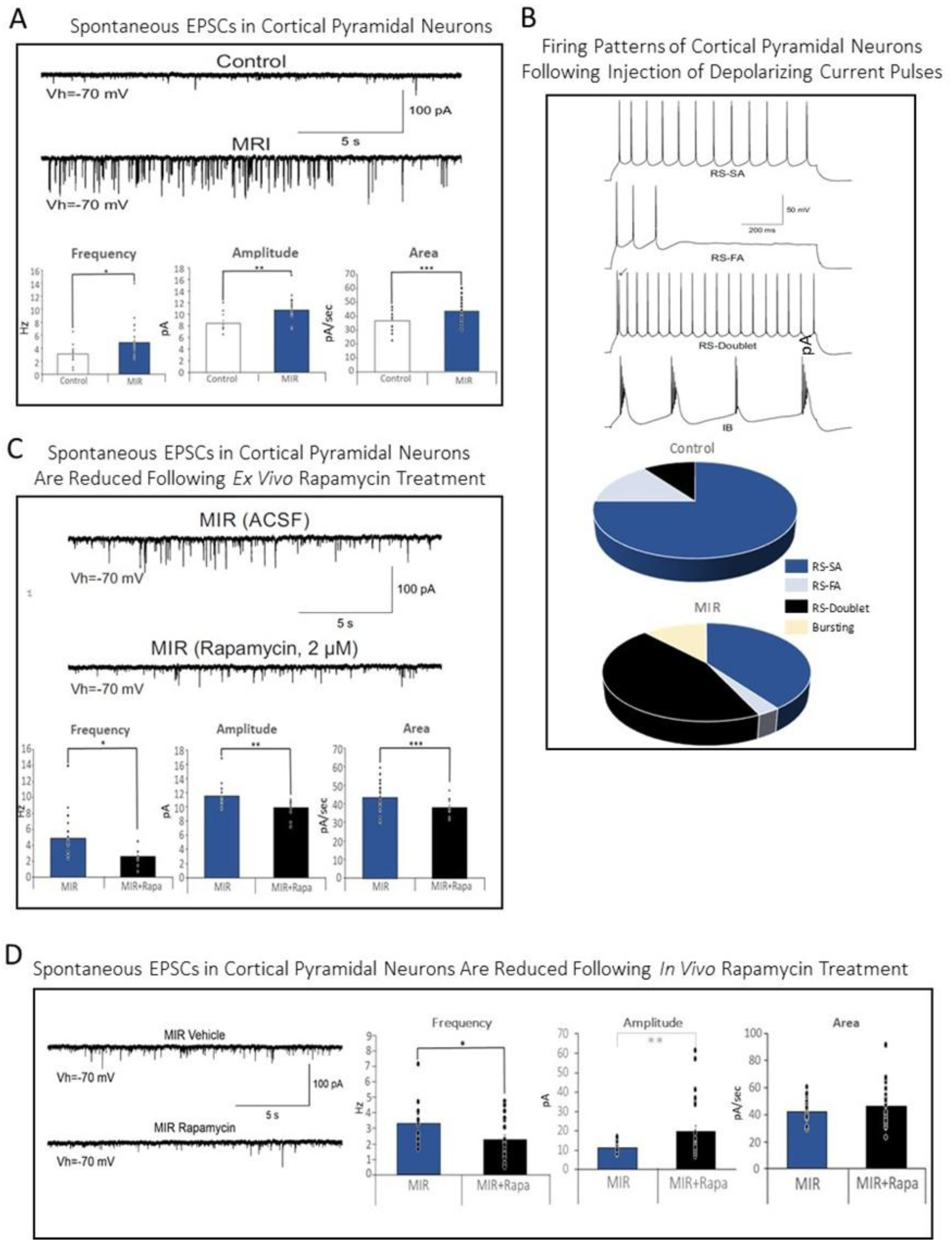
Cortical neurons in MIR offspring display abnormal electrophysiology indicating hyper-excitability which is rescued by acute rapamycin treatment. **(A)** A mixed effects analysis for multiple comparisons was performed on the quantification of the electrophysiological properties of spontaneous excitatory postsynaptic currents (EPSCs) in cortical pyramidal neurons and shows an overall significant effect by variable for fixed effects (p<0.0001, F_(1, 36)_) and group (p=0.0049, F_(1, 33)_). Post-hoc analysis (Sidak’s multiple comparisons) shows a significant difference between MIR and control mice for frequency (adjusted *p=0.0160, DF=29), amplitude (adjusted **p=0.0022, DF=24), and area (***p=0.0312, DF=21), n=24 neurons/group; **(B)** Firing Patterns of Cortical Pyramidal Neurons Following Injection of Depolarizing Current Pulses (500-1000 ms) show that layer 2/3 neurons from MIR mice have decreases in regular spiking slow and fast adaption neuron firing, increased doublet firing, and ectopic burst firing that is not present in the neurons from control mice; **(C)** A mixed effects analysis for multiple comparisons was performed on the quantification of the electrophysiological properties of spontaneous excitatory postsynaptic currents (EPSCs) in cortical pyramidal neurons and shows an overall significant effect by variable (p<0.0001, F_(l, 37)_) and group (p=0.0028, F_(1, 32)_. Post-hoc analysis (Tukey’s) shows a significant rescue effect on MIR neurons treated by ex vivo rapamycin (2uM) in slice culture compared to MIR neurons treated with vehicle during electrophysiology recording in frequency (adjusted *p=0.0013, DF=31), amplitude (adjusted **p= 0.0035, DF=29), and area (adjusted ***p=0.0167, DF=30), n=24 neurons/group. **(D)** A mixed effects analysis for multiple comparisons was performed on the quantification of the electrophysiological properties of spontaneous excitatory postsynaptic currents (EPSCs) in cortical pyramidal neurons and shows an overall significant effect by variable (p<0.0001, F_(2, 65)_). Post-hoc analysis (Tukey’s) shows a significant rescue effect MIR mice treated by in vivo rapamycin (5mg/kg 2 hours prior to brain slice preparation) compared to MIR mice treated with vehicle in frequency (adjusted *p=0.0146, DF=35), amplitude (adjusted **p= 0.0151, DF=24), but not in area (adjusted p=0.3925, DF=26). The following abbreviations were used: RS-SA: regular spiking (slow adapting), RS-FA: regular spiking (fast adapting), Hz: hertz, pA: picoamperes, mV: millivolts, s: sec: seconds, n=24 neurons/group All data mean ± SEM. N=6 animals/group/treatment

**Table 1.** Passive Membrane Properties of Cortical Pyramidal Neurons. Passive membrane properties (membrane capacitance, input resistance, time constant, and resting membrane potential (RMP)), and Rheobase) of cortical pyramidal neurons show MIR mice have significantly higher resting membrane potential (*p<0.05, closer to action potential threshold) and decreased Rheobase (**p<0.05, lower minimum current to depolarize) indicating neuronal hyper-excitability in the cortex and increased membrane capacitance in the striatum compared to controls; The following abbreviations were used: Cm: membrane capacitance, pF: picofarads, Rm: membrane resistance, MW: megaohms, Tau: time constant, ms: milliseconds, RMP: resting membrane potential, mV: millivolts, pA: picoamperes; all data mean ± SEM, N=24 neurons & 6 animals/group/treatment

|  | <u>Cm(pF)</u> | <u>RM(MΩ)</u> | <u>Tau(ms)</u> | <u>RMP(mV)</u> | <u>Rheobase(pA)</u> |
| --- | --- | --- | --- | --- | --- |
| Control | 208.8 ± 21 | 109.2 ± 16 | 4.0 ± 0.4 | -78.1 ± 2.4 | 725.6 ± 98 |
| MIR | 207.8 ± 82 | 129.0 ± 16 | 3.6 ± 0.3 | **-70.6 ± 1.3\*** | **499.2 ± 52\*\*** |
| MIR+Rapa | 214.5 ± 8 | 106.9 ± 8 | 3.7 ± 0.16 | 73.3 ± 1.4 | 590.6 ± 61 |

Because the striatum receives its main excitatory input from the cerebral cortex and, to a lesser extent, from the thalamus we wanted to examine if the cortical hyper-excitability observed in MIR mice was affecting the striatal MSNs. To do this we used Bicuculline (10uM) to block GABA_A_ receptors *ex vivo* in the slice media. We observed that the CPNs display paroxysmal (epileptic-like) discharges that propagate to the striatum and are manifested by large synaptic currents and bursts of spontaneous synaptic currents in MSNs. This indicates unmasked cortical hyper-excitability because control brains rarely displayed this type of MSN activity (Fig S2E). Acute treatment with rapamycin *ex vivo* also reduced Bicuculline-induced hyper-excitability in MSNs in MIR brains (Fig S2F).

Given this neuronal hyper-excitability observed in MIR offspring we also tested the effect of a dose escalation paradigm using the seizure-inducing drug, pentylenetetrazol (PTZ) to determine if MIR mice have brains that are more susceptible to seizure induction. The number of animals per group at each progressive seizure score following PTZ treatment was used to indicate relative seizure susceptibility between MIR and Control offspring. We found that MIR mice progressed up the seizure score ranking at significantly lower doses of PTZ than control offspring with all MIR mice (8/8) experiencing some level of seizure activity at 40 mg/kg compared to only 2/8 control mice (Table 2). Rapamycin treatment significantly reduced this seizure susceptibility in MIR mice, lowering the seizure score of mice but not eliminating the seizure-inducing effect of PTZ on the mice (Table 2, Supplemental Table 3). Taken together, our electrophysiology and seizure susceptibility studies indicate that adult MIR mice display abnormal neuronal hyper-excitability in cortical and striatal neurons that was normalized by acute rapamycin treatment, suggesting that one mechanism by which rapamycin acts acutely on the brain is via the rapid normalization of inhibitory/excitatory balance in the brain.

**Table 2.** Seizure Susceptibility by PTZ Dosage. . Seizure scores during pentylenetetrazol (PTZ) dose escalation indicates a greater severity and larger number of mice per group displaying a seizure behavior in MIR mice compared to Control offspring that is decreased by acute rapamycin treatment. N=8/group

| Maximum Seizure Score | PTZ dosage (mg/kg) |  |  |  |  |  |  |  |  |  |
| --- | --- | --- | --- | --- | --- | --- | --- | --- | --- | --- |
|  | 10 |  | 20 |  | 30 |  | 40 |  | 40 + Rapamycin |  |
|  | CNTL | MIR | CNTL | MIR | CNTL | MIR | CNTL | MIR | CNTL | MIR |
| 0 | 8/8 | 8/8 | 8/8 | 7/8 | 7/8 | 4/8 | 6/8 | 0/8 | 7/8 | 0/8 |
| 1 | 0/8 | 0/8 | 0/8 | 1/8 | 1/8 | 3/8 | 2/8 | 1/8 | 1/8 | 3/8 |
| 2 | 0/8 | 0/8 | 0/8 | 0/8 | 0/8 | 1/8 | 0/8 | 2/8 | 0/8 | 4/8 |
| 3 | 0/8 | 0/8 | 0/8 | 0/8 | 0/8 | 0/8 | 0/8 | 3/8 | 0/8 | 1/8 |
| 4 | 0/8 | 0/8 | 0/8 | 0/8 | 0/8 | 0/8 | 0/8 | 2/8 | 0/8 | 0/8 |
| 5 | 0/8 | 0/8 | 0/8 | 0/8 | 0/8 | 0/8 | 0/8 | 0/8 | 0/8 | 0/8 |
| 6 | 0/8 | 0/8 | 0/8 | 0/8 | 0/8 | 0/8 | 0/8 | 0/8 | 0/8 | 0/8 |

### Circuit-level functional dysregulation of MIR brains is altered by acute rapamycin treatment

A potential mechanism by which acute rapamycin treatment could rapidly affect brain function and behavior is through effects at the level of functional connectivity and network organization. We assessed young adult MIR and control offspring before and 2 hours after rapamycin treatment using resting state functional Magnetic Resonance Imaging (rsfMRI). Functional connectivity (r values transformed to Fisher z scores) across the brain were calculated as an indication of the connectivity strength between cortical and subcortical regions before and after acute Rapamycin treatment (Fig 7A). This comparison demonstrates that at baseline prior to rapamycin, MIR mice showed a general pattern of increased brain-wide FC compared to controls, especially within cortical and subcortical regions and within hemispheres between cortical-subcortical regions (Fig. 7A). The effect of acute rapamycin treatment dramatically increased FC within subcortical regions but decreased FC within cortical regions and between cortical and subcortical regions within each hemisphere. The effects of rapamycin on control brains was small by comparison (Fig. 7A). Thus, the changes on brain FC in MIR offspring from acute rapamycin treatment were large but do not produce a return to control FC conditions.

**Figure 7.**
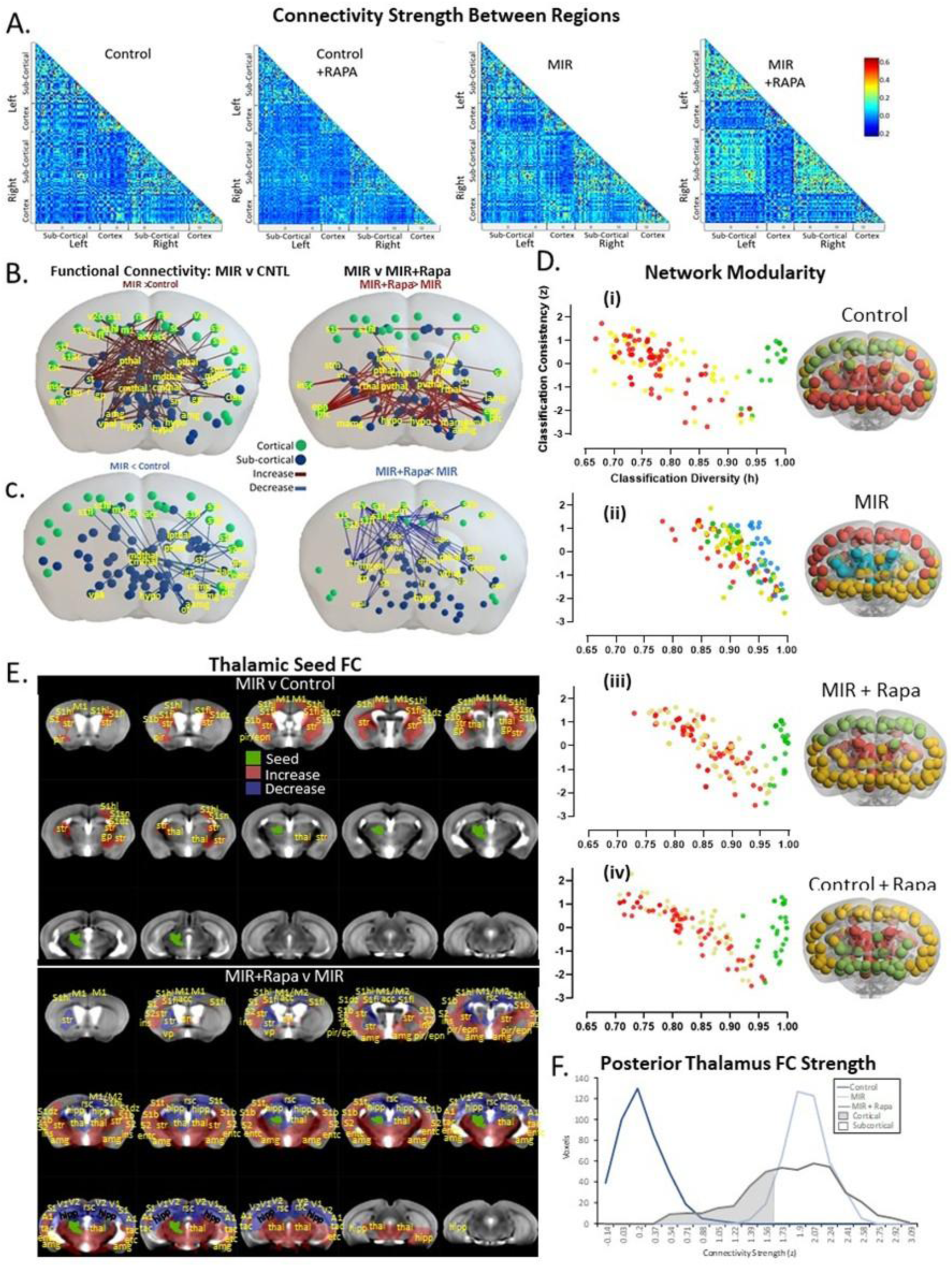
Disordered functional connectivity seen in MIR mice are significantly rescued by acute rapamycin treatment. **(A)** Fisher z scores (transformed r values, 0.2-0.6) indicating functional connectivity strength between cortical and subcortical brain regions in MIR and control mice before and after acute rapamycin treatment; **(B)** Regional increases in functional connectivity in resting state fMRI from MIR and control mice before and after acute rapamycin treatment (P<0.01 NBS, 2-tailed); (**C**) Regional decreases in functional connectivity in resting state fMRI from MIR and control mice before and after acute rapamycin treatment (P<0.01 NBS, 2-tailed); (**D**) Differential network modularity organization across the brain is increased in MIR mice compared to control offspring and is rescued by acute rapamycin treatment; (**E**) Differential functional connectivity by brain region from a left posterior thalamus seed in MIR mice compared to control offspring before and after acute rapamycin treatment; (**F**) Voxel-wise quantification of posterior thalamus functional connectivity is significantly increased in MIR mice compared to control offspring and is differentially affected by acute rapamycin treatment with cortical regions showing decreased thalamic FC and subcortical regions showing increased thalamic FC following treatment. Abbreviations used as follows: **acc**=anterior cingulate cortex; **rsc**=retrosplenial cortex; **sc**=subicular complex; **v2c**=secondary visual cortex; **s1s**=primary somatosensory cortex; **s2s**=secondary somatosensory cortex; **s1b**=Sl barrel cortex; **s1a**=primary auditory cortex; **dthal**=dorsal thalamus; **vthal**=ventral thalamus; **pthal**=posterior thalamus; **pvthal**=paraventricular thalamus; **mdthal**=medial dorsal thalamus; **rthal**=reticular thalamus; **lpthal**=lateral posterior thalamus; **tac**=temporal association cortex; **stm**=stria terminalis; **mgen**=medial geniculate; **lgen**=lateral geniculate; **str**=striatum; **cmthal**=central median thalamus; **clau**=claustrum; **gp**=globus pallidus; **sn**=substantia nigra; **amg** =amygdala; **lamg=**lateral amygdala; **hypo**=hypothalamus; **vpal**=ventral pallidum; **entc**=entorhinal cortex; **insc**=insular cortex; **slf** = primary face somatosensory cortex; **s1fl**=primary forelimb somatosensory cortex; **s1hl**=primary hindlimb somatosensory cortex; **s1t**=primary trunk somatosensory cortex; **m1**=primary motor cortex; **m2**=secondary motor cortex; **ot**=olfactory tubercles; **aamg**=anterior amygdala; **camg**=central amygdala; **bamg**=basal amygdala; **mamg** =medial amygdala; **pfc**=piriform cortex; **epn=**endopiriform nucleus; **s2c**=secondary somatosensory cortex; **supc**=superior colliculus; **fx**=hippocampal fornix; fim=hippocampal fimbria; **cgm=**cingulum; **hab**=habenular; **dg**=dentate gyrus; **ca3**=hippocampus CA3; **paqu**=periaqueductal gray; **rtf**=reticular formation; **lsep=** lateral septum, N=lS/group

Network based statistics (NBS) methodology^177^ (P<0.01 NBS, 2-tailed) was used to quantitatively examine these FC differences at a detailed, subnetwork level (Fig. 7B,C). This showed that the altered FC in MIR mice at baseline was primarily due to increases between cortical primary and higher-order (secondary) sensory processing areas (primary somatosensory and somatosensory for hind limbs, forelimbs, whiskers, trunk, and neck/shoulder, visual, auditory, taste, and olfactory) and subcortical structures (thalamus and basal ganglia). However, there were notable decreases in FC between sensory cortical areas and amygdala (Fig. 7B). Despite the strong repetitive behavior phenotype in the MIR mice, there were surprisingly more limited motor-related FC differences identified in MIR offspring compared to controls. The primary motor cortex only showed increased FC to retrosplenial cortex (sensory integration), subicular complex (memory), and the striatum (motor). The largest proportion of the FC increases in MIR mice relative to controls was seen between subcortical structures, primarily basal ganglia connections with thalamus and sensory cortex (Fig. 7B). FC decreases between MIR and control mice were sparser, and were again primarily sensory cortex changes, but to the thalamus and amygdala (Fig. 7B).

The largest effects of acute rapamycin on FC in MIR mice was a pattern of within subcortical increases in FC (Fig. 7C) and the opposite effect on the cortex with significant FC decreases (Fig. 7B,C). Acute rapamycin treatment had only a limited effect on control mice (Supplemental Fig.S3A). The significant decreases in FC in MIR mice from rapamycin were between somatosensory cortical areas (primary, forelimb, hind limb, trunk, barrel) and basal ganglia and thalamic structures (Fig 7C). The increases in FC in MIR mice following rapamycin treatment were between left and right hemisphere primary somatosensory cortex, from insular cortex (sensory salience), piriform cortex (olfaction), and endo-piriform nucleus (taste) to thalamic structures and between amygdala (limbic emotion) and thalamic and basal ganglia structures (Fig. 7C).

A functional modularity analysis was used to identify the community network architecture (groups of brain regions that are more strongly connected to each other than to other regions) for each experimental group where the number of modules identified indicate the functional specialization or segregation within that group’s brain network. MIR brains were found to have four main modules: cortical, (red), two subcortical/cortical (yellow, green) and a thalamic and basal ganglia module (blue) and a modularity index of 0.693 (Fig 7D(ii)). Control mice were found to only have three modules: anterior cortical (green), posterior cortical (yellow), and a mainly subcortical module (red) and a significantly lower modularity index of 0.511 (Fig 7D(i), p<0.01). The brain regions (nodes) that make up these modules were further quantitatively evaluated using classification consistency and diversity (CC, CD) parameters which respectively reflect: the degree to which each node was classified to the same versus different modules across all mice within a group (7D(i,ii,iii,iv)). With the exception of the cortex in control mice, MIR mice were characterized by all modules having high CD values relative to the controls, indicating increased inter-module connectivity in MIR mice that likely facilitates greater integration between the modules than in controls. (7D(ii)).

Acute rapamycin treatment had a dramatic effect on this community structure in the brain by normalizing the number of modules in MIR mice to 3 main modules that became broadly, spatially similar to controls (Fig 7D(iii)). This was marked by a whole brain reduction in modularity from 0.693 at baseline to 0.552 following acute rapamycin intervention (P<.05), which was approaching control baseline modularity of 0.511, indicating reduced segregation in the network. Within modules, rapamycin shifted the cortical nodes in the MIR mice to a significantly higher CD value that was very similar to the cortex in control mice at baseline (Fig 7D(ii,iii)). The two subcortical network nodes in MIR mice treated with rapamycin were shifted in the opposite direction to lower CD values and closer to controls values. In control mice, while rapamycin had no significant effect on modularity (0.511 to 0.533 in baseline and rapamycin treated conditions respectively), it had the opposite effect on nodes within modules compared to in MIR mice. It significantly reduced the cortical module CD values compared to baseline, but significantly increased subcortical CD values, indicating much greater flexibility (Fig 7D(i,iv).

A functional connectivity seed analysis from the posterior thalamus (multi-sensory processing, attention, motor control) to whole brain was conducted on the data from MIR and Control mice before and after acute rapamycin treatment (P<.01, cluster corrected). MIR mice at baseline were found to have significantly increased cortico-thalamic FC for primary motor cortex, piriform cortex (olfactory), and primary somatosensory cortex (S1, S1DZ, hind limb, forelimb, barrel, shoulder/neck) compared to controls (Fig 7E). There were also increases in FC from posterior thalamus to external globus pallidus (multi-sensory processing and motor control), dorsal striatum (multi-sensory processing and motor control), and contralateral anterior thalamus (somatosensory and proprioception sensory processing) compared to controls (Fig 7E). However, following acute rapamycin treatment these MIR FC increases were reversed for cortico-thalamic FC between primary motor cortex and primary somatosensory cortex (hind limb, forelimb), and from posterior thalamus to left striatum and to contralateral anterior thalamus (Fig 7E). Finally, rapamycin further increased cortico-thalamic FC in primary somatosensory cortex (S1, and barrel cortex) and piriform cortex and between posterior thalamus and dorsal striatum and external globus pallidus compared to MIR mice at baseline (Fig 7E). Rapamycin treatment bilaterally increased FC for several cortico-thalamic connections that weren’t differentially connected in MIR mice pre-treatment (insular cortex (salience network), and entorhinal cortex (memory, sensory salience, spatial navigation) and between posterior thalamus and amygdala, bilateral dorsal striatum and external globus pallidus (Fig 7E). Rapamycin treatment also bilaterally decreased posterior thalamus FC to the anterior cingulate cortex (salience network), retrosplenial cortex (visual/auditory sensory processing and associations), primary auditory cortex, primary and secondary visual cortex, temporal association cortex (memory, visual/auditory processing), ventral striatum (sensory integration with emotion/reward processing), and hippocampus (memory) which also weren’t differentially connected to the posterior thalamus in MIR mice prior to rapamycin treatment (Fig 7E). Therefore, rapamycin treatment’s effect on brain functional connectivity was largely found to be a reduction to thalamic-mediated sensory processing regions. Voxelwise rsfMRI mapping of the thalamic seed FC data showed a pattern of increased FC strength and hyper-connectivity in MIR mice compared to controls. Global histogram analysis of this data using the difference between MIR and control mice (Fig. 7E) confirmed a large shift in FC strength for both cortical and subcortical structures in MIR mice and a divergent effect of rapamycin treatment, both decreasing cortical hyper connectivity with the posterior thalamus and increasing connectivity for subcortical structures (Fig 7F).

In summary, our analysis of resting state functional connectivity and circuitry demonstrates that MIR mice have increased FC in widespread areas of the brain and this is largely normalized towards controls following an acute 2-hour treatment with rapamycin, at the expense of dramatically increased subcortical FC (Fig. 7A). Specific brain nodes that have increased functional synchrony in MIR mice compared to controls include retrosplenial cortex, auditory and somatosensory cortex (sensory) and cingulate cortex and amygdala (limbic), and insular cortex (sensory salience) areas (Fig. 7B) which is congruent with the sensory, anxiety, and social behavioral abnormalities seen in the MIR mice that are rescued by the acute rapamycin treatment^118^ (Fig. 3C-D). Significant decreases in FC in MIR mice compared to controls were seen between forelimb (motor), medial amygdala, striatum, globus pallidus, and periaqueductal gray areas which have been implicated in repetitive behaviors, and is also rescued by rapamycin (Fig. 7C). There were very few significant changes in FC in control mice following rapamycin treatment (Fig S3C) which aligns with our observation that there were no significant behavioral effects of rapamycin on control mice (Fig 3A-B, 4B-F, 5A-B).

We validated several of these imaging findings with immediate early gene (IE) immunostaining for Egr1+ neurons in the brains of mice euthanized 30 minutes after behaving freely in an open field environment where MIR mice displayed characteristic repetitive behaviors and control mice performed normal exploratory behaviors. Using Egr1 expression as a correlate of recent neuronal activity we found that the motor cortex and striatum of MIR mice had increased egr1 positive neurons, which also confirms our electrophysiology findings of neuron hyper-excitability in those brain regions (Fig S1D). We also found increased egr1+ neurons in MIR brains compared to controls in the retrosplenial cortex and decreased egr1+ neurons in the medial amygdala (Fig S1D).

Taken together, our studies of network connectivity indicate that a mechanism by which rapamycin may acutely affect brain function is through rapid alterations in functional activation and connectivity and normalization of network flexibility that is affected by mTOR inhibition. Somewhat surprisingly, the major alterations in MIR mice that were corrected by rapamycin were in areas thought to be related to sensory processing.

### Analysis of Gene Expression Alterations in MIR Mice and After Rapamycin Treatment

In order to identify additional potential mechanisms underlying MIR-induced behavioral changes and their mitigation with rapamycin, we performed single nucleus RNA sequencing (snRNA seq) of adult MIR and control mice and bulk RNA sequencing of MIR and control animals. SnRNA sequencing of tissue from the somatosensory cortex of adult MIR and control offspring did not indicate significant changes in the numbers of cells found in the different cell populations in this cortical brain region. Volcano plots visualized the most significantly differentially expressed neuronal (Fig 8A-C) genes in MIR mice relative to controls. Although there weren’t significant changes in the cellular makeup of the cortex, the number of differentially expressed genes (DEGs) within each cell type analyzed between MIR and Control offspring shows that there were many significant cell intrinsic changes with the greatest number of DEGs found in oligodendrocyte, microglia, upper layer and deep layer excitatory neuron populations, and other excitatory neurons (Fig 8D, p<0.05). Overall, there are more significant DEGs in the excitatory neurons compared to inhibitory neurons in MIR mice (Fig 8D). mTOR pathway-related genes are significantly enriched in MIR mice in oligodendrocytes and their precursor cells, microglia, PV interneurons, excitatory upper layer neurons, other excitatory (neurogranin) neurons, endothelial cells, and astrocytes (Fig 8E; p<0.05). A number of autism-associated, epilepsy/seizure-associated, and neuro-inflammatory-associated genes identified by cell type were significantly differentially expressed in MIR compared to control offspring using pseudobulk analysis of the snRNA seq data (LogFC>1, adjusted p value <0.05, FDR<0.1, Table 3). Two of these genes, Kirrel and GRIP1, that were significantly down-regulated in MIR mice are also known to have similarly reduced function in people and in other mouse models of sensory over-responsivity and repetitive behaviors^202–205^. Gene set enrichment analysis (GSEA) of snRNA seq data by pathway for all cell types indicated significant increases in genes for the energy-dependent regulation of mTOR and activity of kainate receptors, among others (Fig S3A) in MIR mice. GSEA of pathways by cell type shows (1) enriched gene expression involved in ion channel transport and homeostasis and decreases in TGF-beta and potassium channel signaling in all neurons in MIR mice (Fig S3B), (2) enriched gene expression involved in NF-kappa beta, Notch, and Toll-like receptors 5 and 8 and decreased expression of cytosolic calcium level and AP-2 transcription factors in excitatory MIR neurons (Fig S3C), and (3) enriched gene expression involved in GABA receptor activation and interleukin-1 signaling but decreased expression of pathways for voltage-gated potassium channels, NCAM-regulated neurite outgrowth, and gap junction regulation in inhibitory neurons from MIR mice (Fig S3D). These data indicate that the lack of significant cellular population changes in MIR mice indicates that the neuronal hyperexcitability found in these mice is more likely due to alterations in cell intrinsic properties because there is no loss or over-abundance of a particular inhibitory or excitatory neuronal population. Therefore, the mechanism of acute rapamycin treatment in rescuing the abnormal phenotypes in MIR mice is likely to be targeting intrinsic neurotransmission or ion channel phenotypes in the neural cell types with activated mTOR signaling which includes PV interneurons and upper layer excitatory neurons which can be rapidly altered (Fig 8E).

**Figure 8.**
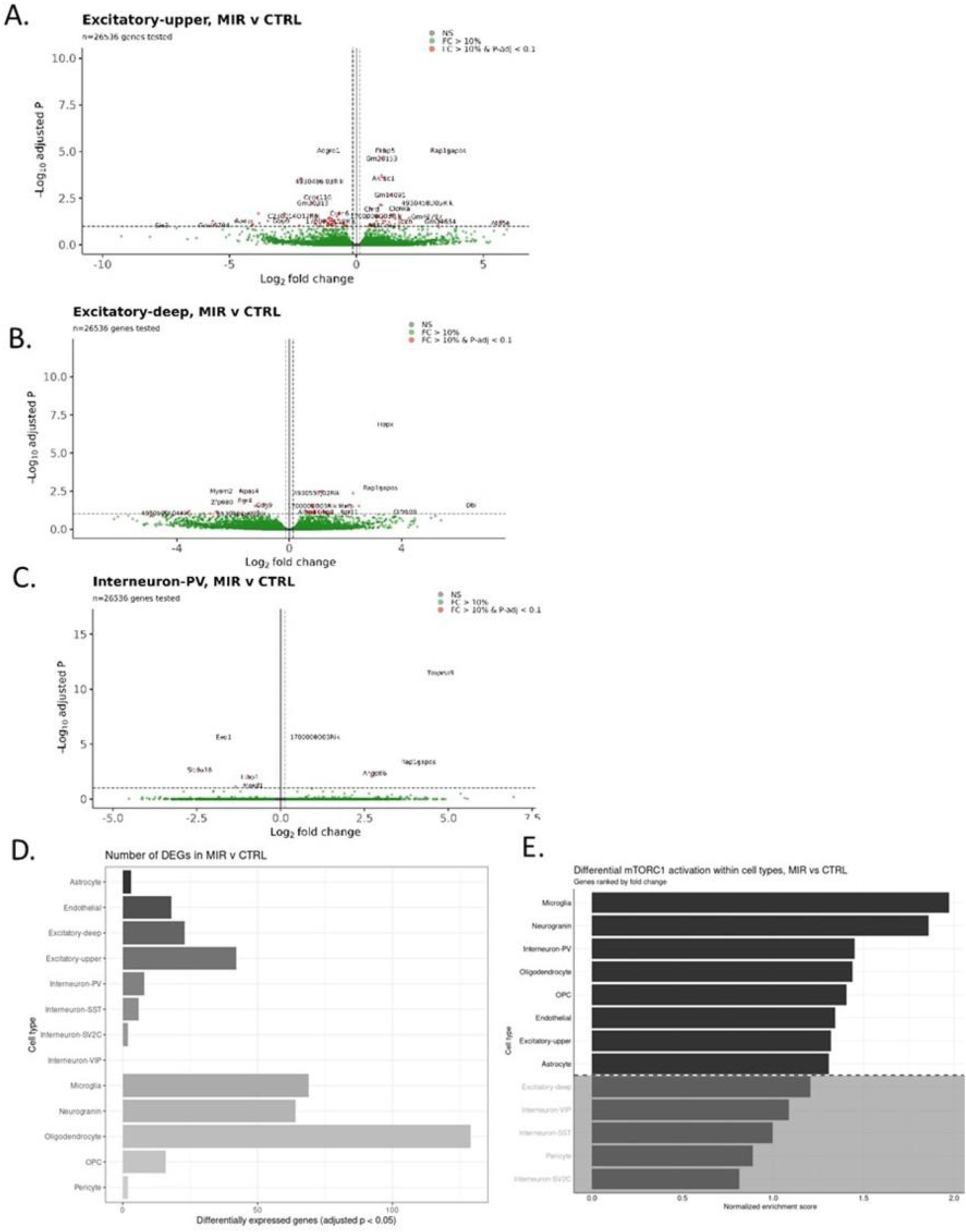
Cell sequencing from sensorimotor cortex microdissections from MIR and control adult offspring. Clustering of single-cell RNA seq data from sensorimotor cortex microdissections as volcano plots of differentially expressed genes in MIR offspring compared to Controls in **(A)** upper layer and **(B)** deep layer excitatory neurons and in (**C**) parvalbumin (PV) inhibitory interneurons; **(D)** The number of genes enriched in MIR mice by cell type; **(E)** Relative enrichment (fold change) of mTOR pathway regulated genes significantly enriched in MIR mice by cell type indicates that MIR mice have an increase in these genes in microglia, neurogranin neurons, PV+ interneurons, oligodendrocytes, oligodendrocyte precursor cells (OPC), endothelial cells, upper layer excitatory neurons, and astrocytes compared to control offspring. N=6/group

**Table 3.** Single nucleus sequencing analysis of significantly differentially expressed genes in MIR mice compared to control offspring are shown for Autism-associated genes, Epilepsy/Seizure-associated genes, and neuro-inflammation-associated genes; All data use (LogFC >1, P_val_adj <0.05, FDR<0.1) for abundance and significance criteria. *Sensory over-responsivity and repetitive behavior associated genes. N=6/group/treatment.

| Cell Type | ASD-associated Genes |  | Epilepsy/Seizure-associated Genes |  | Neuroinflammation-associated Genes |  |
| --- | --- | --- | --- | --- | --- | --- |
|  | Up MIR | Down MIR | Up MIR | Down MIR | Up MIR | Down MIR |
| Excitatory Deep Neurons | DBI HMGA2, CCND3, MAFB | NPAS4, MYOM2, EGR4, KIRREL* | EGR4, DBI, MAFB | NPAS4 |  |  |
| Excitatory Upper Neurons | FKBP5, CCND3, CD7 | NPAS4, MYOM2, EGR4, COL11A1 | FKBP5, EGR4 | NPAS4, ACTN2 |  |  |
| Neurogranin Neurons | TBR1 | ACTN2, MCM6, MRC2, DISP1, NOSTRIN, DISP1, KCTD18, KCNMB2, TTC28, NKAIN3, GPC6, CDK6, GRIP1*, EGFR, FILIP1, ST8SIA2 | TBR1, DISP1 | MCM6, DISP1, KCNMB2, NKAIN3, GRIP1*, ST8SIA2, EGFR, FILIP1 |  |  |
| Interneurons | WIPF1, SLC15A2, RPL4 | Exd1, SLC17A8 | WIPF1 |  |  | SLC17A8 |
| Microglia | FKBP5, E2F1, MCM6, C4B, PIK3C2G, FAM111A, DIAPH3, CCL12, HSPD1, TUBB5 | SEMA4B, SLC44A2 | FKBP5, FAM111A, DIAPH3 | SEMA4B | XDH, CCL12, C4B, TUBB5 |  |
| Astrocytes | IRS2 |  |  |  |  |  |
| Oligodendrocytes | HIF3A, FKBP5, C4B, CDH6, FGF1, DUSP1, BCAT1, KCNJ3 | LYPD6, SLPI | DUSP1, KCNMA1 |  | PHYHD1, C4B, MAT2a | SLPI |
| OPCs | HIF3A, FKBP5, NPAS1, HSP90AA1 |  | FKBP5, ARF6 |  |  |  |

Bulk RNA sequencing was performed on somatosensory cortical tissue from MIR and Control offspring with and without acute rapamycin treatment. Several DEGs that can rapidly affect neuronal function within the short time period of the acute rapamycin treatment were significantly altered in MIR mice including GABA and glutamate-related genes, transcription factors, and ion channel genes (LogFC>1, p<0.05, FDR<0.1, Table 4). Bulk sequencing data was also used to examine the expression of high-confidence ASD-associated risk factor genes in MIR and control offspring and in offspring treated with rapamycin or vehicle using published gene sets^68,96,206^. We found a small number of these ASD-associated genes were significantly up- or down-regulated in MIR mice, including TEK, PLAUR, ID1, Homer1, ASPM, TEKT4, CNTNAP3, Slc35D3, Grin2c, CHD7, CDKN1A (p21), and NeuroD1 (Table 5). Rapamycin rescued the over-expression of several ASD-associated genes, including TEK, Plaur, CDKN1A (p21), and ID1, and the under-expression of TEKT4, CHD7, and CNTNAP3 in MIR mice (Table 6). Rapamycin treatment also altered expression of several genes that were differentially expressed in MIR mice relative to controls that aren’t known to be specifically associated with autism, including decreasing Nestin, GEM, TIPARP, and APOLD1 expression and increasing HIF3A and PLIN4 expression (Table 6). GSEA analysis of pathways in these bulk sequencing experiments showed significantly enriched expression of genes involved in glutamate receptor signaling, oxytocin signaling, PI3K pathway signaling, apoptosis, and angiogenesis pathways in MIR mice (Fig S3E). Gene ontology for biological processes showed that MIR brains are enriched for G-protein coupled receptor signaling and negative regulation of hydrolase activity among others but decreased expression of genes involved in the regulation of cytokines, amine metabolic processes, and autonomic nervous system development in MIR mice (Fig S3F). Following rapamycin treatment, GPCR signaling, innate immune system, signal transduction, post-translational, and protein modification signaling were all increased (Supplemental Fig S3G). GSEA for pathways indicated significantly elevated gene sets involved in regulation of neurotransmitter levels, lipid regulation, and response to growth factors but decreases in those involved in epithelial and skeletal systems following acute rapamycin treatment (Fig S3H).

**Table 4.**
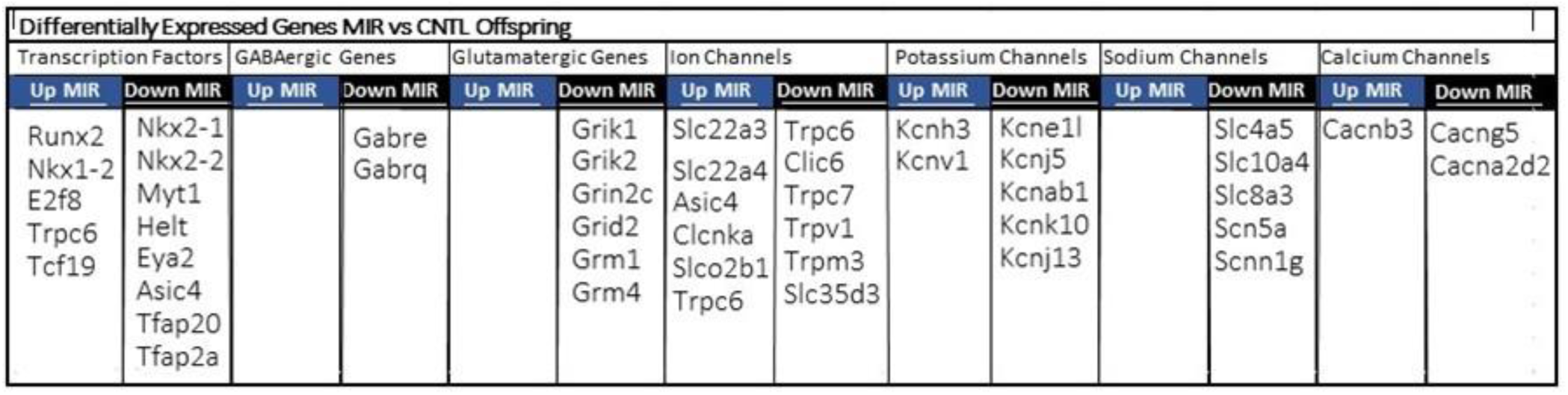
Bulk sequencing analysis of significantly differentially expressed genes in MIR mice compared to control offspring are shown for transcription factors, GABAergic genes, Glutamatergic genes, ion channel genes, potassium ion genes, sodium ion genes, and calcium ion genes which may affect neuronal excitability; All data use (LogFC >1, P<0.05, FDR<0.l) for abundance and significance criteria. N=12/group

**Table 5.**
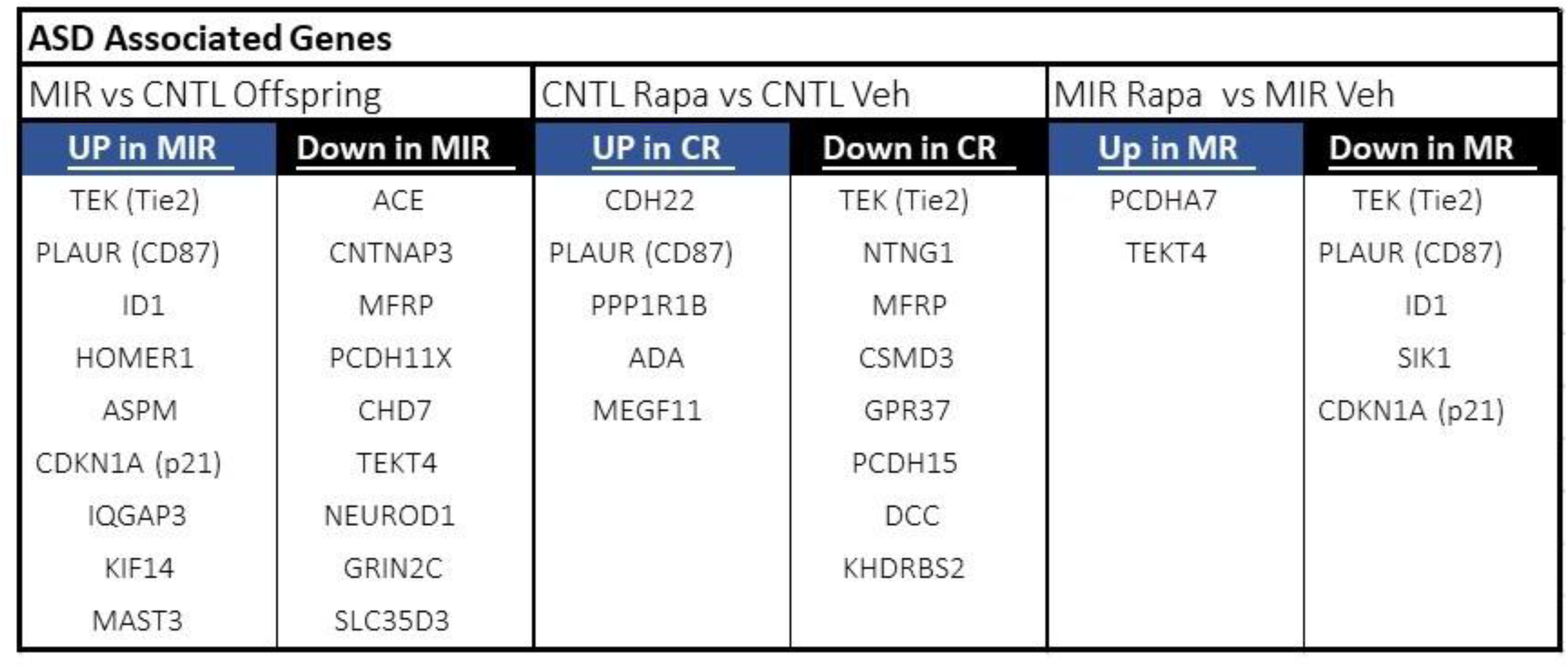
Bulk sequencing analysis of autism-associated genes that are significantly enriched in MIR mice relative to control offspring; All data use (LogFC >1, IP<0.05, FDR<0. l) for abundance and significance criteria. N=12/group

| ASD Associated Genes |  |  |  |  |  |
| --- | --- | --- | --- | --- | --- |
| MIR vs CNTL Offspring |  | CNTL Rapa vs CNTL Veh |  | MIR Rapa vs MIR Veh |  |
| UP in MIR | Down in MIR | UP in CR | Down in CR | Up in MR | Down in MR |
| TEK (Tie2) | ACE | CDH22 | TEK (Tie2) | PCDHA7 | TEK (Tie2) |
| PLAUR (CD87) | CNTNAP3 | PLAUR (CD87) | NTNG1 | TEKT4 | PLAUR (CD87) |
| ID1 | MFRP | PPP1R1B | MFRP |  | ID1 |
| HOMER1 | PCDH11X | ADA | CSMD3 |  | SIK1 |
| ASPM | CHD7 | MEGF11 | GPR37 |  | CDKN1A (p21) |
| CDKN1A (p21) | TEKT4 |  | PCDH15 |  |  |
| IQGAP3 | NEUROD1 |  | DCC |  |  |
| KIF14 | GRIN2C |  | KHDRBS2 |  |  |
| MAST3 | SLC35D3 |  |  |  |  |

**Table 6.** Bulk sequencing analysis of differentially expressed genes that were enriched in MIR offspring relative to control offspring that are rescued (reversed), by acute rapamycin treatment. All data use (LogFC >1, P<0.05, FDR<0.1) for abundance and significance criteria. N=12/group

| MIR v Control DEGs Reversed by Rapamycin |  |
| --- | --- |
| Decreased by Rapa | Increased by Rapa |
| TEK (Tie 2) | HIF3A |
| PLAUR (uPAR, CD87) | PLIN4 |
| NESTIN | TEKT4 |
| GEM | CHD7 |
| ID1 | CNTNAP3 |
| TIPARP |  |
| APOLD1 |  |
| CDKN1A (p21) |  |

Thus, we found a number of differentially expressed genes in MIR offspring which control cellular mechanisms like excitatory & inhibitory neurotransmitter production, transcription factors, and ion channel functions which could contribute to the pathophysiological neuron excitability and aberrant functional connectivity that we have observed in MIR mice and which can be rapidly altered by an acute rapamycin treatment. There were also a number of autism-associated genes that were differentially regulated in MIR offspring that are consistent with the behavioral phenotypes of these mice. Although changes at the level of gene expression within a 2-hour time frame of our rapamycin treatment might not be expected to be large, we did see the rescue of several of the autism-associated genes that are differentially expressed in MIR mice, identifying a number of promising new therapeutic targets.

## Discussion

Exposure to mild maternal inflammatory response (MIR) in early gestation in CD-1 mice recapitulated multiple ASD associated physiological and behavioral phenotypes. This included mild brain overgrowth in juvenile ages followed by regionally variable volumetric brain changes in later adulthood like cortical thinning and larger hippocampal, basal ganglia, and ventricular structures. Similar growth trajectories have been reported in clinical ASD and neuro-inflammation studies^188–194,207^. MIR mice also demonstrated other common ASD pathology like chronically elevated blood cytokines and brain microglia which have also been reported in human ASD populations^19,83–97,208–213^. Furthermore, this model produces offspring with multiple ASD-related behavioral phenotypes like social, communication (vocalization), anxiety, and repetitive behaviors^118^. We have now extended the findings to show that MIR mice demonstrate significant sensory over-responsivity (SOR) and sensory processing dysfunction as well. SOR is an extremely impairing clinical condition marked by avoidance and/or sensitivity to sensations such as scratchy clothing, loud noises, or visually stimulating environments. SOR is present at high rates across the autism spectrum and is associated with other core features of ASD, as well as difficulties with behavioral and emotion regulation^214–217^. Despite its significance, very little is known about the causes or underlying biological mechanisms of SOR and there are essentially no evidence-based treatments for SOR. Several genetic murine models of mutations in autism risk factor genes like Mecp2 (Rett Syndrome) and FMR1 (Fragile X)^159,199^ are known to display SOR behaviors but ours is the first report of SOR in an acquired model of ASD with a neuro-inflammatory origin.

Although it may be surprising that a single exposure to low-level maternal inflammation, a level that does not cause sickness behavior in the pregnant dams or reduced litter sizes, could cause such lasting effects on the brain and behavior, we and others have found that early *in utero* exposure sets up chronic dysregulation in systemic immune function^218–223^. Our analysis of both young and older adult MIR offspring showed a persistent elevation in pro-inflammatory cytokines in the blood and increased brain-wide increases in microglia, suggesting a chronic inflammatory state across the life-span compared to age-matched controls. Both clinical and pre-clinical studies have previously reported some success in the clinical treatment of ASD in children with an anti-inflammatory drug, pioglitazone, which improved behavioral phenotypes along with reducing neuro-inflammation^224–232^. We also found a concomitant chronic elevation of mTOR signaling in the neurogenic adult subventricular niche^118^ and now in other regions in adult (cortex, amygdala, and striatum). Induction of systemic and brain inflammation in adult wildtype/control mice does not produce any of the ASD-associated abnormal behaviors that are produced in MIR mice where inflammation was induced *in utero* and persists chronically^233^. Therefore, an elevated inflammatory state in the adult brain does not, on its own, explain most of the MIR phenotypes, suggesting that there are likely important vulnerable periods in brain development where inflammatory exposure is especially deleterious and/or the effects of chronic inflammation and increased mTOR pathway activity have stronger effects. It is well known that chronic neuro-inflammation can contribute to significant brain pathophysiology^222^. Maternal inflammation is also known to be a significant risk factor for both ASD and brain overgrowth^19,85,101–117^. Therefore, one of our initial hypotheses was that the chronically increased brain microglia underlies some or all of the abnormal brain pathology and behaviors in the MIR model as microglia have been demonstrated to influence several related behaviors and mTOR signaling^90,234–240^. However, our experiments which ablated microglia in the adult brain suggest that the role of microglia in the etiology of MIR-induced abnormalities is complex. Firstly, microglia ablation with a CSF1 inhibitor can rescue some abnormal behaviors in young adult mice but not in older adult mice suggesting that they likely play a role in setting up physical and functional brain changes but are not the primary cause of behavioral dysfunction since acute treatment with the mTOR inhibitor, rapamycin, can significantly reduce abnormal MIR phenotypes in both young and old MIR offspring. Thus, the effects of MIR exposure on behavior are complex and evolving over the lifespan as we can see in the physical changes to the MIR brain such as the early overgrowth followed by later regional decrease in cortical brain volume. Chronic post-natal neuro-inflammation and mTOR activation in the brain has also been associated with epilepsy and a disturbed excitatory/inhibitory balance^207,241–243^. In keeping with a putative neuro-inflammatory etiology, our MIR mice also display significant cortical hyper-excitability.

Given the long-term elevation in mTOR signaling that we have seen in MIR offspring and the similarity of some ASD-associated phenotypes in genetic models of autism (TSC and PTEN mutants) that also have increased mTOR pathway activation, we examined the role of mTOR signaling in MIR pathology. It is known that the mTOR inhibitor, rapamycin, can rescue several abnormal ASD-associated behaviors in murine genetic models of PTEN and TSC mutations even in adult mice with existing/ongoing brain physiological abnormalities like brain overgrowth^9,19,124,125,131,132^. Therefore, we hypothesized a similar rescue effect was possible in our MIR model. However, we observed an unanticipated and potentially useful rapid effect of rapamycin on several abnormal behaviors in MIR offspring. We confirmed via western blot analysis that the rapamycin treatment was able to access the CNS and significantly reduce mTOR signaling within the brain during this acute time course. Additionally, it has been reported that sensory abnormalities in Mecp2 and FMR1 mutant mice can be reversed through entirely peripheral neuron interventions^159,199^. Therefore, to test this in our maternal inflammation model we used a newly developed FKBP12 antagonist, RapaBlock, which prevents rapamycin effects in the peripheral nervous system to demonstrate a CNS-mediated effect on MIR mice. These insights allowed us to focus our search for the causes of MIR abnormalities and rapamycin effects in the brain on rapidly changeable physiological and functional mechanisms which would not be apparent from more chronic rapamycin treatment paradigms that have been shown to induce permanent physical transformations such as restoring normal dendritic and synaptic complexity^9,22,120–124^. In this way, studying the rapid effects of rapamycin may reveal novel biological and functional mechanisms underlying multiple ASD-associated abnormal phenotypes that can be overlooked with chronic treatment paradigms.

While synaptic strengthening/weakening through altered levels of neurotransmitter release or post-synaptic receptor expression can happen on short (minutes to hours) time scales^244–246^, widespread physical synaptic remodeling is an unlikely mechanism for our acute rapamycin effects. This led us to examine changes in neuronal gene expression, excitation, and functional activation and organization at the circuit level as possible fast-acting mechanisms that could underlie the brain and behavioral dysfunction rescued by acute rapamycin treatment in the MIR model. Our single cell sequencing results did not find that MIR mice have significant neuronal or glial cell population size changes. However, there was significantly upregulated gene expression related to mTOR pathway activation in multiple cell types including microglia, astrocytes, oligodendrocytes and their progenitors, excitatory upper layer neurons, and inhibitory parvalbumin neurons in MIR mice. This suggests that cell-intrinsic changes are responsible for mTOR dysregulation in MIR offspring rather than alterations in the balance of specific cell population numbers. Additionally, we found that a number of autism, epilepsy, and neuro-inflammation associated genes alterations were differentially expressed in upper and deep layer excitatory neurons from MIR offspring that have been associated with repetitive behaviors, sensory processing dysfunction, synaptic plasticity, and epilepsy. Bulk sequencing with and without rapamycin treatment suggests several gene expression alterations occur in several rapidly changeable cellular properties such as transcription factors, ion channels, and neurotransmitter synthesis-related genes in MIR mice. In addition, several high confidence ASD risk genes^68,96,206^ were altered in MIR offspring and a number of these were reversed by acute rapamycin treatment. Several of these ASD genes also have associations with alterations in brain size (ASPM, KIF14), epilepsy (PLAUR, ASPM, MAST3, HOMER1, ACE, ID1, CHD7, CDKN1A (p21)), intellectual disability (CHD7, TEKT4), sensory processing (PLAUR), synaptic plasticity (HOMER1, CNTNAP3, IQGAP3) and neuroinflammation (TEK/Tie2, CNTNAP3, ACE, CDKN1A^247–275^. Although the single cell sequencing results showed that there were no significant changes in the relative balance of excitatory and inhibitory neurons in MIR mice, we did observe a significant hyper-excitability in cortical neurons in our electrophysiology experiments, which could be related to the cell-intrinsic gene expression changes that we saw in excitatory and inhibitory neurotransmitter- and ion channel-related genes in MIR neurons. Rapid reduction of this neuronal hyper-excitability and seizure susceptibility with rapamycin treatment within the same acute time-frame as behavioral rescue suggests that restoring excitatory-inhibitory regulation in the brain may underlie the normalization of sensory dysfunction and ASD-related behaviors in the MIR model. In support of this, others have shown that an acute 2-hour pre-treatment with the rapalogue everolimus also reduces seizure susceptibility and suppressed neuro-inflammatory responses in a kainic acid rat epilepsy model^276^. A recent clinical trial (EXIST-3) of everolimus for epilepsy in patients with tuberous sclerosis complex (TSC) found moderate success with chronic treatment improving seizure status in around 60% of subjects that lasted at least 48 weeks^277^. However, the clinical evidence to date for everolimus effectiveness for treating other autism phenotypes like neurocognitive function or social behavioral skills is mixed with some studies finding no positive effects and others finding small improvements in autism symptoms^138,139,278^. Our findings of tachyphylaxis in animals chronically treated with rapamycin could also help explain why daily rapalog treatment may not have a continued, robust effect on seizure frequency in mTORopathies.

Functional connectivity (FC) is another level of brain regulation which can be rapidly altered by acute rapamycin treatment. The majority of the FC changes in MIR offspring compared to controls in our resting-state fMRI data were increases in FC and only a few brain regions showed decreases. We found that the largest proportion of FC differences between MIR and control offspring involved largely sensory rather than motor circuits despite the strong repetitive motor phenotype in MIR mice. While there was an increase in M1 motor cortex and basal ganglia FC in MIR brains the majority of the hyper-connectivity in cortico-striatal cortico-limbic functional connections were from sensory cortical areas. Additionally, the basal ganglia regions with significantly increased FC connections to motor cortex in MIR were the dorsolateral striatum and external GP which are known to have sensory processing roles in addition to motor functions^279–282^. The most significant FC changes in MIR mice relative to controls were found in the thalamus which prompted us to do a seed analysis from this region. Thalamic sensory FC was altered in multiple primary sensory cortical areas as well as sensory salience network structures (insula and anterior cingulate) in MIR brains. We interpret this dominant sensory phenotype in FC alterations to be in keeping with the behavioral SOR findings in the MIR model. However, it does not entirely fit with the strong repetitive behavior phenotype even though there is considerable comorbidity for sensory dysregulation with repetitive behaviors^283–287^. We propose that recent clinical findings may offer an explanation for the lack of a stronger functional connectivity motor phenotype in our mice. A recent human study showed that reductions in repetitive behaviors in individuals are associated with an increase in other autism phenotypes like anxiety and social difficulties and vice versa, suggesting that some repetitive behaviors may be a self-regulation mechanism rather than a primary phenotype in autism^288^. Given the close correlations that have been found between repetitive behaviors and sensory sensitivity and the beneficial effects of repetitive behaviors^289^, our finding that sensory dysregulation is the more dominant brain phenotype in rsfMRI of MIR mice with SOR, supports the idea of an emergence of repetitive behaviors as self-soothing or self-regulating mechanism rather than being a primary dysfunction. The acute rapamycin treatment in MIR mice reduced FC towards control levels in the cortex but curiously caused even greater FC differences in subcortical structures. This shows that the rescue effects of the treatment are not entirely via a restoration back to control levels and patterns of functional connectivity across the brain. This is likely related to the fact that brain structural abnormalities remain, such as the increases in hippocampus, caudate, and amygdala size and decreased thalamus and cortical volumes and rapamycin is acting in the context of these physical changes. Longer treatments with rapamycin like those in Pagani et al.^22^ in TSC mutant mice show that rescuing the physical synaptic abnormalities produces a greater return to control FC patterns. These circuit-level insights into mechanisms of dysregulation and the pattern of changes that are associated with behavioral recovery have identified potentially novel neuromodulatory targets for non-pharmaceutical based treatments like TMS and focused ultrasound.

This sensory-dominant, hyper-connectivity profile closely aligns with recent large-scale cross-species fMRI analyses^290^which identified a reproducible autism-related hyper-connectivity subtype linked to immune, transcriptional, and mTOR-related signaling pathways, in contrast to a hypoconnectivity subtype associated primarily with synaptic dysfunction. Notably, mTOR signaling emerged as one of the few synaptic-related pathways enriched within the hyper-connectivity subtype, positioning mTOR as a key molecular bridge between immune activation and functional network alterations. This is particularly relevant to our MIR model, in which maternal inflammation induces persistent upregulation of mTOR signaling in offspring. Thus, elevated mTOR activity provides a plausible mechanism for how chronic immune activation could bias developing circuits toward increased excitatory drive, altered synaptic maturation, and aberrant functional coupling. The prominent involvement of thalamic, sensory, and salience networks in MIR mice FC alterations mirrors the hyper-connectivity signatures reported across immune- and mTOR-linked mouse models in the Gozzi lab dataset.

Analysis of network community architecture is a data-driven approach to evaluate brain wide synchrony and network properties rather than using a strictly anatomical analysis of functional connectivity that relies on predefined brain regions. These circuit-level properties can be rapidly modulated and significantly affect brain function. Graph analysis of this functional circuitry allows the unbiased detection of communities of brain regions with strong functional connectivity, which may represent specialized functional units or systems in the brain. Graph theory can also address cognitive flexibility and information integration by examining how hub nodes (highly connected nodes within a module that maintain network integrity and facilitate information flow between brain regions) are structured, the strength of this connectivity, and the size (specialization) of modules. This network analysis of resting-state fMRI revealed that the MIR offspring have a larger number of modules compared to control offspring, suggesting that their brain network is more segregated into distinct functional sub-networks within the subcortical regions, representing an increased specialization of function. However, the nodes within these subcortical modules also have a significantly shifted higher classification diversity values than in controls, indicating nodes more frequently transition membership between modules within the MIR group, reflecting potentially increased information transfer between modules, and greater functional integration within the brain. However, the opposite is true within the cortex of MIR mice, with a lower classification index indicating reduced information transfer to other network modules of the brain. The combination of increased module number and both higher and lower intra-module nodal diversity suggests a complex reorganization of brain networks which may reflect both deficits and compensatory adaptive changes in functional connectivity. We are unaware of other resting state fMRI studies in preclinical models like ours that have examined this type of global functional reorganization at the network level. However, there have been clinical studies that have undertaken this type of network analysis that have similar findings of increased numbers of functional modules, increased connections between modules, and increased module diversity in individuals with ASD with resting-state fMRI^291–296^. Our MIR offspring share this aberrant network architecture with humans, demonstrating another cross-species phenotype that is present in our inflammatory, acquired model of ASD. It has been suggested that the functional modular reorganization in the ASD brain may be a compensatory mechanism for restoring functional homeostasis at the circuit level in response to physical connectivity changes, and that therapeutic normalization of this altered modular organization would therefore be likely to worsen ASD dysfunction^292^. However, our acute rapamycin treatment data suggest otherwise. The rapid rescue of this abnormal network modularity in MIR mice following rapamycin treatment was one of the largest effects of the pharmacological treatment on the MIR model of brain dysfunction. Both the number of modules and the distribution of the nodal connectivity within these modules was returned to control levels after acute rapamycin, suggesting that at a macro circuit level, the treatment restores functional information flow and network stability within the same time frame that we observe behavioral and electrophysiological rescue from this treatment. Therefore, our findings strongly suggest that modular organization of functional brain networks are related to behavioral traits in ASD and that this may be a central functional mechanism underlying inflammation-mediated behavioral dysfunction in several core and comorbid autism phenotypes. There is evidence in the literature that supports the theory that these functional brain alterations that we have observed in the MIR model are induced by maternal inflammation and sustained by the chronic inflammation in the offspring. For example, neuro-inflammation has been increasingly recognized as a factor contributing to atypical functional connectivity patterns and network abnormalities^297–303^. Neuro-inflammation can affect synaptic function by modulating the release of neurotransmitters, altering receptor expression, and ion channel function which impacts long-term potentiation and long-term depression^72,86,304–309^. These changes can disrupt the formation and maintenance of functional connections leading to functional network abnormalities. For example, enhanced long-term depression in humans and animal ASD models have been shown to underlie hyper-reactive sensory processing^310^. Neuro-inflammation during critical periods of brain development can also interfere with processes like axonal growth and synaptogenesis^87,227,311–315^ which can contribute to the increased modularity and module diversity seen in ASD and in our MIR model as the brain adapts to these physical changes with functional compensation towards the goal of homeostasis. Disruption to normal glutamate and GABAergic neurotransmission and receptor expression can also cause imbalances in neuronal excitation thresholds that further impacts functional network^207,243,316–321^ which is consistent with the phenotypes that we have observed in MIR offspring.

In summary, our experiments demonstrate a plausible etiology for MIR-induced brain and behavioral dysfunction that is revealed through the acute effects of rapamycin: (1) exposure to MIR *in utero* causes neuroinflammation and chronic post-natal neuroinflammation in the offspring that, (2) elevates mTOR signaling chronically, (3) increases neurogenesis and synaptogenesis at a critical period of early brain development leading to, (4) brain overgrowth, abnormal physical connectivity, and (5) a compensatory functional reorganization of the brain connectome. Our data also suggest that exposure to maternal inflammation could account for the non-macrocephalic brain overgrowth in ASD that has been identified in early childhood^98–100^ and that even mild alterations in brain growth can produce aberrant functional connectivity and cellular dysfunction that leads to brain-wide functional network changes and behavioral abnormalities. Although we have shown that aberrant brain and behavioral phenotypes can be understood as dysfunction at the level of large-scale networks and their information integration, we have also established in this MIR model and with acute rapamycin treatment that this global activity is dependent on processes happening at the level of physical systems like the immune system, of cells and their excitability, and of sub-cellular processes like pathway activation and gene expression regulation. Therefore, although longer interventions in suppressing excessive mTOR signaling have been shown to also impact the physical features of maladaptive plasticity^22^, we have identified a number of potential therapeutic targets at various levels of dysfunction. We demonstrate that the MIR brain is amenable to rapid normalization of functional organization and neuronal excitability with short rapamycin treatments which results in rescue of core and comorbid ASD behaviors in adult animals without requiring long-term physical alterations to the brain. Thus, restoring excitatory/inhibitory imbalance and sensory functional network modularity may be key mTOR-related targets for therapeutically addressing both primary sensory phenotypes and compensatory repetitive behavior phenotypes.

## Acknowledgements

Funding: Dr. Miriam and Sheldon G. Adelson Medical Research Foundation (HIK, AS, RK, DHG)

UCLA Intellectual and Developmental Disabilities Research Center NIH grant HD103557 (HIK, DHG, RK, JL, KT)

Center for Autism Research and Treatment (CART) Pilot Grant Award, which is supported by NIH/NICHD grant P50-HD-055784 (JL)

UCLA Brain Injury Research Center (NH, JL)

Simons Foundation Cross-Species Studies of ASD Grant SFI-AN-AR-Cross Species-00005172 (JL, NH, HIK)

UCLA Clinical and Translational Science Institute (CTSI) Accelerator Core Voucher Program supported by grant number UL1TR001881(HIK)

Autism Speaks Basic and Clinical grant (JL, HIK)

Autism Speaks Environmental Sciences grant (JL)

Sequencing was done by the UCLA Technology Center for Genomics and Bioinformatics, Dr. Xinmin Li Director and supported by the Jonsson Comprehensive Cancer Center (P30 CA016042)

Z.Z. is a Damon Runyon Fellow supported by the Damon Runyon Cancer Research Foundation (DRG-2281-17)

Electrophysiology experiments were supported by the Cell, Circuits and Systems Analysis Core (NIH P50HD103557)

KT was supported by NIH F31 MH122205

---

**Supplemental Figure 1.**
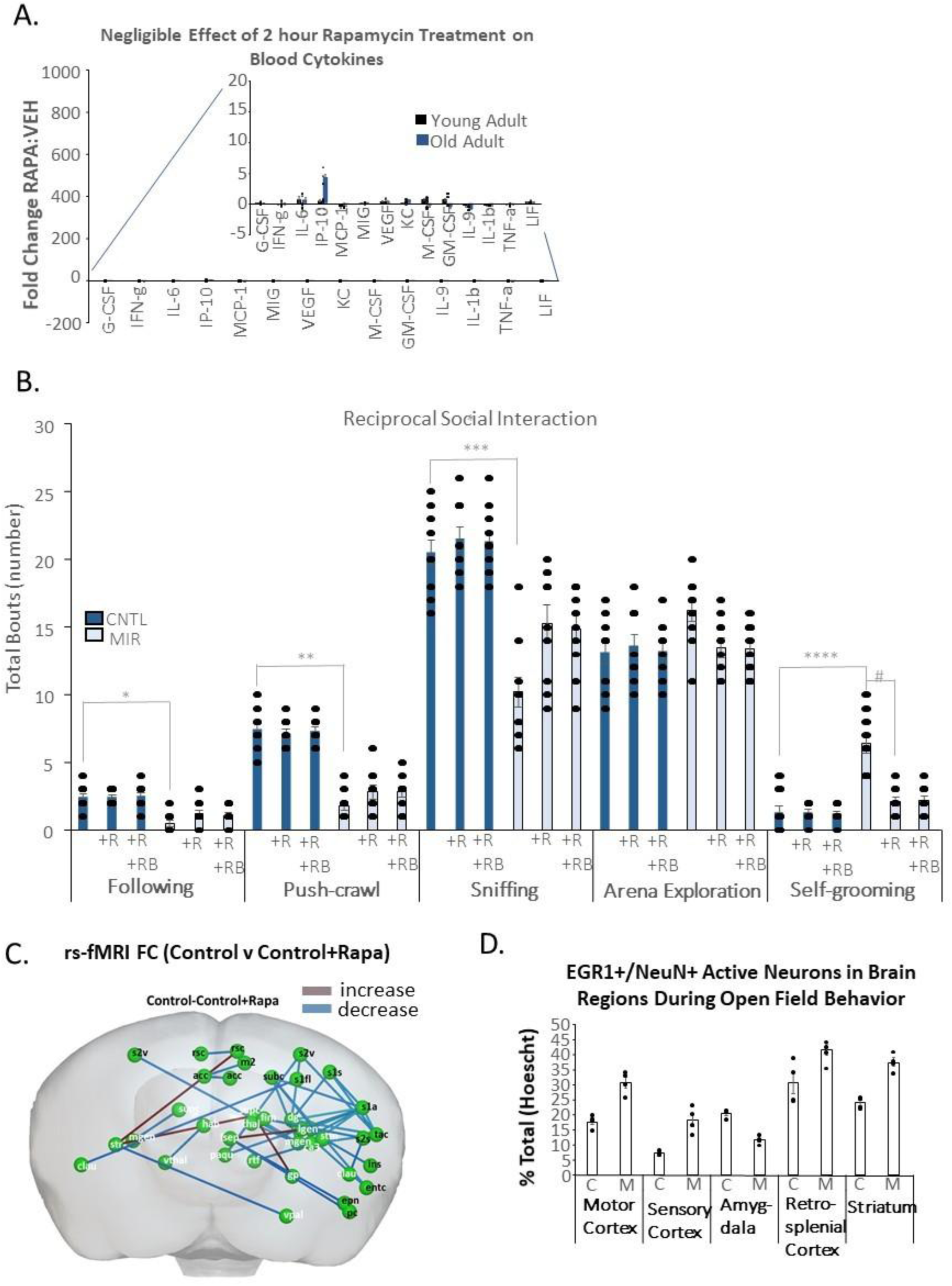
Acute Rapamycin Doesn’t Decrease Inflammatory Cytokines in MIR Mice, MIR mice have deficits in some reciprocal social interaction behaviors that is improved by acute rapamycin and is not affected by RapaBlock, Rapamycin·induced Functional Connectivity Changes in Resting State fMRI in Control Offspring, and lmmunohistochemistry Validation of Increased Neuron Activity in Brain Regions Indicated by fMRI Results. **(A)** Blood cytokines are unchanged by 2-hour Rapamycin treatment in adult MIR mice (all n.s.), see Figure 1 legend for cytokine abbreviations and full names, N=4/group/treatement; data shown as mean +/·SEM; **(B)** A two-way ANOVA analysis for multiple comparisons of social approach behaviors in MIR and control offspring treated with rapamycin or vehicle control in combination with RapaBlock or RapaBlock-vehicle found significant behavior x group (p<0.0001, F(ll, 115)), behavior (p<0.0001, F(2, 115)) and group (p<0.0001, F(5, 54)) effects. Post-hoc analysis (Tukey’s) shows significant deficits in MIR mice treated with vehicle-rapamycin and vehicle-RapaBlock compared to control offspring treated with vehicle-rapamycin and vehicle-RapaBlock for following behavior (adjusted *p=0.0005, DF=18), push-crawl behavior (**p<0.0001, DF=16), and sniffing behavior (adjusted ***p<0.0001, DF=17). There was no significant difference between MIR offspring receiving rapamycin and no RapaBlock compared to MIR offspring receiving rapamycin and RapaBlock for following behavior (adjusted #p=0.9998, DF=17), push-crawl behavior (adjusted p>0.9999, DF=17), and sniffing behavior (adjusted p=0.9999, DF=17). There was a similar but opposite effect with self-grooming during the test with MIR offspring treated with rapamycin-vehicle and RapaBlock-vehicle had significantly increased behavior compared to control offspring treated with rapamycin-vehicle and RapaBlock-vehicle (adjusted ****p=0.0001, DF=16), a significant difference between MIR offspring treated with rapamycin-vehicle and RapaBlock-vehicle and MIR mice treated with rapamycin and RapaBlock-vehicle (adjusted p=0.0007, DF=13) but there was no significant difference in grooming for MIR offspring treated with rapamycin and RapaBock-vehicle and rapamycin and RapaBlock (adjusted p>0.9999, DF=17), demonstrating no RapaBlock effect on rapamycin treatment which suggests that the effects of rapamycin are not via the peripheral nervous system, N=8/group/treatment; data shown as mean+/· SEM (**C**) The changes in FC in control mice after acute rapamycin treatment were small compared to MIR mice (Fig. 6C) and were mostly decreases (blue lines) between cortical and subcortical regions with few increases (red lines) between retrosplenial cortex and basal ganglia structures; N=15/group; (**D**) A two-way ANOVA analysis for multiple comparisons of the number of NeuN+ neurons that express the immediate early gene protein EGFl (as a percentage of total Hoescht+ cells) shows significant overall effects in brain region x group (p=0.0005, F_(2, 12)_), brain region (p<0.0001, F_(2, 12)_), and group (p=0.0003, F_(l, 6)_). Post-hoc analysis (Tukey’s) shows that there are significant increases in these cells in the brains from MIR mice compared to control offspring in the motor cortex (adjusted p=0.0024, OF =5), the sensory-motor cortex (adjusted p=0.0126, DF=3), and striatum (adjusted p=0.0005, DF=5) and a significant decrease in the amygdala (adjusted p<0.0001, DF=6). There was no significant difference in the retrosplenial cortex (adjusted p=0.0587, DF=5), N=4/group; data shown as mean +/· SEM.

**Supplemental Figure 2.**
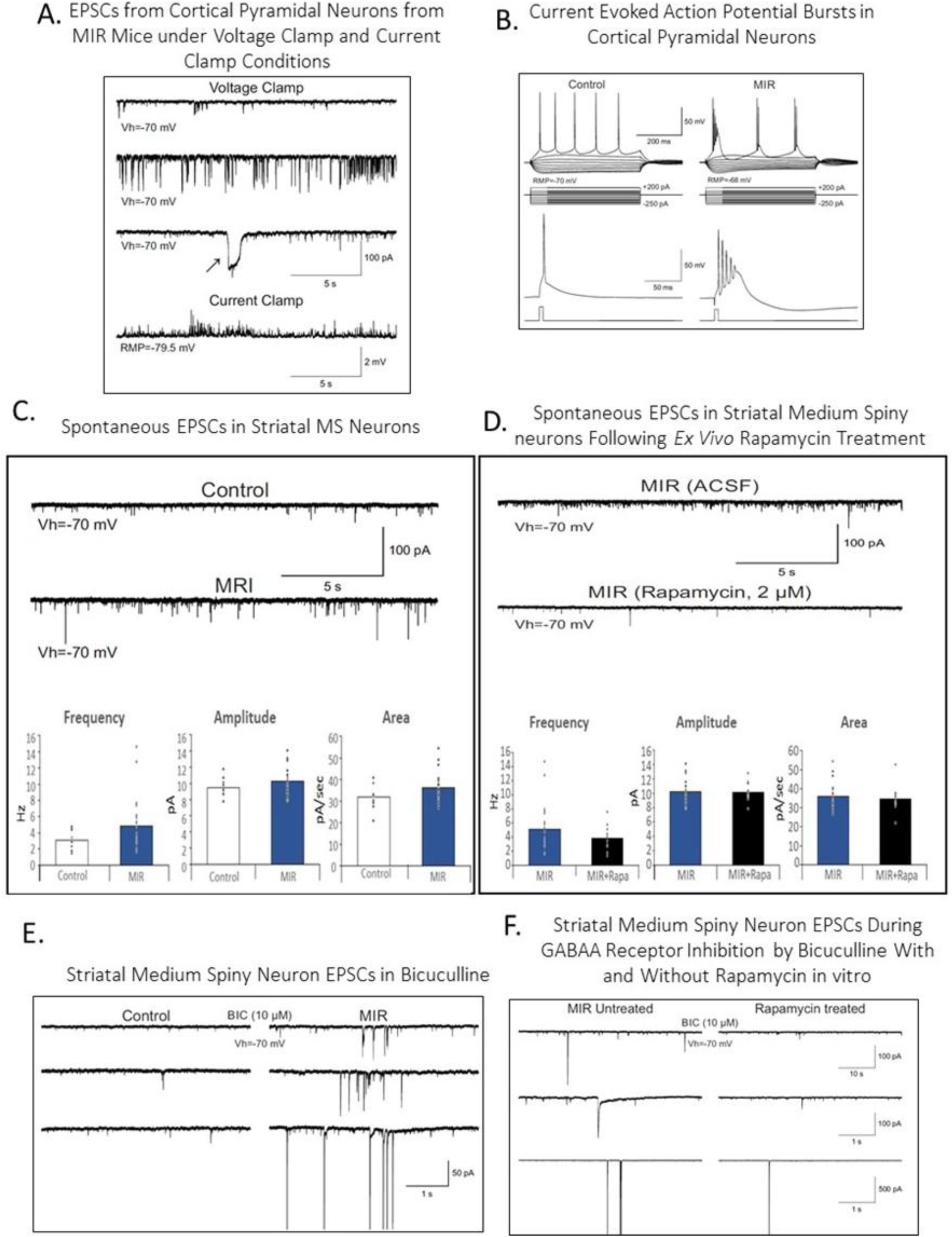
Cortical Somatosensory Pyramidal Neurons Display Multiple Signs of Hyper-Excitability in MIR Brains and Medium Spiny Neurons in the Striatum Show Trends Towards Hyper-excitability that is Reduced by Acute Rapamycin Treatment. **(A)** Example of excitatory post-synaptic current activity in cortical pyramidal neurons under voltage clamp and current clamp conditions; **(B)** Example of action potential bursting from cortical pyramidal neurons under current clamp conditions; (**C**) Traces and quantification of the frequency, amplitude, and area of spontaneous excitatory postsynaptic currents in striatal medium spiny neurons from adult MIR and control offspring show a trend for increases in MIR mice compared to control but do not reach significance; N=24 neurons/group;**(D)** Traces and quantification of the frequency, amplitude, and area of spontaneous excitatory postsynaptic currents in striatal medium spiny neurons from adult MIR mice treated acutely with rapamycin ex vivo (2uM) compared to controls shows a trend for decreased spiking frequency that does not reach significance; N=24 neurons/group **(E)** Traces showing treatment with Bicuculline, a competitive antagonist of GABA A receptors, increases excitatory post-synaptic currents in medium spiny neurons in the striatum, **(F)** Traces showing acute *ex vivo* rapamycin treatment given simultaneously with Bicuculline prevents the increase in medium spiny neurons EPSCs; The following abbreviations were used: RMP: resting membrane potential, Hz: hertz, pA: picoamperes, mV: millivolts, s: sec: seconds, BIC: Bicuculline; ACSF: artificial cerebrospinal fluid, Vh: holding potential

**Supplemental Figure 3.**
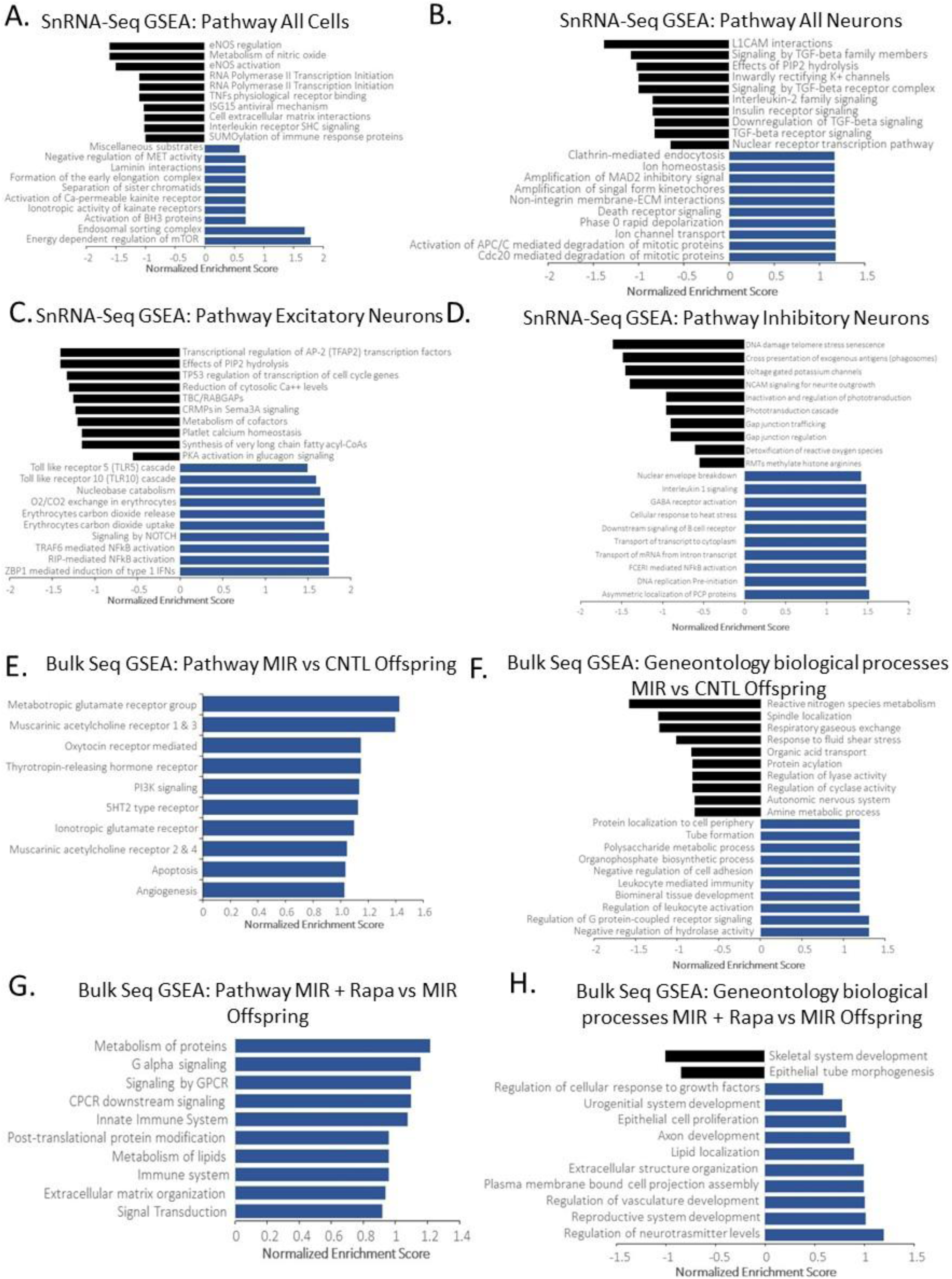
Gene Set Enrichment Analysis of snRNA seq and Bulk sequencing of Genes that Are Significantly Differentially Expressed in MIR Offspring. (**A**) Gene set enrichment analysis (GSEA) of pseudo-bulk expression from single-nucleus RNA sequencing in cellular pathways for all cell types in MIR mice relative to control offspring; (B) GSEA of pathways enriched in MIR mice in all neurons; (C) GSEA of pathways enriched in MIR mice in excitatory neurons; (D) GSEA of pathways enriched in MIR mice in inhibitory neurons; (E) Gene set enrichment analysis (GSEA) of pathways enriched in MIR mice compared to control offspring from bulk sequencing; (F) Geneontology of biological processes enriched in MIR mice compared to control offspring; (G) GSEA of pathways differentially expressed in MIR mice treated with rapamycin compared to vehicle treated MIR mice; (H) Geneontology of biological processes differentially expressed in MIR mice treated with rapamycin compared to vehicle-treated MIR mice; expression and significance criteria used for all are LogFC>l, P≤0.05, FDR ≤0.1, N=6/group/treatement

**Supplemental Table 1.** Single nucleus sequencing comparison of cell populations between MIR and Control offspring sensory-motor cortex indicates that there are no significant differences between treatment groups for any cell type. N=6/group

|  | Rank | logFC_mir_vs_ctrl | pVal_mir_vs_ctrl | FDR_pVal_mir_vs_ctrl |
| --- | --- | --- | --- | --- |
| Microglia_1 | 6 | 0.024198538 | 0.730088556 | 0.988144314 |
| Microglia_2 | 15 | 0.001630415 | 0.981174139 | 0.988144314 |
| Endothelial | 13 | 0.007223573 | 0.898156918 | 0.988144314 |
| Pericyte | 14 | -0.036032657 | 0.575069239 | 0.988144314 |
| Unclassified | 12 | 0.066055026 | 0.406867978 | 0.988144314 |
| Oligodendrocyte | 3 | -0.045686542 | 0.658849051 | 0.988144314 |
| Astrocyte_1 | 4 | -0.053901143 | 0.656765516 | 0.988144314 |
| Astrocyte_2 | 17 | 0.01232224 | 0.848949092 | 0.988144314 |
| OPC_1 | 8 | -0.011208874 | 0.86612819 | 0.988144314 |
| OPC_2 | 16 | -0.000997818 | 0.988144314 | 0.988144314 |
| Interneuron-SST | 7 | -0.012737802 | 0.870655188 | 0.988144314 |
| Interneuron-PV | 5 | 0.057783538 | 0.479172194 | 0.988144314 |
| Interneuron-VIP | 10 | 0.125438347 | 0.120224361 | 0.841010574 |
| Interneuron-SV2C | 11 | 0.069811105 | 0.365002986 | 0.988144314 |
| Neurogranin | 9 | 0.362809282 | 0.148413631 | 0.841010574 |
| Excitatory-deep | 2 | -0.497502762 | 0.000931602 | 0.01583724 |
| Excitatory-upper | 1 | 0.114508117 | 0.457762268 | 0.988144314 |

**Supplemental Table 2.** Litters Generated for Experiments. Fifty-eight independent litters were generated over several years for the experiments performed in this manuscript. The experiments and the figures the mice contributed to by group (MIH or CNTL) are indicated. Abbreviations are as follows: MIR= Maternal Inflammatory Response Offspring, MRI = Magnetic Resonance Imaging, e-Phys = electrophsilogical CNTL = Vehicle Control Offspring, Rapa = Rapamycin, PPI. = pre-pulse inhibition test, APO = apocynin; PTZ = pentylenetetrazol, IHC = immunohistochemistry.

| Litter | Group | Litter Testing | Figure | Table |
| --- | --- | --- | --- | --- |
| 1 | CNTL | Blood Cytokines Old/Microglia IHC +/- Rapa | 1B, 1D |  |
| 2 | CNTL | Blood Cytokines Old/Microglia IHC +/- Rapa | 1B, 1D |  |
| 3 | MIR | Blood Cytokines Old/Microglia IHC +/- Rapa | 1B, 1D |  |
| 4 | MIR | Blood Cytokines Old/Microglia IHC +/- Rapa | 1B, 1D |  |
| 5 | CNTL | repetitive behaviors Old +/- Rapa/Longevity of Rapa/Chronic Rapa | 3C-E |  |
| 6 | CNTL | repetitive behaviors Old +/- Rapa/Longevity of Rapa/Chronic Rapa | 3C-E |  |
| 7 | CNTL | repetitive behaviors Old +/- Rapa/Longevity of Rapa/Chronic Rapa | 3C-E |  |
| 8 | MIR | repetitive behaviors Old +/- Rapa/Longevity of Rapa/Chronic Rapa | 3C-E |  |
| 9 | MIR | repetitive behaviors Old +/- Rapa/Longevity of Rapa/Chronic Rapa | 3C-E |  |
| 10 | MIR | repetitive behaviors Old +/- Rapa/Longevity of Rapa/Chronic Rapa | 3C-E |  |
| 11 | CNTL | repetitive behaviors Old/light-dark box/Von Frey/PPI/MRI +/- Rapa/Egfr1 IHC | 4A-F, 2B-C, 7A-F, S1 |  |
| 12 | CNTL | repetitive behaviors Old/light-dark box/Von Frey/PPI/MRI +/- Rapa/Egfr1 IHC | 4A-F, 2B-C, 7A-F, S1 |  |
| 13 | CNTL | repetitive behaviors Old/light-dark box/Von Frey/PPI/MRI +/- Rapa/Egfr1 IHC | 4A-F, 2B-C, 7A-F, S1 |  |
| 14 | CNTL | repetitive behaviors Old/light-dark box/Von Frey/PPI/MRI +/- Rapa/Egfr1 IHC | 4A-F, 2B-C, 7A-F, S1 |  |
| 15 | MIR | repetitive behaviors Old/light-dark box/Von Frey/PPI/MRI +/- Rapa/Egfr1 IHC | 4A-F, 2B-C, 7A-F, S1 |  |
| 16 | MIR | repetitive behaviors Old/light-dark box/Von Frey/PPI/MRI +/- Rapa/Egfr1 IHC | 4A-F, 2B-C, 7A-F, S1 |  |
| 17 | MIR | repetitive behaviors Old/light-dark box/Von Frey/PPI/MRI +/- Rapa/Egfr1 IHC | 4A-F, 2B-C, 7A-F, S1 |  |
| 18 | MIR | repetitive behaviors Old/light-dark box/Von Frey/PPI/MRI +/- Rapa/Egfr1 IHC | 4A-F, 2B-C, 7A-F, S1 |  |
| 19 | CNTL | repetitive behaviors Old/Von Frey +/- PLEX5622/e-Phys slice recording +/- Rapa | 1F-G, 6A-D, S2 | T1 |
| 20 | CNTL | repetitive behaviors Old/Von Frey +/- PLEX5622/e-Phys slice recording +/- Rapa | 1F-G, 6A-D, S2 | T1 |
| 21 | MIR | repetitive behaviors Old/Von Frey +/- PLEX5622/e-Phys slice recording +/- Rapa | 1F-G, 6A-D, S2 | T1 |
| 22 | MIR | repetitive behaviors Old/Von Frey +/- PLEX5622/e-Phys slice recording +/- Rapa | 1F-G, 6A-D, S2 | T1 |
| 23 | CNTL | repetitive behaviors Old/Von Frey +/- PLEX5622/e-Phys slice recording +/- Rapa | 1F-G, 6A-D, S2 | T1 |
| 24 | CNTL | repetitive behaviors Old/Von Frey +/- PLEX5622/e-Phys slice recording +/- Rapa | 1F-G, 6A-D, S2 | T1 |
| 25 | MIR | repetitive behaviors Old/Von Frey +/- PLEX5622/e-Phys slice recording +/- Rapa | 1F-G, 6A-D, S2 | T1 |
| 26 | MIR | repetitive behaviors Old/Von Frey +/- PLEX5622/e-Phys slice recording +/- Rapa | 1F-G, 6A-D, S2 | T1 |
| 27 | CNTL | repetitive behaviors Young/Von Frey/PPI/light-dark box +/- Rapa +/- PF470 | 3C, 4A-F, 5C, 5E |  |
| 28 | CNTL | repetitive behaviors Young/Von Frey/PPI/light-dark box +/- Rapa +/- PF470 | 3C, 4A-F, 5C, 5E |  |
| 29 | CNTL | repetitive behaviors Young/Von Frey/PPI/light-dark box +/- Rapa +/- PF470 | 3C, 4A-F, 5C, 5E |  |
| 30 | MIR | repetitive behaviors Young/Von Frey/PPI/light-dark box +/- Rapa +/- PF470 | 3C, 4A-F, 5C, 5E |  |
| 31 | MIR | repetitive behaviors Young/Von Frey/PPI/light-dark box +/- Rapa +/- PF470 | 3C, 4A-F, 5C, 5E |  |
| 32 | MIR | repetitive behaviors Young/Von Frey/PPI/light-dark box +/- Rapa +/- PF470 | 3C, 4A-F, 5C, 5E |  |
| 33 | CNTL | repetitive behaviors Young/Von Frey +/- PLEX5622/Sequencing +/- Rapa/PTZ seizure | 1F-G, 8A-E, S3 | T2-6, ST1 |
| 34 | CNTL | repetitive behaviors Young/Von Frey +/- PLEX5622/Sequencing +/- Rapa/PTZ seizure | 1F-G, 8A-E, S3 | T2-6, ST1 |
| 35 | CNTL | repetitive behaviors Young/Von Frey +/- PLEX5622/Sequencing +/- Rapa/PTZ seizure | 1F-G, 8A-E, S3 | T2-6, ST1 |
| 36 | MIR | repetitive behaviors Young/Von Frey +/- PLEX5622/Sequencing +/- Rapa/PTZ seizure | 1F-G, 8A-E, S3 | T2-6, ST1 |
| 37 | MIR | repetitive behaviors Young/Von Frey +/- PLEX5622/Sequencing +/- Rapa/PTZ seizure | 1F-G, 8A-E, S3 | T2-6, ST1 |
| 38 | MIR | repetitive behaviors Young/Von Frey +/- PLEX5622/Sequencing +/- Rapa/PTZ seizure | 1F-G, 8A-E, S3 | T2-6, ST1 |
| 39 | CNTL | repetitive behaviors +/- Rapa +/- Rapablock | 5A-B |  |
| 40 | CNTL | repetitive behaviors +/- Rapa +/- Rapablock | 5A-B |  |
| 41 | MIR | repetitive behaviors +/- Rapa +/- Rapablock | 5A-B |  |
| 42 | MIR | repetitive behaviors +/- Rapa +/- Rapablock | 5A-B |  |
| 43 | CNTL | Brain Wet Weight | 2A |  |
| 44 | CNTL | Brain Wet Weight | 2A |  |
| 45 | MIR | Brain Wet Weight | 2A |  |
| 46 | MIR | Brain Wet Weight | 2A |  |
| 47 | CNTL | PPI/light-dark box +/- Rapa +/- Rapablock +/- Apo/Reciprocal Social Interaction +/- Rapa +/- Rapablock | 1E, 3B, 5A-B, 5D, 5E, S1 |  |
| 48 | CNTL | PPI/light-dark box +/- Rapa +/- Rapablock +/- Apo/Reciprocal Social Interaction +/- Rapa +/- Rapablock | 1E, 3B, 5A-B, 5D, 5E, S1 |  |
| 49 | CNTL | PPI/light-dark box +/- Rapa +/- Rapablock +/- Apo/Reciprocal Social Interaction +/- Rapa +/- Rapablock | 1E, 3B, 5A-B, 5D, 5E, S1 |  |
| 50 | MIR | PPI/light-dark box +/- Rapa +/- Rapablock +/- Apo/Reciprocal Social Interaction +/- Rapa +/- Rapablock | 1E, 3B, 5A-B, 5D, 5E, S1 |  |
| 51 | MIR | PPI/light-dark box +/- Rapa +/- Rapablock +/- Apo/Reciprocal Social Interaction +/- Rapa +/- Rapablock | 1E, 3B, 5A-B, 5D, 5E, S1 |  |
| 52 | MIR | PPI/light-dark box +/- Rapa +/- Rapablock +/- Apo/Reciprocal Social Interaction +/- Rapa +/- Rapablock | 1E, 3B, 5A-B, 5D, 5E, S1 |  |
| Dams | Group | Dam Testing | Figure |  |
| 1-4 | CNTL | Blood Cytokines Pregnant Dams | 1C |  |
| 5-6 | MIR | Blood Cytokines Pregnant Dams | 1C |  |

**Supplemental Table 3.** Quantified Seizure Response by Mouse. The highest seizure level reached on the 1-6 scale (see methods for descriptors) was recorded for each mouse at: each PTZ dosage from 10-40 mg/kg), and at 40 mg/kg with 2 hour rapamycin treatment prior to PTZ administration. A 2-way ANOVA mixed model analysis for multiple comparisons showed significant treatment .x group (p<0.0001, DF=4), treatment (p<0.0001, DF=4), and group (p<0.0001, DF=1) effects. Posthoc Tukey’s multiple comparisons show that at 40mg/kg there is a significant difference between MIR and Control offspring (p=0.0001, DF=9.692) and between MIR 40mg/kg PTZ and MIR 40mg/kg PTZ + rapamycin (p=0.0303, DF=7). N=8/group/treatment

|  | PTZ dosage (mg/kg) |  |  |  |  |  |  |  |  |  |
| --- | --- | --- | --- | --- | --- | --- | --- | --- | --- | --- |
| seizure level mouse is in | 10 |  | 20 |  | 30 |  | 40 |  | 40 + Rapamycin |  |
| mouse | CNTL | MIR | CNTL | MIR | CNTL | MIR | CNTL | MIR | CNTL | MIR |
| m1 | 0 | 0 | 0 | 1 | 1 | 1 | 1 | 1 | 1 | 1 |
| m2 | 0 | 0 | 0 | 0 | 0 | 1 | 1 | 2 | 0 | 1 |
| m3 | 0 | 0 | 0 | 0 | 0 | 1 | 0 | 2 | 0 | 1 |
| m4 | 0 | 0 | 0 | 0 | 0 | 2 | 0 | 3 | 0 | 2 |
| m5 | 0 | 0 | 0 | 0 | 0 | 0 | 0 | 3 | 0 | 2 |
| m6 | 0 | 0 | 0 | 0 | 0 | 0 | 0 | 3 | 0 | 2 |
| m7 | 0 | 0 | 0 | 0 | 0 | 0 | 0 | 4 | 0 | 2 |
| m8 | 0 | 0 | 0 | 0 | 0 | 0 | 0 | 4 | 0 | 3 |

## Supplemental Methods

### Reciprocal social Interactions

Experimental and age- and sex-matched juvenile (P25) stranger mice were placed together in standard housing cages with clean bedding and both social and non-social behaviors were measured for the number of bouts of each during a 10-minute session as described by Silverman et al., 2010. Social parameters included following (experimental mouse walks slowly behind the stranger mouse, keeping pace), push-crawl (physical contact including pushing the snout or head underneath the partner’s body, squeezing between the partner and the arena wall or floor, and crawling over or under the partner’s body), and sniffing (nose-to-nose or nose-to-anogenital). Non-social parameters include solo arena exploration and self-grooming. The main outcome measures are the number of bouts of each type of behavior the mice performed during the testing session. Social and non-social behavior bouts are compared.

## Notes

### Competing Interest Statement

The authors have declared no competing interest.

### Summary of Updates

The reciprocal social interaction test has been added to supplemental data and the three chamber social approach test has been removed at the recommendation of peer review. Additional experimental details have been added to the manucscipt in accordance with the ARRIVE 2 guidelines. Some data has been re-graphed using violin plots to better display the individual data points from independent animals. Discussion of a new large-scale cross-species fMRI study of functional brain connectivity subtypes in autism from the Gozzi lab that aligns with our fMRI findgins has been added to the manuscript. doi: https://doi.org/10.1101/2025.03.04.641400

## References

1. Tanaka, M., Spekker, E., Szabó, Á., Polyák, H. & Vécsei, L. Modelling the neurodevelopmental pathogenesis in neuropsychiatric disorders. Bioactive kynurenines and their analogues as neuroprotective agents-in celebration of 80th birthday of Professor Peter Riederer. J Neural Transm (Vienna) 129, 627–642 (2022).

2. Li, Y.-J., Zhang, X. & Li, Y.-M. Antineuroinflammatory therapy: potential treatment for autism spectrum disorder by inhibiting glial activation and restoring synaptic function. CNS Spectr 25, 493–501 (2020).

3. Chen, J., Alberts, I. & Li, X. Dysregulation of the IGF-I/PI3K/AKT/mTOR signaling pathway in autism spectrum disorders. Int J Dev Neurosci 35, 35–41 (2014).

4. Thomas, S. D., Jha, N. K., Ojha, S. & Sadek, B. mTOR Signaling Disruption and Its Association with the Development of Autism Spectrum Disorder. Molecules 28, 1889 (2023).

5. Arenella, M. et al. Immunogenetics of autism spectrum disorder: A systematic literature review. Brain Behav Immun 114, 488–499 (2023).

6. Lugo, J. N. et al. Deletion of PTEN produces autism-like behavioral deficits and alterations in synaptic proteins. Front Mol Neurosci 7, 27 (2014).

7. Kwon, C.-H. et al. Pten Regulates Neuronal Arborization and Social Interaction in Mice. Neuron 50, 377–388 (2006).

8. Rademacher, S. & Eickholt, B. J. PTEN in Autism and Neurodevelopmental Disorders. Cold Spring Harb Perspect Med 9, a036780 (2019).

9. Zhou, J. et al. Pharmacological inhibition of mTORC1 suppresses anatomical, cellular, and behavioral abnormalities in neural-specific Pten knock-out mice. J Neurosci 29, 1773–1783 (2009).

10. Ljungberg, M. C., Sunnen, C. N., Lugo, J. N., Anderson, A. E. & D’Arcangelo, G. Rapamycin suppresses seizures and neuronal hypertrophy in a mouse model of cortical dysplasia. Dis Model Mech 2, 389–398 (2009).

11. Nguyen, L. H. et al. mTOR inhibition suppresses established epilepsy in a mouse model of cortical dysplasia. Epilepsia 56, 636–646 (2015).

12. Sunnen, C. N. et al. Inhibition of the mammalian target of rapamycin blocks epilepsy progression in NS-Pten conditional knockout mice. Epilepsia 52, 2065–2075 (2011).

13. Amaral, D. G., Schumann, C. M. & Nordahl, C. W. Neuroanatomy of autism. Trends Neurosci 31, 137–145 (2008).

14. Lee, J. K. et al. Longitudinal Evaluation of Cerebral Growth Across Childhood in Boys and Girls With Autism Spectrum Disorder. Biol Psychiatry 90, 286–294 (2021).

15. Libero, L. E. et al. Persistence of megalencephaly in a subgroup of young boys with autism spectrum disorder. Autism Res 9, 1169–1182 (2016).

16. van Rooij, D. et al. Cortical and Subcortical Brain Morphometry Differences Between Patients With Autism Spectrum Disorder and Healthy Individuals Across the Lifespan: Results From the ENIGMA ASD Working Group. Am J Psychiatry 175, 359–369 (2018).

17. Allen, M. et al. Astrocytes derived from ASD individuals alter behavior and destabilize neuronal activity through aberrant Ca2+ signaling. Mol Psychiatry 27, 2470–2484 (2022).

18. Vakilzadeh, G. & Martinez-Cerdeño, V. Pathology and Astrocytes in Autism. Neuropsychiatr Dis Treat 19, 841–850 (2023).

19. López-Aranda, M. F. et al. Postnatal immune activation causes social deficits in a mouse model of tuberous sclerosis: Role of microglia and clinical implications. Sci Adv 7, eabf2073 (2021).

20. Specchio, N. et al. Autism and Epilepsy in Patients With Tuberous Sclerosis Complex. Front Neurol 11, 639 (2020).

21. Cummings, K., Watkins, A., Jones, C., Dias, R. & Welham, A. Behavioural and psychological features of PTEN mutations: a systematic review of the literature and meta-analysis of the prevalence of autism spectrum disorder characteristics. J Neurodev Disord 14, 1 (2022).

22. Pagani, M. et al. mTOR-related synaptic pathology causes autism spectrum disorder-associated functional hyperconnectivity. Nat Commun 12, 6084 (2021).

23. Zahedi Abghari, F., Moradi, Y. & Akouchekian, M. PTEN gene mutations in patients with macrocephaly and classic autism: A systematic review. Med J Islam Repub Iran 33, 10 (2019).

24. Kim, W.-Y. Brain size is controlled by the mammalian target of rapamycin (mTOR) in mice. Commun Integr Biol 8, e994377 (2015).

25. LiCausi, F. & Hartman, N. W. Role of mTOR Complexes in Neurogenesis. Int J Mol Sci 19, 1544 (2018).

26. Winden, K. D., Ebrahimi-Fakhari, D. & Sahin, M. Abnormal mTOR Activation in Autism. Annu Rev Neurosci 41, 1–23 (2018).

27. Zoghbi, H. Y. & Bear, M. F. Synaptic dysfunction in neurodevelopmental disorders associated with autism and intellectual disabilities. Cold Spring Harb Perspect Biol 4, a009886 (2012).

28. Onore, C., Yang, H., Van de Water, J. & Ashwood, P. Dynamic Akt/mTOR Signaling in Children with Autism Spectrum Disorder. Front Pediatr 5, 43 (2017).

29. Costa-Mattioli, M. & Monteggia, L. M. mTOR complexes in neurodevelopmental and neuropsychiatric disorders. Nat Neurosci 16, 1537–1543 (2013).

30. Lipton, J. O. & Sahin, M. The neurology of mTOR. Neuron 84, 275–291 (2014).

31. Sato, A. & Ikeda, K. Genetic and Environmental Contributions to Autism Spectrum Disorder Through Mechanistic Target of Rapamycin. Biol Psychiatry Glob Open Sci 2, 95–105 (2022).

32. Groszer, M. et al. PTEN negatively regulates neural stem cell self-renewal by modulating G0-G1 cell cycle entry. Proc Natl Acad Sci U S A 103, 111–116 (2006).

33. Gregorian, C. et al. Pten Deletion in Adult Neural Stem/Progenitor Cells Enhances Constitutive Neurogenesis. J Neurosci 29, 1874–1886 (2009).

34. Hui, K. K. & Tanaka, M. Autophagy links MTOR and GABA signaling in the brain. Autophagy 15, 1848–1849 (2019).

35. Antoine, M. W., Langberg, T., Schnepel, P. & Feldman, D. E. Increased Excitation-Inhibition Ratio Stabilizes Synapse and Circuit Excitability in Four Autism Mouse Models. Neuron 101, 648–661.e4 (2019).

36. Chaudry, S. & Vasudevan, N. mTOR-Dependent Spine Dynamics in Autism. Front Mol Neurosci 15, 877609 (2022).

37. Kassai, H. et al. Selective activation of mTORC1 signaling recapitulates microcephaly, tuberous sclerosis, and neurodegenerative diseases. Cell Rep 7, 1626–1639 (2014).

38. Sadowski, K., Kotulska-Jóźwiak, K. & Jóźwiak, S. Role of mTOR inhibitors in epilepsy treatment. Pharmacological Reports 67, 636–646 (2015).

39. Marotta, R. et al. The Neurochemistry of Autism. Brain Sci 10, 163 (2020).

40. Luo, C. et al. Perfect match: mTOR inhibitors and tuberous sclerosis complex. Orphanet J Rare Dis 17, 106 (2022).

41. Laplante, M. & Sabatini, D. M. mTOR signaling in growth control and disease. Cell 149, 274–293 (2012).

42. Angliker, N., Burri, M., Zaichuk, M., Fritschy, J.-M. & Rüegg, M. A. mTORC1 and mTORC2 have largely distinct functions in Purkinje cells. Eur J Neurosci 42, 2595–2612 (2015).

43. McCabe, M. P. et al. Genetic inactivation of mTORC1 or mTORC2 in neurons reveals distinct functions in glutamatergic synaptic transmission. Elife 9, e51440 (2020).

44. Backman, S. A. et al. Deletion of Pten in mouse brain causes seizures, ataxia and defects in soma size resembling Lhermitte-Duclos disease. Nat Genet 29, 396–403 (2001).

45. Williams, K. A. & Swedo, S. E. Post-infectious autoimmune disorders: Sydenham’s chorea, PANDAS and beyond. Brain Res 1617, 144–154 (2015).

46. Weston, M. C., Chen, H. & Swann, J. W. Loss of mTOR repressors Tsc1 or Pten has divergent effects on excitatory and inhibitory synaptic transmission in single hippocampal neuron cultures. Front Mol Neurosci 7, 1 (2014).

47. Kwon, C. H. et al. Pten regulates neuronal soma size: a mouse model of Lhermitte-Duclos disease. Nat Genet 29, 404–411 (2001).

48. Barrows, C. M., McCabe, M. P., Chen, H., Swann, J. W. & Weston, M. C. PTEN Loss Increases the Connectivity of Fast Synaptic Motifs and Functional Connectivity in a Developing Hippocampal Network. J Neurosci 37, 8595–8611 (2017).

49. Trifonova, E. A., Mustafin, Z. S., Lashin, S. A. & Kochetov, A. V. Abnormal mTOR Activity in Pediatric Autoimmune Neuropsychiatric and MIA-Associated Autism Spectrum Disorders. Int J Mol Sci 23, 967 (2022).

50. Zhao, J.-P. & Yoshii, A. Hyperexcitability of the local cortical circuit in mouse models of tuberous sclerosis complex. Mol Brain 12, 6 (2019).

51. Malik, R. et al. Tsc1 represses parvalbumin expression and fast-spiking properties in somatostatin lineage cortical interneurons. Nat Commun 10, 4994 (2019).

52. Artinian, J. et al. Regulation of Hippocampal Memory by mTORC1 in Somatostatin Interneurons. J Neurosci 39, 8439–8456 (2019).

53. Fu, C. et al. GABAergic Interneuron Development and Function Is Modulated by the Tsc1 Gene. Cereb Cortex 22, 2111–2119 (2012).

54. Haji, N. et al. Tsc1 haploinsufficiency in Nkx2.1 cells upregulates hippocampal interneuron mTORC1 activity, impairs pyramidal cell synaptic inhibition, and alters contextual fear discrimination and spatial working memory in mice. Mol Autism 11, 29 (2020).

55. Weston, M. C., Chen, H. & Swann, J. W. Multiple Roles for Mammalian Target of Rapamycin Signaling in Both Glutamatergic and GABAergic Synaptic Transmission. J Neurosci 32, 11441–11452 (2012).

56. Marchese, M. et al. Autism-epilepsy phenotype with macrocephaly suggests PTEN, but not GLIALCAM, genetic screening. BMC Med Genet 15, 26 (2014).

57. Jansen, L. A. et al. PI3K/AKT pathway mutations cause a spectrum of brain malformations from megalencephaly to focal cortical dysplasia. Brain 138, 1613–1628 (2015).

58. Koboldt, D. C. et al. PTEN somatic mutations contribute to spectrum of cerebral overgrowth. Brain 144, 2971–2978 (2021).

59. Rosina, E. et al. Disruption of mTOR and MAPK pathways correlates with severity in idiopathic autism. Transl Psychiatry 9, 1–10 (2019).

60. Arenella, M., Mota, N. R., Teunissen, M. W. A., Brunner, H. G. & Bralten, J. Autism spectrum disorder and brain volume link through a set of mTOR-related genes. J Child Psychol Psychiatry 64, 1007–1014 (2023).

61. Aldinger, K. A., Plummer, J. T., Qiu, S. & Levitt, P. SnapShot: genetics of autism. Neuron 72, 418–418.e1 (2011).

62. Xing, X. et al. Hyperactive Akt-mTOR pathway as a therapeutic target for pain hypersensitivity in Cntnap2-deficient mice. Neuropharmacology 165, 107816 (2020).

63. Trifonova, E. A., Klimenko, A. I., Mustafin, Z. S., Lashin, S. A. & Kochetov, A. V. The mTOR Signaling Pathway Activity and Vitamin D Availability Control the Expression of Most Autism Predisposition Genes. Int J Mol Sci 20, 6332 (2019).

64. Bi, X., Sun, J., Ji, A. X. & Baudry, M. Potential therapeutic approaches for Angelman syndrome. Expert Opin Ther Targets 20, 601–613 (2016).

65. Kabitzke, P. A. et al. Comprehensive analysis of two Shank3 and the Cacna1c mouse models of autism spectrum disorder. Genes Brain Behav 17, 4–22 (2018).

66. Roy, B., Amemasor, E., Hussain, S. & Castro, K. UBE3A: The Role in Autism Spectrum Disorders (ASDs) and a Potential Candidate for Biomarker Studies and Designing Therapeutic Strategies. Diseases 12, 7 (2023).

67. Canales, C. P. et al. Sequential perturbations to mouse corticogenesis following in utero maternal immune activation. Elife 10, e60100 (2021).

68. Basu, S. N., Kollu, R. & Banerjee-Basu, S. AutDB: a gene reference resource for autism research. Nucleic Acids Res 37, D832–836 (2009).

69. Leblond, C. S. et al. Operative list of genes associated with autism and neurodevelopmental disorders based on database review. Mol Cell Neurosci 113, 103623 (2021).

70. Wen, Y., Alshikho, M. J. & Herbert, M. R. Pathway Network Analyses for Autism Reveal Multisystem Involvement, Major Overlaps with Other Diseases and Convergence upon MAPK and Calcium Signaling. PLoS One 11, e0153329 (2016).

71. Hu, Y. et al. mTOR-mediated metabolic reprogramming shapes distinct microglia functions in response to lipopolysaccharide and ATP. Glia 68, 1031–1045 (2020).

72. Hodges, S. L. & Lugo, J. N. Therapeutic role of targeting mTOR signaling and neuroinflammation in epilepsy. Epilepsy Res 161, 106282 (2020).

73. Haidinger, M. et al. A versatile role of mammalian target of rapamycin in human dendritic cell function and differentiation. J Immunol 185, 3919–3931 (2010).

74. Lehman, J. A., Calvo, V. & Gomez-Cambronero, J. Mechanism of ribosomal p70S6 kinase activation by granulocyte macrophage colony-stimulating factor in neutrophils: cooperation of a MEK-related, THR421/SER424 kinase and a rapamycin-sensitive, m-TOR-related THR389 kinase. J Biol Chem 278, 28130–28138 (2003).

75. Weichhart, T., Hengstschläger, M. & Linke, M. Regulation of innate immune cell function by mTOR. Nat Rev Immunol 15, 599–614 (2015).

76. Xiao, Z., Peng, J., Wu, L., Arafat, A. & Yin, F. The effect of IL-1β on synaptophysin expression and electrophysiology of hippocampal neurons through the PI3K/Akt/mTOR signaling pathway in a rat model of mesial temporal lobe epilepsy. Neurol Res 39, 640–648 (2017).

77. Argandona Lopez, C. & Brown, A. M. Microglial-neuronal crosstalk in chronic viral infection through mTOR, SPP1/OPN and inflammasome pathway signaling. Front Immunol 15, 1368465 (2024).

78. Lashgari, N.-A. et al. TLR/mTOR inflammatory signaling pathway: novel insight for the treatment of schizophrenia. Can J Physiol Pharmacol 102, 150–160 (2024).

79. Sharma, A. & Mehan, S. Targeting PI3K-AKT/mTOR signaling in the prevention of autism. Neurochem Int 147, 105067 (2021).

80. Yang, H. et al. Modulation of TSC-mTOR signaling on immune cells in immunity and autoimmunity. J Cell Physiol 229, 17–26 (2014).

81. Byles, V. et al. The TSC-mTOR pathway regulates macrophage polarization. Nat Commun 4, 2834 (2013).

82. Vergadi, E., Ieronymaki, E., Lyroni, K., Vaporidi, K. & Tsatsanis, C. Akt Signaling Pathway in Macrophage Activation and M1/M2 Polarization. J Immunol 198, 1006–1014 (2017).

83. Krakowiak, P. et al. Neonatal Cytokine Profiles Associated With Autism Spectrum Disorder. Biol Psychiatry 81, 442–451 (2017).

84. Estes, M. L. & McAllister, A. K. Immune mediators in the brain and peripheral tissues in autism spectrum disorder. Nat Rev Neurosci 16, 469–486 (2015).

85. Patterson, P. H. Immune involvement in schizophrenia and autism: etiology, pathology and animal models. Behav Brain Res 204, 313–321 (2009).

86. Xiong, Y., Chen, J. & Li, Y. Microglia and astrocytes underlie neuroinflammation and synaptic susceptibility in autism spectrum disorder. Front Neurosci 17, 1125428 (2023).

87. Matta, S. M., Hill-Yardin, E. L. & Crack, P. J. The influence of neuroinflammation in Autism Spectrum Disorder. Brain Behav Immun 79, 75–90 (2019).

88. Vargas, D. L., Nascimbene, C., Krishnan, C., Zimmerman, A. W. & Pardo, C. A. Neuroglial activation and neuroinflammation in the brain of patients with autism. Ann Neurol 57, 67–81 (2005).

89. Gesundheit, B. et al. Immunological and autoimmune considerations of Autism Spectrum Disorders. J Autoimmun 44, 1–7 (2013).

90. Theoharides, T. C., Asadi, S. & Patel, A. B. Focal brain inflammation and autism. J Neuroinflammation 10, 46 (2013).

91. Voineagu, I. et al. Transcriptomic analysis of autistic brain reveals convergent molecular pathology. Nature 474, 380–384 (2011).

92. Morgan, J. T. et al. Microglial activation and increased microglial density observed in the dorsolateral prefrontal cortex in autism. Biol Psychiatry 68, 368–376 (2010).

93. Laurence, J. A. & Fatemi, S. H. Glial fibrillary acidic protein is elevated in superior frontal, parietal and cerebellar cortices of autistic subjects. Cerebellum 4, 206–210 (2005).

94. Edmonson, C., Ziats, M. N. & Rennert, O. M. Altered glial marker expression in autistic post-mortem prefrontal cortex and cerebellum. Mol Autism 5, 3 (2014).

95. Masi, A., Glozier, N., Dale, R. & Guastella, A. J. The Immune System, Cytokines, and Biomarkers in Autism Spectrum Disorder. Neurosci Bull 33, 194–204 (2017).

96. Gandal, M. J. et al. Transcriptome-wide isoform-level dysregulation in ASD, schizophrenia, and bipolar disorder. Science 362, eaat8127 (2018).

97. Li, X. et al. Elevated immune response in the brain of autistic patients. J Neuroimmunol 207, 111–116 (2009).

98. Courchesne, E. Brain development in autism: early overgrowth followed by premature arrest of growth. Ment Retard Dev Disabil Res Rev 10, 106–111 (2004).

99. Courchesne, E., Campbell, K. & Solso, S. Brain growth across the life span in autism: age-specific changes in anatomical pathology. Brain Res 1380, 138–145 (2011).

100. Redcay, E. & Courchesne, E. When is the brain enlarged in autism? A meta-analysis of all brain size reports. Biol Psychiatry 58, 1–9 (2005).

101. Fatemi, S. H. et al. Prenatal viral infection leads to pyramidal cell atrophy and macrocephaly in adulthood: implications for genesis of autism and schizophrenia. Cell Mol Neurobiol 22, 25–33 (2002).

102. Smith, S. E. P., Li, J., Garbett, K., Mirnics, K. & Patterson, P. H. Maternal Immune Activation Alters Fetal Brain Development through Interleukin-6. J Neurosci 27, 10695–10702 (2007).

103. Shi, L., Fatemi, S. H., Sidwell, R. W. & Patterson, P. H. Maternal influenza infection causes marked behavioral and pharmacological changes in the offspring. J Neurosci 23, 297–302 (2003).

104. Malkova, N. V., Yu, C. Z., Hsiao, E. Y., Moore, M. J. & Patterson, P. H. Maternal immune activation yields offspring displaying mouse versions of the three core symptoms of autism. Brain Behav Immun 26, 607–616 (2012).

105. Choi, G. B. et al. The maternal interleukin-17a pathway in mice promotes autism-like phenotypes in offspring. Science 351, 933–939 (2016).

106. Atladóttir, H. O. et al. Association of family history of autoimmune diseases and autism spectrum disorders. Pediatrics 124, 687–694 (2009).

107. Atladóttir, H. O. et al. Maternal infection requiring hospitalization during pregnancy and autism spectrum disorders. J Autism Dev Disord 40, 1423–1430 (2010).

108. Brown, A. S. et al. Elevated maternal C-reactive protein and autism in a national birth cohort. Mol Psychiatry 19, 259–264 (2014).

109. Ashwood, P., Wills, S. & Van de Water, J. The immune response in autism: a new frontier for autism research. J Leukoc Biol 80, 1–15 (2006).

110. Lee, B. K. et al. Maternal hospitalization with infection during pregnancy and risk of autism spectrum disorders. Brain Behav Immun 44, 100–105 (2015).

111. Modabbernia, A., Velthorst, E. & Reichenberg, A. Environmental risk factors for autism: an evidence-based review of systematic reviews and meta-analyses. Mol Autism 8, 13 (2017).

112. Goines, P. E. & Ashwood, P. Cytokine dysregulation in autism spectrum disorders (ASD): possible role of the environment. Neurotoxicol Teratol 36, 67–81 (2013).

113. Jiang, H.-Y. et al. Maternal infection during pregnancy and risk of autism spectrum disorders: A systematic review and meta-analysis. Brain Behav Immun 58, 165–172 (2016).

114. Kolevzon, A., Gross, R. & Reichenberg, A. Prenatal and perinatal risk factors for autism: a review and integration of findings. Arch Pediatr Adolesc Med 161, 326–333 (2007).

115. Kancherla, V. & Dennis, L. K. A Meta-analysis of Prenatal, Perinatal, and Neonatal Risks for Autism. American Journal of Epidemiology 163, S20 (2006).

116. Gardener, H., Spiegelman, D. & Buka, S. L. Prenatal Risk Factors for Autism: A Comprehensive Meta-analysis. Br J Psychiatry 195, 7–14 (2009).

117. Gardener, H., Spiegelman, D. & Buka, S. L. Perinatal and neonatal risk factors for autism: a comprehensive meta-analysis. Pediatrics 128, 344–355 (2011).

118. Le Belle, J. E. et al. Maternal inflammation contributes to brain overgrowth and autism-associated behaviors through altered redox signaling in stem and progenitor cells. Stem Cell Reports 3, 725–734 (2014).

119. Hazen, E. P., Stornelli, J. L., O’Rourke, J. A., Koesterer, K. & McDougle, C. J. Sensory symptoms in autism spectrum disorders. Harv Rev Psychiatry 22, 112–124 (2014).

120. Narvaiz, D. A. et al. Rapamycin improves social and stereotypic behavior abnormalities induced by pre-mitotic neuronal subset specific Pten deletion. Genes Brain Behav 22, e12854 (2023).

121. Zhang, J., Zhang, J.-X. & Zhang, Q.-L. PI3K/AKT/mTOR-mediated autophagy in the development of autism spectrum disorder. Brain Res Bull 125, 152–158 (2016).

122. Lieberman, O. J. et al. mTOR Suppresses Macroautophagy During Striatal Postnatal Development and Is Hyperactive in Mouse Models of Autism Spectrum Disorders. Front Cell Neurosci 14, 70 (2020).

123. Ehninger, D. & Silva, A. J. Rapamycin for treating Tuberous sclerosis and Autism spectrum disorders. Trends Mol Med 17, 78–87 (2011).

124. Ehninger, D. et al. Reversal of learning deficits in a Tsc2+/- mouse model of tuberous sclerosis. Nat Med 14, 843–848 (2008).

125. Kotajima-Murakami, H. et al. Effects of rapamycin on social interaction deficits and gene expression in mice exposed to valproic acid in utero. Mol Brain 12, 3 (2019).

126. Meikle, L. et al. Response of a Neuronal Model of Tuberous Sclerosis to Mammalian Target of Rapamycin (mTOR) Inhibitors: Effects on mTORC1 and Akt Signaling Lead to Improved Survival and Function. J Neurosci 28, 5422–5432 (2008).

127. Getz, S. A., DeSpenza, T., Li, M. & Luikart, B. W. Rapamycin prevents, but does not reverse, aberrant migration in Pten knockout neurons. Neurobiol Dis 93, 12–20 (2016).

128. Amegandjin, C. A. et al. Sensitive period for rescuing parvalbumin interneurons connectivity and social behavior deficits caused by TSC1 loss. Nat Commun 12, 3653 (2021).

129. Tsai, P. T. et al. Autistic-like behaviour and cerebellar dysfunction in Purkinje cell Tsc1 mutant mice. Nature 488, 647–651 (2012).

130. Zeng, L.-H., Xu, L., Gutmann, D. H. & Wong, M. Rapamycin prevents epilepsy in a mouse model of tuberous sclerosis complex. Ann Neurol 63, 444–453 (2008).

131. Tsai, P. T. et al. Sensitive Periods for Cerebellar-Mediated Autistic-like Behaviors. Cell Rep 25, 357–367.e4 (2018).

132. Tang, G. et al. Loss of mTOR-dependent macroautophagy causes autistic-like synaptic pruning deficits. Neuron 83, 1131–1143 (2014).

133. Krueger, D. A. et al. Everolimus for subependymal giant-cell astrocytomas in tuberous sclerosis. N Engl J Med 363, 1801–1811 (2010).

134. Krueger, D. A. et al. Everolimus treatment of refractory epilepsy in tuberous sclerosis complex. Ann Neurol 74, 679–687 (2013).

135. Cardamone, M. et al. Mammalian target of rapamycin inhibitors for intractable epilepsy and subependymal giant cell astrocytomas in tuberous sclerosis complex. J Pediatr 164, 1195–1200 (2014).

136. Kotulska, K. et al. Long-term effect of everolimus on epilepsy and growth in children under 3 years of age treated for subependymal giant cell astrocytoma associated with tuberous sclerosis complex. Eur J Paediatr Neurol 17, 479–485 (2013).

137. MacKeigan, J. P. & Krueger, D. A. Differentiating the mTOR inhibitors everolimus and sirolimus in the treatment of tuberous sclerosis complex. Neuro Oncol 17, 1550–1559 (2015).

138. Srivastava, S. et al. A randomized controlled trial of everolimus for neurocognitive symptoms in PTEN hamartoma tumor syndrome. Hum Mol Genet 31, 3393–3404 (2022).

139. Overwater, I. E. et al. A randomized controlled trial with everolimus for IQ and autism in tuberous sclerosis complex. Neurology 93, e200–e209 (2019).

140. Davies, D. M. et al. Sirolimus therapy for angiomyolipoma in tuberous sclerosis and sporadic lymphangioleiomyomatosis: a phase 2 trial. Clin Cancer Res 17, 4071–4081 (2011).

141. Mizuguchi, M. et al. Everolimus for epilepsy and autism spectrum disorder in tuberous sclerosis complex: EXIST-3 substudy in Japan. Brain Dev 41, 1–10 (2019).

142. Hwang, S.-K. et al. Everolimus improves neuropsychiatric symptoms in a patient with tuberous sclerosis carrying a novel TSC2 mutation. Mol Brain 9, 56 (2016).

143. Kilincaslan, A. et al. Beneficial Effects of Everolimus on Autism and Attention-Deficit/Hyperactivity Disorder Symptoms in a Group of Patients with Tuberous Sclerosis Complex. J Child Adolesc Psychopharmacol 27, 383–388 (2017).

144. Ess, K. C. & Franz, D. N. Everolimus for cognition/autism in children with tuberous sclerosis complex: Definitive outcomes deferred. Neurology 93, 51–52 (2019).

145. Saffari, A. et al. Safety and efficacy of mTOR inhibitor treatment in patients with tuberous sclerosis complex under 2 years of age - a multicenter retrospective study. Orphanet J Rare Dis 14, 96 (2019).

146. Hodges, S. L. et al. Rapamycin, but not minocycline, significantly alters ultrasonic vocalization behavior in C57BL/6J pups in a flurothyl seizure model. Behav Brain Res 410, 113317 (2021).

147. Qin, L., Dai, X. & Yin, Y. Valproic acid exposure sequentially activates Wnt and mTOR pathways in rats. Mol Cell Neurosci 75, 27–35 (2016).

148. McMahon, J. J. et al. Seizure-dependent mTOR activation in 5-HT neurons promotes autism-like behaviors in mice. Neurobiol Dis 73, 296–306 (2015).

149. Burket, J. A., Benson, A. D., Tang, A. H. & Deutsch, S. I. Rapamycin improves sociability in the BTBR T+Itpr3tf/J mouse model of autism spectrum disorders. Brain Res Bull 100, 70–75 (2014).

150. Percie du Sert, N., et al. The ARRIVE guidelines 2.0: Updated guidelines for reporting animal research. Br J Pharmacol 177, 3617–3624 (2020).

151. López-Aranda, M. F. et al. Postnatal immune activation causes social deficits in a mouse model of tuberous sclerosis: Role of microglia and clinical implications. Sci Adv 7, eabf2073 (2021).

152. Zhang, Z. et al. Brain-restricted mTOR inhibition with binary pharmacology. Nature 609, 822–828 (2022).

153. Ehinger, Y. et al. Brain-specific inhibition of mTORC1 eliminates side effects resulting from mTORC1 blockade in the periphery and reduces alcohol intake in mice. Nat Commun 12, 4407 (2021).

154. Huang, W.-C., Chen, Y. & Page, D. T. Hyperconnectivity of prefrontal cortex to amygdala projections in a mouse model of macrocephaly/autism syndrome. Nat Commun 7, 13421 (2016).

155. Shimada, T. & Yamagata, K. Pentylenetetrazole-Induced Kindling Mouse Model. J Vis Exp 56573 (2018) doi:10.3791/56573.

156. Peñagarikano, O. et al. Absence of CNTNAP2 leads to epilepsy, neuronal migration abnormalities, and core autism-related deficits. Cell 147, 235–246 (2011).

157. Bonin, R. P., Bories, C. & De Koninck, Y. A simplified up-down method (SUDO) for measuring mechanical nociception in rodents using von Frey filaments. Mol Pain 10, 26 (2014).

158. Crawley, J. & Goodwin, F. K. Preliminary report of a simple animal behavior model for the anxiolytic effects of benzodiazepines. Pharmacol Biochem Behav 13, 167–170 (1980).

159. Orefice, L. L. et al. Peripheral Mechanosensory Neuron Dysfunction Underlies Tactile and Behavioral Deficits in Mouse Models of ASDs. Cell 166, 299–313 (2016).

160. Fleming, S. J. et al. Unsupervised removal of systematic background noise from droplet-based single-cell experiments using CellBender. Nat Methods 20, 1323–1335 (2023).

161. Germain, P.-L., Lun, A., Garcia Meixide, C., Macnair, W. & Robinson, M. D. Doublet identification in single-cell sequencing data using scDblFinder. F1000Res 10, 979 (2021).

162. Heinemann, J. Cluster Analysis of Untargeted Metabolomic Experiments. Methods Mol Biol 1859, 275–285 (2019).

163. Stuart, T. et al. Comprehensive integration of single-cell data. Cell 177, 1888–1902.e21 (2019).

164. Hao, Y. et al. Integrated analysis of multimodal single-cell data. Cell 184, 3573–3587.e29 (2021).

165. Phipson, B. et al. propeller: testing for differences in cell type proportions in single cell data. Bioinformatics 38, 4720–4726 (2022).

166. Squair, J. W. et al. Confronting false discoveries in single-cell differential expression. Nat Commun 12, 5692 (2021).

167. Subramanian, A. et al. Gene set enrichment analysis: a knowledge-based approach for interpreting genome-wide expression profiles. Proc Natl Acad Sci U S A 102, 15545–15550 (2005).

168. Liberzon, A. et al. The Molecular Signatures Database (MSigDB) hallmark gene set collection. Cell Syst 1, 417–425 (2015).

169. Castanza, A. S. et al. Extending support for mouse data in the Molecular Signatures Database (MSigDB). Nat Methods 20, 1619–1620 (2023).

170. Korotkevich, G. et al. Fast gene set enrichment analysis. 060012 Preprint at 10.1101/060012 (2021).

171. Adamczak, J. M., Farr, T. D., Seehafer, J. U., Kalthoff, D. & Hoehn, M. High field BOLD response to forepaw stimulation in the mouse. Neuroimage 51, 704–712 (2010).

172. Magnuson, M. E., Thompson, G. J., Pan, W.-J. & Keilholz, S. D. Time-dependent effects of isoflurane and dexmedetomidine on functional connectivity, spectral characteristics, and spatial distribution of spontaneous BOLD fluctuations. NMR Biomed 27, 291–303 (2014).

173. Oguz, I., Zhang, H., Rumple, A. & Sonka, M. RATS: Rapid Automatic Tissue Segmentation in rodent brain MRI. J Neurosci Methods 221, 175–182 (2014).

174. Jenkinson, M. & Smith, S. A global optimisation method for robust affine registration of brain images. Med Image Anal 5, 143–156 (2001).

175. Avants, B. B. et al. The Insight ToolKit image registration framework. Front Neuroinform 8, 44 (2014).

176. Smith, S. M. et al. Advances in functional and structural MR image analysis and implementation as FSL. Neuroimage 23 **Suppl 1**, S208–219 (2004).

177. Zalesky, A., Fornito, A. & Bullmore, E. T. Network-based statistic: identifying differences in brain networks. Neuroimage 53, 1197–1207 (2010).

178. Kruschwitz, J. D., List, D., Waller, L., Rubinov, M. & Walter, H. GraphVar: a user-friendly toolbox for comprehensive graph analyses of functional brain connectivity. J Neurosci Methods 245, 107–115 (2015).

179. Worsley, K. J. Statistical analysis of activation images. in Functional Magnetic Resonance Imaging: An Introduction to Methods (eds Jezzard, P., Matthews, P. M. & Smith, S. M.) 0 (Oxford University Press, 2001). doi:10.1093/acprof:oso/9780192630711.003.0014.

180. Blondel, V. D., Guillaume, J.-L., Lambiotte, R. & Lefebvre, E. Fast unfolding of communities in large networks. J. Stat. Mech. 2008, P10008 (2008).

181. Fornito, A., Zalesky, A. & Breakspear, M. Graph analysis of the human connectome: promise, progress, and pitfalls. Neuroimage 80, 426–444 (2013).

182. M, J. BET2 : MR-Based Estimation of Brain, Skull and Scalp Surfaces. Eleventh Annual Meeting of the Organization for Human Brain Mapping, 2005 17, 167 (2005).

183. Avants, B. B. et al. A reproducible evaluation of ANTs similarity metric performance in brain image registration. Neuroimage 54, 2033–2044 (2011).

184. Avants, B. B., Epstein, C. L., Grossman, M. & Gee, J. C. Symmetric diffeomorphic image registration with cross-correlation: evaluating automated labeling of elderly and neurodegenerative brain. Med Image Anal 12, 26–41 (2008).

185. Winkler, A. M., Ridgway, G. R., Webster, M. A., Smith, S. M. & Nichols, T. E. Permutation inference for the general linear model. Neuroimage 92, 381–397 (2014).

186. Benjamini, Y. & Hochberg, Y. Controlling the False Discovery Rate: A Practical and Powerful Approach to Multiple Testing. Journal of the Royal Statistical Society: Series B (Methodological) 57, 289–300 (1995).

187. Genovese, C. R., Lazar, N. A. & Nichols, T. Thresholding of statistical maps in functional neuroimaging using the false discovery rate. Neuroimage 15, 870–878 (2002).

188. Braden, B. B. & Riecken, C. Thinning Faster? Age-Related Cortical Thickness Differences in Adults with Autism Spectrum Disorder. Res Autism Spectr Disord 64, 31–38 (2019).

189. Tamura, R., Kitamura, H., Endo, T., Hasegawa, N. & Someya, T. Reduced thalamic volume observed across different subgroups of autism spectrum disorders. Psychiatry Res 184, 186–188 (2010).

190. Qiu, T. et al. Two years changes in the development of caudate nucleus are involved in restricted repetitive behaviors in 2-5-year-old children with autism spectrum disorder. Dev Cogn Neurosci 19, 137–143 (2016).

191. Banker, S. M., Gu, X., Schiller, D. & Foss-Feig, J. H. Hippocampal contributions to social and cognitive deficits in autism spectrum disorder. Trends Neurosci 44, 793–807 (2021).

192. Schumann, C. M. et al. The amygdala is enlarged in children but not adolescents with autism; the hippocampus is enlarged at all ages. J Neurosci 24, 6392–6401 (2004).

193. Schumann, C. M. & Amaral, D. G. Stereological analysis of amygdala neuron number in autism. J Neurosci 26, 7674–7679 (2006).

194. Turner, A. H., Greenspan, K. S. & van Erp, T. G. M. Pallidum and lateral ventricle volume enlargement in autism spectrum disorder. Psychiatry Res Neuroimaging 252, 40–45 (2016).

195. Green, S. A. & Ben-Sasson, A. Anxiety Disorders and Sensory Over-Responsivity in Children with Autism Spectrum Disorders: Is There a Causal Relationship? J Autism Dev Disord 40, 1495–1504 (2010).

196. Simmons, D. H. et al. Sensory Over-responsivity and Aberrant Plasticity in Cerebellar Cortex in a Mouse Model of Syndromic Autism. Biol Psychiatry Glob Open Sci 2, 450–459 (2022).

197. Tavassoli, T., Miller, L. J., Schoen, S. A., Nielsen, D. M. & Baron-Cohen, S. Sensory over-responsivity in adults with autism spectrum conditions. Autism 18, 428–432 (2014).

198. Tavassoli, T. et al. Sensory over-responsivity: parent report, direct assessment measures, and neural architecture. Molecular Autism 10, 4 (2019).

199. Orefice, L. L. et al. Targeting peripheral somatosensory neurons to improve tactile-related phenotypes in ASD models. Cell 178, 867–886.e24 (2019).

200. Le Belle, J. E. et al. Proliferative neural stem cells have high endogenous ROS levels that regulate self-renewal and neurogenesis in a PI3K/Akt-dependant manner. Cell Stem Cell 8, 59–71 (2011).

201. Otsuka, T. & Kawaguchi, Y. Firing-pattern-dependent specificity of cortical excitatory feed-forward subnetworks. J Neurosci 28, 11186–11195 (2008).

202. Gandhi, T. & Lee, C. C. Neural Mechanisms Underlying Repetitive Behaviors in Rodent Models of Autism Spectrum Disorders. Front Cell Neurosci 14, 592710 (2021).

203. Hisaoka, T., Komori, T., Kitamura, T. & Morikawa, Y. Abnormal behaviours relevant to neurodevelopmental disorders in Kirrel3-knockout mice. Sci Rep 8, 1408 (2018).

204. Mejias, R. et al. Gain-of-function glutamate receptor interacting protein 1 variants alter GluA2 recycling and surface distribution in patients with autism. Proc Natl Acad Sci U S A 108, 4920–4925 (2011).

205. Mejias, R. et al. Purkinje cell-specific Grip1/2 knockout mice show increased repetitive self-grooming and enhanced mGluR5 signaling in cerebellum. Neurobiol Dis 132, 104602 (2019).

206. Gandal, M. J. et al. Shared molecular neuropathology across major psychiatric disorders parallels polygenic overlap. Science 359, 693–697 (2018).

207. Lewis, M. L., Kesler, M., Candy, S. A., Rho, J. M. & Pittman, Q. J. Comorbid epilepsy in autism spectrum disorder: Implications of postnatal inflammation for brain excitability. Epilepsia 59, 1316–1326 (2018).

208. Masi, A. et al. Cytokine aberrations in autism spectrum disorder: a systematic review and meta-analysis. Mol Psychiatry 20, 440–446 (2015).

209. Williams, J. A. et al. Inflammation and Brain Structure in Schizophrenia and Other Neuropsychiatric Disorders: A Mendelian Randomization Study. JAMA Psychiatry 79, 498–507 (2022).

210. Than, U. T. T. et al. Inflammatory mediators drive neuroinflammation in autism spectrum disorder and cerebral palsy. Sci Rep 13, 22587 (2023).

211. Kern, J. K., Geier, D. A., Sykes, L. K. & Geier, M. R. Relevance of Neuroinflammation and Encephalitis in Autism. Front Cell Neurosci 9, 519 (2015).

212. Onore, C., Careaga, M. & Ashwood, P. The role of immune dysfunction in the pathophysiology of autism. Brain Behav Immun 26, 383–392 (2012).

213. Pardo, C. A., Vargas, D. L. & Zimmerman, A. W. Immunity, neuroglia and neuroinflammation in autism. Int Rev Psychiatry 17, 485–495 (2005).

214. Ben-Sasson, A., Cermak, S. A., Orsmond, G. I., Carter, A. S. & Fogg, L. Can we differentiate sensory over-responsivity from anxiety symptoms in toddlers? perspectives of occupational therapists and psychologists. Infant Ment Health J 28, 536–558 (2007).

215. Ben-Sasson, A. et al. Sensory clusters of toddlers with autism spectrum disorders: differences in affective symptoms. J Child Psychol Psychiatry 49, 817–825 (2008).

216. Glod, M., Riby, D. M., Honey, E. & Rodgers, J. Psychological Correlates of Sensory Processing Patterns in Individuals with Autism Spectrum Disorder: A Systematic Review. Rev J Autism Dev Disord 2, 199–221 (2015).

217. Ausderau, K. K. et al. Sensory subtypes and associated outcomes in children with autism spectrum disorders. Autism Res 9, 1316–1327 (2016).

218. Siniscalco, D., Schultz, S., Brigida, A. L. & Antonucci, N. Inflammation and Neuro-Immune Dysregulations in Autism Spectrum Disorders. Pharmaceuticals (Basel) 11, 56 (2018).

219. Tartaglione, A. M. et al. Maternal immune activation induces autism-like changes in behavior, neuroinflammatory profile and gut microbiota in mouse offspring of both sexes. Transl Psychiatry 12, 384 (2022).

220. Kim, E. et al. Maternal gut bacteria drive intestinal inflammation in offspring with neurodevelopmental disorders by altering the chromatin landscape of CD4+ T cells. Immunity 55, 145–158.e7 (2022).

221. Shimizu, Y., Sakata-Haga, H., Saikawa, Y. & Hatta, T. Influence of Immune System Abnormalities Caused by Maternal Immune Activation in the Postnatal Period. Cells 12, 741 (2023).

222. Lucchina, L. & Depino, A. M. Altered peripheral and central inflammatory responses in a mouse model of autism. Autism Res 7, 273–289 (2014).

223. Hsiao, E. Y., McBride, S. W., Chow, J., Mazmanian, S. K. & Patterson, P. H. Modeling an autism risk factor in mice leads to permanent immune dysregulation. Proc Natl Acad Sci U S A 109, 12776–12781 (2012).

224. Kirsten, T. B., Casarin, R. C., Bernardi, M. M. & Felicio, L. F. Pioglitazone abolishes autistic-like behaviors via the IL-6 pathway. PLoS One 13, e0197060 (2018).

225. Kirsten, T. B., Casarin, R. C., Bernardi, M. M. & Felicio, L. F. Pioglitazone abolishes cognition impairments as well as BDNF and neurotensin disturbances in a rat model of autism. Biol Open 8, bio041327 (2019).

226. Sandhu, A. et al. Ameliorating effect of pioglitazone on prenatal valproic acid-induced behavioral and neurobiological abnormalities in autism spectrum disorder in rats. Pharmacol Biochem Behav 237, 173721 (2024).

227. Buehler, M. R. A proposed mechanism for autism: an aberrant neuroimmune response manifested as a psychiatric disorder. Med Hypotheses 76, 863–870 (2011).

228. Emanuele, E., Lossano, C., Politi, P. & Barale, F. Pioglitazone as a therapeutic agent in autistic spectrum disorder. Medical Hypotheses 69, 699 (2007).

229. Boris, M. et al. Effect of pioglitazone treatment on behavioral symptoms in autistic children. J Neuroinflammation 4, 3 (2007).

230. Mirza, R. & Sharma, B. Beneficial effects of pioglitazone, a selective peroxisome proliferator-activated receptor-γ agonist in prenatal valproic acid-induced behavioral and biochemical autistic like features in Wistar rats. Int J Dev Neurosci 76, 6–16 (2019).

231. Mirza, R. & Sharma, B. A selective peroxisome proliferator-activated receptor-γ agonist benefited propionic acid induced autism-like behavioral phenotypes in rats by attenuation of neuroinflammation and oxidative stress. Chem Biol Interact 311, 108758 (2019).

232. Capano, L. et al. A pilot dose finding study of pioglitazone in autistic children. Mol Autism 9, 59 (2018).

233. Reed, M. D. et al. IL-17a promotes sociability in mouse models of neurodevelopmental disorders. Nature 577, 249–253 (2020).

234. Luo, Y. et al. Minocycline improves autism-related behaviors by modulating microglia polarization in a mouse model of autism. Int Immunopharmacol 122, 110594 (2023).

235. Kim, H.-J. et al. Deficient autophagy in microglia impairs synaptic pruning and causes social behavioral defects. Mol Psychiatry 22, 1576–1584 (2017).

236. Xu, X., Hanganu-Opatz, I. L. & Bieler, M. Cross-Talk of Low-Level Sensory and High-Level Cognitive Processing: Development, Mechanisms, and Relevance for Cross-Modal Abilities of the Brain. Frontiers in Neurorobotics 14, (2020).

237. Patel, A. B., Tsilioni, I., Leeman, S. E. & Theoharides, T. C. Neurotensin stimulates sortilin and mTOR in human microglia inhibitable by methoxyluteolin, a potential therapeutic target for autism. Proc Natl Acad Sci U S A 113, E7049–E7058 (2016).

238. Dong, J. et al. The effect of intrauterine inflammation on mTOR signaling in mouse fetal brain. Dev Neurobiol 80, 149–159 (2020).

239. Zhao, X. et al. Noninflammatory Changes of Microglia Are Sufficient to Cause Epilepsy. Cell Rep 22, 2080–2093 (2018).

240. Zimmer, T. S. et al. Tuberous Sclerosis Complex as Disease Model for Investigating mTOR-Related Gliopathy During Epileptogenesis. Front Neurol 11, 1028 (2020).

241. Huang, A. S. et al. Characterizing effects of age, sex and psychosis symptoms on thalamocortical functional connectivity in youth. Neuroimage 243, 118562 (2021).

242. Villasana-Salazar, B. & Vezzani, A. Neuroinflammation microenvironment sharpens seizure circuit. Neurobiol Dis 178, 106027 (2023).

243. Jiang, P. et al. Icariin alleviates autistic-like behavior, hippocampal inflammation and vGlut1 expression in adult BTBR mice. Behav Brain Res 445, 114384 (2023).

244. O’Donnell, C. Nonlinear slow-timescale mechanisms in synaptic plasticity. Current Opinion in Neurobiology 82, 102778 (2023).

245. Gonzalez, K. C., Negrean, A., Liao, Z., Polleux, F. & Losonczy, A. Synaptic Basis of Behavioral Timescale Plasticity. 2023.10.04.560848 Preprint at 10.1101/2023.10.04.560848 (2023).

246. Duarte, R., Seeholzer, A., Zilles, K. & Morrison, A. Synaptic patterning and the timescales of cortical dynamics. Current Opinion in Neurobiology 43, 156–165 (2017).

247. Okur, Z. et al. Control of neuronal excitation-inhibition balance by BMP-SMAD1 signalling. Nature 629, 402–409 (2024).

248. Peddada, S., Yasui, D. H. & LaSalle, J. M. Inhibitors of differentiation (ID1, ID2, ID3 and ID4) genes are neuronal targets of MeCP2 that are elevated in Rett syndrome. Hum Mol Genet 15, 2003–2014 (2006).

249. van Ravenswaaij-Arts, C. M., Hefner, M., Blake, K. & Martin, D. M. CHD7 Disorder. in GeneReviews® (eds Adam, M. P. et al.) (University of Washington, Seattle, Seattle (WA), 1993).

250. Chen, H., Xu, G., Du, H., Yi, M. & Li, C. Integrative analysis of gene expression associated with epilepsy in human epilepsy and animal models. Mol Med Rep 13, 4920–4926 (2016).

251. Lee, T.-S. et al. Gene Expression in Temporal Lobe Epilepsy is Consistent with Increased Release of Glutamate by Astrocytes. Mol Med 13, 1–13 (2007).

252. Ryan, R. et al. Functional characterization of tektin-1 in motile cilia and evidence for TEKT1 as a new candidate gene for motile ciliopathies. Hum Mol Genet 27, 266–282 (2018).

253. Khalil, R. et al. PSMD12 haploinsufficiency in a neurodevelopmental disorder with autistic features. Am J Med Genet B Neuropsychiatr Genet 177, 736–745 (2018).

254. Montgomery, S. H. & Mundy, N. I. Evolution of ASPM is associated with both increases and decreases in brain size in primates. Evolution 66, 927–932 (2012).

255. Bishop, D. V. M. Genes, cognition, and communication: insights from neurodevelopmental disorders. Ann N Y Acad Sci 1156, 1–18 (2009).

256. Shen, J. et al. ASPM mutations identified in patients with primary microcephaly and seizures. J Med Genet 42, 725–729 (2005).

257. Eagleson, K. L. et al. Genetic disruption of the autism spectrum disorder risk gene PLAUR induces GABAA receptor subunit changes. Neuroscience 168, 797–810 (2010).

258. Eagleson, K. L., Campbell, D. B., Thompson, B. L., Bergman, M. Y. & Levitt, P. The autism risk genes MET and PLAUR differentially impact cortical development. Autism Res 4, 68–83 (2011).

259. Spinelli, E. et al. Pathogenic MAST3 Variants in the STK Domain Are Associated with Epilepsy. Ann Neurol 90, 274–284 (2021).

260. Xu, Q. et al. MAST3 modulates the inflammatory response and proliferation of fibroblast-like synoviocytes in rheumatoid arthritis. Int Immunopharmacol 77, 105900 (2019).

261. Wang, Y., Li, Y., Wang, G., Lu, J. & Li, Z. Overexpression of Homer1b/c induces valproic acid resistance in epilepsy. CNS Neurosci Ther 29, 331–343 (2023).

262. Phadke, L. et al. A primary rodent triculture model to investigate the role of glia-neuron crosstalk in regulation of neuronal activity. Front Aging Neurosci 14, 1056067 (2022).

263. Du, Y., Brennan, F. H., Popovich, P. G. & Zhou, M. Microglia maintain the normal structure and function of the hippocampal astrocyte network. Glia 70, 1359–1379 (2022).

264. Shan, W. et al. Neuronal PAS domain protein 4 (Npas4) controls neuronal homeostasis in pentylenetetrazole-induced epilepsy through the induction of Homer1a. J Neurochem 145, 19–33 (2018).

265. Gomez Perdiguero, E., et al. Tissue-resident macrophages originate from yolk-sac-derived erythro-myeloid progenitors. Nature 518, 547–551 (2015).

266. Wang, S. et al. IQGAP3, a novel effector of Rac1 and Cdc42, regulates neurite outgrowth. J Cell Sci 120, 567–577 (2007).

267. Reilly, M. L. et al. Loss-of-function mutations in KIF14 cause severe microcephaly and kidney development defects in humans and zebrafish. Hum Mol Genet 28, 778–795 (2019).

268. Zheng, E., Cai, Z., Li, W., Ni, C. & Fang, Q. Achaete-scute complex-like 2 regulated inflammatory mechanism through Toll-like receptor 4 activating in stomach adenocarcinoma. World J Surg Oncol 20, 266 (2022).

269. Tong, D.-L. et al. The critical role of ASD-related gene CNTNAP3 in regulating synaptic development and social behavior in mice. Neurobiol Dis 130, 104486 (2019).

270. Firouzabadi, N. et al. Genetic Variants of Angiotensin-Converting Enzyme Are Linked to Autism: A Case-Control Study. PLoS One 11, e0153667 (2016).

271. Pereira, M. G. A. G. et al. Inhibition of the renin-angiotensin system prevents seizures in a rat model of epilepsy. Clin Sci (Lond) 119, 477–482 (2010).

272. Yu, S., Wang, G., Yao, B., Xiao, L. & Tuo, H. Arc and Homer1 are involved in comorbid epilepsy and depression: A microarray data analysis. Epilepsy Behav 132, 108738 (2022).

273. Tărlungeanu, D. C. et al. Impaired Amino Acid Transport at the Blood Brain Barrier Is a Cause of Autism Spectrum Disorder. Cell 167, 1481–1494.e18 (2016).

274. Fudge, N. J. & Mearow, K. M. Extracellular matrix-associated gene expression in adult sensory neuron populations cultured on a laminin substrate. BMC Neurosci 14, 15 (2013).

275. Chénier, S. et al. CHD2 haploinsufficiency is associated with developmental delay, intellectual disability, epilepsy and neurobehavioural problems. J Neurodev Disord 6, 9 (2014).

276. Cepeda, C. et al. Cellular antiseizure mechanisms of everolimus in pediatric tuberous sclerosis complex, cortical dysplasia, and non-mTOR-mediated etiologies. Epilepsia Open 3, 180–190 (2018).

277. Franz, D. N. et al. Adjunctive everolimus therapy for tuberous sclerosis complex-associated refractory seizures: Results from the postextension phase of EXIST-3. Epilepsia 62, 3029–3041 (2021).

278. Liu, Y. et al. Sensitivity Analysis of the Efficacy of Everolimus for Neurocognitive Symptoms in PTEN Hamartoma Tumor Syndrome. Adv Ther 10.1007/s12325-025-03441-y (2025) doi:10.1007/s12325-025-03441-y.

279. de la Torre-Martinez, R., Ketzef, M. & Silberberg, G. Ongoing movement controls sensory integration in the dorsolateral striatum. Nat Commun 14, 1004 (2023).

280. Alloway, K. D., Smith, J. B., Mowery, T. M. & Watson, G. D. R. Sensory Processing in the Dorsolateral Striatum: The Contribution of Thalamostriatal Pathways. Front. Syst. Neurosci. 11, (2017).

281. Tomioka, R. et al. The External Globus Pallidus as the Hub of the Auditory Cortico-Basal Ganglia Loop. eNeuro 11, (2024).

282. Johansson, Y. & Ketzef, M. Sensory processing in external globus pallidus neurons. Cell Rep 42, 111952 (2023).

283. Boyd, B. A. et al. Sensory features and repetitive behaviors in children with autism and developmental delays. Autism Res 3, 78–87 (2010).

284. Joosten, A. V. & Bundy, A. C. Sensory processing and stereotypical and repetitive behaviour in children with autism and intellectual disability. Aust Occup Ther J 57, 366–372 (2010).

285. Riby, D. M., Janes, E. & Rodgers, J. Brief report: exploring the relationship between sensory processing and repetitive behaviours in Williams Syndrome. J Autism Dev Disord 43, 478–482 (2013).

286. Fetta, A. et al. Relationship between Sensory Alterations and Repetitive Behaviours in Children with Autism Spectrum Disorders: A Parents’ Questionnaire Based Study. Brain Sciences 11, 484 (2021).

287. Williams, K. L., Campi, E. & Baranek, G. T. Associations among sensory hyperresponsiveness, restricted and repetitive behaviors, and anxiety in autism: An integrated systematic review. Research in Autism Spectrum Disorders 83, 101763 (2021).

288. Waizbard-Bartov, E. et al. Changes in the severity of autism symptom domains are related to mental health challenges during middle childhood. Autism 28, 1216–1230 (2024).

289. Nwaordu, G. & Charlton, R. A. Repetitive Behaviours in Autistic and Non-Autistic Adults: Associations with Sensory Sensitivity and Impact on Self-Efficacy. J Autism Dev Disord 10.1007/s10803-023-06133-0 (2023) doi:10.1007/s10803-023-06133-0.

290. Pagani, M. et al. Biological subtyping of autism via cross-species fMRI. 2025.03.04.641400 Preprint at 10.1101/2025.03.04.641400 (2025).

291. Rudie, J. D. et al. Reduced functional integration and segregation of distributed neural systems underlying social and emotional information processing in autism spectrum disorders. Cereb Cortex 22, 1025–1037 (2012).

292. Sigar, P., Uddin, L. Q. & Roy, D. Altered global modular organization of intrinsic functional connectivity in autism arises from atypical node-level processing. Autism Res 16, 66–83 (2023).

293. You, X. et al. Atypical modulation of distant functional connectivity by cognitive state in children with Autism Spectrum Disorders. Front Hum Neurosci 7, 482 (2013).

294. Harlalka, V., Bapi, R. S., Vinod, P. K. & Roy, D. Age, Disease, and Their Interaction Effects on Intrinsic Connectivity of Children and Adolescents in Autism Spectrum Disorder Using Functional Connectomics. Brain Connect 8, 407–419 (2018).

295. Henry, T. R., Dichter, G. S. & Gates, K. Age and Gender Effects on Intrinsic Connectivity in Autism Using Functional Integration and Segregation. Biol Psychiatry Cogn Neurosci Neuroimaging 3, 414–422 (2018).

296. Keown, C. L. et al. Network organization is globally atypical in autism: A graph theory study of intrinsic functional connectivity. Biol Psychiatry Cogn Neurosci Neuroimaging 2, 66–75 (2017).

297. Passamonti, L. et al. Neuroinflammation and Functional Connectivity in Alzheimer’s Disease: Interactive Influences on Cognitive Performance. J Neurosci 39, 7218–7226 (2019).

298. Colasanti, A. et al. Hippocampal Neuroinflammation, Functional Connectivity, and Depressive Symptoms in Multiple Sclerosis. Biol Psychiatry 80, 62–72 (2016).

299. Marsland, A. L. et al. Systemic inflammation and resting state connectivity of the default mode network. Brain Behav Immun 62, 162–170 (2017).

300. Alvarez, G. M., Hackman, D. A., Miller, A. B. & Muscatell, K. A. Systemic inflammation is associated with differential neural reactivity and connectivity to affective images. Social Cognitive and Affective Neuroscience 15, 1024–1033 (2020).

301. Wang, H. et al. Functional brain rewiring and altered cortical stability in ulcerative colitis. Mol Psychiatry 27, 1792–1804 (2022).

302. Wei, Y. et al. Applying dimensional psychopathology: transdiagnostic prediction of executive cognition using brain connectivity and inflammatory biomarkers. Psychological Medicine 53, 3557–3567 (2023).

303. Liu, Y. et al. Altered functional connectivity and topology structures in default mode network induced by inflammatory exposure in aged rat: A resting-state functional magnetic resonance imaging study. Frontiers in Aging Neuroscience 14, (2022).

304. Harutyunyan, A. A., Harutyunyan, H. A. & Yenkoyan, K. B. Novel Probable Glance at Inflammatory Scenario Development in Autistic Pathology. Front Psychiatry 12, 788779 (2021).

305. Pape, K., Tamouza, R., Leboyer, M. & Zipp, F. Immunoneuropsychiatry - novel perspectives on brain disorders. Nat Rev Neurol 15, 317–328 (2019).

306. Miller, A. H., Haroon, E., Raison, C. L. & Felger, J. C. Cytokine Targets in the Brain: Impact on Neurotransmitters and Neurocircuits. Depress Anxiety 30, 297–306 (2013).

307. Breitinger, U. & Breitinger, H.-G. Excitatory and inhibitory neuronal signaling in inflammatory and diabetic neuropathic pain. Mol Med 29, 53 (2023).

308. Eren-Koçak, E. & Dalkara, T. Ion Channel Dysfunction and Neuroinflammation in Migraine and Depression. Front. Pharmacol. 12, (2021).

309. Min, S. S. et al. Chronic brain inflammation impairs two forms of long-term potentiation in the rat hippocampal CA1 area. Neurosci Lett 456, 20–24 (2009).

310. Wilson, J. F., Lodhia, V., Courtney, D. P., Kirk, I. J. & Hamm, J. P. Evidence of hyper-plasticity in adults with Autism Spectrum Disorder. Research in Autism Spectrum Disorders 43–44, 40–52 (2017).

311. Rodriguez, J. I. & Kern, J. K. Evidence of microglial activation in autism and its possible role in brain underconnectivity. Neuron Glia Biol 7, 205–213 (2011).

312. Skaper, S. D., Facci, L., Zusso, M. & Giusti, P. An Inflammation-Centric View of Neurological Disease: Beyond the Neuron. Front Cell Neurosci 12, 72 (2018).

313. Deverman, B. E. & Patterson, P. H. Cytokines and CNS development. Neuron 64, 61–78 (2009).

314. Di Marco, B., Bonaccorso, C. M., Aloisi, E., D’Antoni, S. & Catania, M. V. Neuro-Inflammatory Mechanisms in Developmental Disorders Associated with Intellectual Disability and Autism Spectrum Disorder: A Neuro-Immune Perspective. CNS Neurol Disord Drug Targets 15, 448–463 (2016).

315. Wei, H. et al. IL-6 is increased in the cerebellum of autistic brain and alters neural cell adhesion, migration and synaptic formation. Journal of Neuroinflammation 8, 52 (2011).

316. Thion, M. S. et al. Biphasic Impact of Prenatal Inflammation and Macrophage Depletion on the Wiring of Neocortical Inhibitory Circuits. Cell Rep 28, 1119–1126.e4 (2019).

317. Thion, M. S. & Garel, S. On place and time: microglia in embryonic and perinatal brain development. Curr Opin Neurobiol 47, 121–130 (2017).

318. Win-Shwe, T.-T. et al. Social behavior, neuroimmune markers and glutamic acid decarboxylase levels in a rat model of valproic acid-induced autism. J Toxicol Sci 43, 631–643 (2018).

319. Mittli, D. Inflammatory processes in the prefrontal cortex induced by systemic immune challenge: Focusing on neurons. Brain Behav Immun Health 34, 100703 (2023).

320. Filipello, F. et al. The Microglial Innate Immune Receptor TREM2 Is Required for Synapse Elimination and Normal Brain Connectivity. Immunity 48, 979–991.e8 (2018).

321. Hunt, R. F., Boychuk, J. A. & Smith, B. N. Neural circuit mechanisms of post-traumatic epilepsy. Front Cell Neurosci 7, 89 (2013).

